# Trichome and inflorescence evolution in the paleopolyploid tree genus *Greyia Hook. & Harv*

**DOI:** 10.64898/2026.05.28.728437

**Authors:** Xiao Ma, Jiyang Chang, Jana Botes, Yves Van de Peer, Dave Kenneth Berger

## Abstract

The southern African endemic tree genus *Greyia* Hook. & Harv. (Francoaceae, Geraniales) comprises three species distinguished by striking variation in trichome morphology and floral architecture, yet the genomic and regulatory basis of these traits has remained largely unexplored. Here, we present chromosome-scale genome assemblies for all recognized *Greyia* species, namely *Greyia radlkoferi*, *Greyia sutherlandii*, and *Greyia flanaganii*, providing the first high-quality genomic resources for Geraniales. Phylogenomic analyses placed Geraniales as sister to Crossosomatales, with divergence at ∼103 million years ago (Mya), and indicated that *Greyia* diversified recently during the Quaternary (∼1.39 Mya). Comparative genomics revealed a *Greyia*-specific whole-genome triplication event dated to ∼86.9 Mya, generating extensive gene duplication which has been retained. Transcriptomic analyses linked divergence in trichome morphology to regulatory subfunctionalization of the conserved MYB–bHLH–WD40 (MBW) complex, including transcriptional silencing of GL1 and shifts in GL3/EGL3 expression in *G. sutherlandii*. In addition, analyses of flower and inflorescence identified expansion of MADS-box transcription factor families and species-specific cis-regulatory divergence in *LFY* and *FT*. Notably, predicted AGL6 binding sites are retained in the *LFY* and *FT* promoters of *G. radlkoferi* and *G. sutherlandii* but absent in *G. flanaganii*, which aligns with differences in inflorescence density among species. Together, these results demonstrate how polyploidy and regulatory divergence of conserved developmental pathways have shaped morphological diversification in *Greyia*.

**Significance Statement:** Understanding how morphological diversity arises from conserved pathways is a central evolutionary challenge, particularly in lineages shaped by ancient polyploidy. We present the first chromosome-scale genomes for the order Geraniales, focusing on three southern African endemic *Greyia* species. We demonstrate that regulatory divergence following a lineage-specific whole-genome triplication, rather than changes in gene content, drives variation in trichome morphology and floral architecture.

## Introduction

*Greyia* Hook. & Harv. is a small genus of trees/shrubs endemic to South Africa and Eswatini (DAHLGREN and van Wyk, 1988), comprising three currently recognised species: *Greyia sutherlandii* Hook. & Harv., *Greyia radlkoferi* Szyszyl., and *Greyia flanaganii* Bolus (Phillips and Gower, 1923, Killick and Kimpton, 1977, Steyn *et al*., 1999). They are commonly known as bottlebrush or beacon trees on account of the spectacular red inflorescences of *G. sutherlandii* and *G. radlkoferi* that appear in late winter/early spring prior to leaf sprouting in these deciduous species (Figure 1a, b). *Greyia* are prolific nectar producers, thus contributing ecosystem services to a range of generalist bird and insect pollinators at times of nectar scarcity in the mountainous landscapes of the Great Escarpment and Drakensberg of South Africa (DAHLGREN and van Wyk, 1988). *G. radlkoferi* (woolly bottlebrush) is the most northerly species distributed in Limpopo and Mpumalanga provinces (De la Cruz, 2016), with *G. sutherlandii* (glossy bottlebrush) overlapping in distribution in the latter province, but mainly occupying KwaZulu-Natal province (Mbambezeli, 2016). The most southerly species, *G. flanaganii*, is called the Kei bottlebrush on account of its limited distribution along the Kei River and its tributaries in the Eastern Cape province of South Africa (Mbambezeli, 2002). Unlike its deciduous congeneric species, *G. flanaganii* is an evergreen and lower-altitude species, and possesses distinct inflorescences characterized by sparsely arranged, lax, pendulous, urn-shaped red flowers (Figure 1c). Differentiation between *G. radlkoferi* and *G. sutherlandii* in the field is often based on the “woolly” abaxial leaf morphology of *G. radlkoferi*, in contrast to the glabrous (hairless) leaves of *G. sutherlandii*. (Figure 1a, b). However, several authors have noted that this can be an unreliable identification trait (DAHLGREN and van Wyk, 1988).

**Figure 1.**
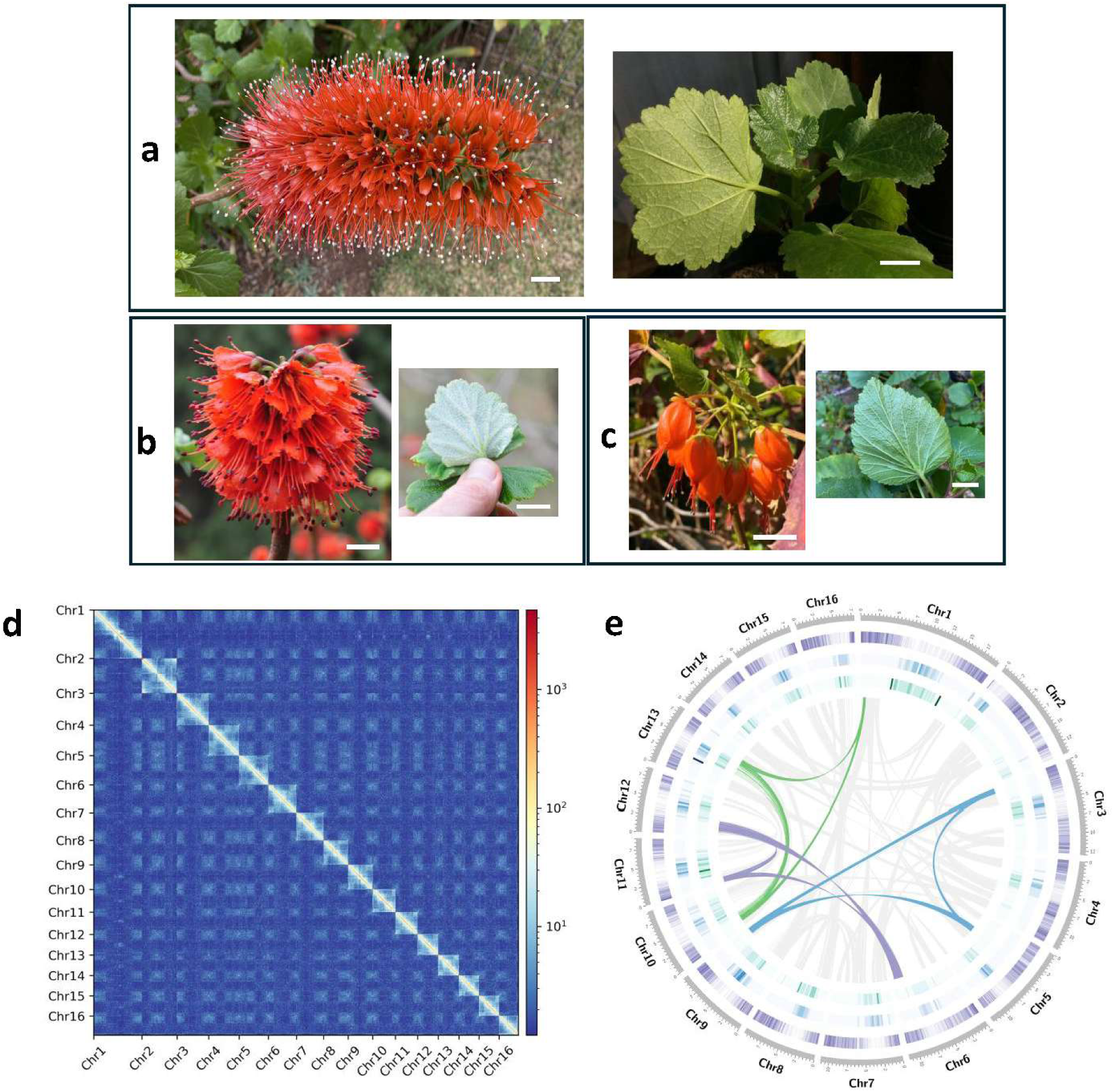
*Greyia* species morphology, genome assembly of *Greyia radlkoferi*. **a-c.** Inflorescence and abaxial leaf morphologies of *Greyia sutherlandii*, *Greyia radlkoferi,* and *Greyia flanaganii*, respectively. Scale bars = 2 cm. **d**. Hi-C interaction heatmap of the assembled *Greyia radlkoferi* genome. **e.** *G. radlkoferi* genome assembly. The tracks from the inside to the outside correspond to Copia elements, Gypsy elements, gene density per 100 kb, and chromosomes. The innermost grey curves represent all syntenic regions, while the three colored lines denote representative syntenic regions retained in triplicate.

Apart from the ecological importance of *Greyia* spp., they have economic value due to their horticultural potential as garden plants, having been popular already in the last century in Europe and South Africa (F, 2010). Extracts from *Greyia* leaves have also attracted interest for medicinal applications, particularly for the treatment of hyperpigmentation, with studies demonstrating anti-tyrosinase activity in *G. radlkoferi* and *G. flanaganii* without detectable cytotoxic or mutagenic effects (Mapunya *et al*., 2011, Lall *et al*., 2016). The active biomarker compound was identified as 2’,4’,6’-trihydroxydihydrochalcone (Lall *et al*., 2016). A public-private partnership is underway in South Africa to develop the technology pipeline from orchard cultivation to herbal remedy formulation for the commercialization of *Greyia* extracts for skin disorders.

Taxonomic classification of the *Greyia* genus based on morphology has long been problematic. Since its initial placement in Saxifragaceae by Harvey (Harvey, 1862), *Greyia* has subsequently been affiliated with Melianthaceae or treated as a separate family (Greyiaceae), reflecting uncertainty in its systematic position (DAHLGREN and van Wyk, 1988). To date, the amount of phylogenetically informative molecular data available for *Greyia* is limited with only plastid gene or ITS data for one or two individuals available on Genbank (Morgan and Soltis, 1993, Linder *et al*., 2006). Recently, DNA barcoding genes (ITS, *matK*, *psbA-trnH*, *trnLF*) were sequenced from geographically separated populations of *Greyia* (five trees per species) (Botha *et al*., 2026). However, phylogenetic analysis indicated there were insufficient polymorphisms to confidently differentiate the species from one another, although *G. flanaganii* was the most distinct. Nevertheless, molecular-based classification of *Greyia* has placed it in the Order Geraniales, which comprises two families, Geraniaceae and Francoaceae with *Greyia* in the latter (Byng *et al*., 2016). There are seven genera in Geraniaceae (*California*, *Erodium*, *Geranium*, *Hypseocharis*, *Monsonia*, *Pelargonium*, *Sarcocaulon*), and seven genera in Francoaceae (*Balbisia*, *Bersama*, *Francoa*, *Greyia*, *Melianthus*, *Rhynchotheca*, *Viviania*). The family name Francoaceae takes precedence over its synonyms Greyiaceae and Melianthaceae (Byng *et al*., 2016). Linder et al. (Linder *et al*., 2006) presented molecular data for members of this family with a study focused on *Melianthus* species with *Greyia* as the outgroup. DNA sequence data for a nuclear (ITS) and plastid (*trnL-trnF*, *psb*A-*trn*H) gene fragments were determined, which confirmed monophyly of *Melianthus* distinct from *Greyia*.

More broadly, the phylogenetic placement of Geraniales has remained unresolved, largely due to the limited availability of genomic data. Phylogenomic trees of angiosperm lineages, such as APG IV, have relied primarily on plastid and rDNA sequences, which placed Geraniales and Myrtales as early-diverging sister group within the Malvids clade (Byng *et al*., 2016). Another phylogenetic analysis based on 4,792 chloroplast genomes, representing all currently recognized families, also supported Geraniales and Myrtales as the sister group to the remaining Malvids (Li *et al*., 2021). However, recent nuclear phylogenomic studies using 353 protein-coding genes of angiosperms (Angiosperms353) have recovered a different topology, supporting a sister relationship between Geraniales and Crossosomatales, with Myrtales grouping instead with Zygophyllales (Zuntini *et al*., 2024). These four orders formed a clade that is sister to the rosids, but not to the Malvid clade. Although plastid– and nuclear-based phylogenies are largely congruent for most angiosperm orders, Geraniales represents a notable exception. Importantly, *Greyia* species are poorly represented or entirely absent from these large-scale phylogenomic datasets, including the Angiosperms353 framework. Genome-scale data from *Greyia* therefore provides a critical opportunity to clarify the phylogenetic position of Geraniales and to establish a much-needed genomic reference for this understudied order.

The remarkable evolutionary success and diversification of angiosperms are intrinsically linked to genomic plasticity, with whole-genome duplication (WGD) serving as a pivotal driver of morphological and ecological innovation (Cai *et al*., 2019, Walden *et al*., 2020). Following WGD, most genomes experience a phase of reorganization, including the loss, pseudogenization, subfunctionalization and neofunctionalization of coding sequences (Heslop-Harrison *et al*., 2023). These processes create opportunities for functional diversification (Heslop-Harrison *et al*., 2023) and adaptive radiation (Van de Peer *et al*., 2021). *Greyia* exhibits pronounced variation in trichome morphology and inflorescence architecture (Figure 1a–c), making it an attractive system to investigate if and how WGD-derived processes contribute to morphological diversification within a restricted lineage. For example, floral ontology and development in the Geraniales order has been the subject to several morphological studies (Ronse Decraene *et al*., 2001, Jeiter *et al*., 2017a) including a comparison between *Greyia* spp. and *Francoa* (Ronse Decraene and Smets, 1999). Overall flower morphologies in the order are distinct since genera such as *Melianthus* and *Pelargonium* have zygomorphic flowers, in contrast to others such as *Greyia* and *Geranium* with actinomorphic flowers (Steyn *et al*., 1987, Jeiter *et al*., 2017b). However, the absence of genome-scale resources in the Geraniales has so far limited exploration of the molecular basis of these morphological adaptations.

Here, we present high-quality, chromosome-scale genome assemblies and annotations for all three recognized *Greyia* species, representing the first genomic resources for the order Geraniales. Using comparative genomics and transcriptomic analyses, we reconstruct the evolutionary history of *Greyia*, identify a lineage-specific whole-genome triplication event, and examine how gene retention, expression divergence, and cis-regulatory evolution have shaped trichome differentiation and inflorescence architecture. Together, these data provide new insights into the genomic mechanisms underlying morphological diversification in *Greyia* and establish a foundation for future evolutionary and functional studies within Geraniales.

## Results

### Assemblies and annotations of three *Greyia* genomes

Prior to *de novo* assembly, genome sizes were estimated by k-mer analysis of Illumina short reads, indicating genome sizes of 178–192 Mb for *Greyia* (2n = 2x = 32; Figure S1). Oxford Nanopore long-read sequencing generated 16.1 Gb (102×), 13.4 Gb (75×), and 15.0 Gb (84×) of data for *G. radlkoferi*, *G. sutherlandii*, and *G. flanaganii*, respectively. De novo assembly using Flye v2.9.3 (Kolmogorov *et al*., 2019) yielded contig-level assemblies of comparable size (175–178 Mb), consistent with k-mer estimates. The *G. radlkoferi* assembly comprised 314 contigs (N50 = 5.9 Mb), *G. sutherlandii* 394 contigs (N50 = 2.5 Mb), and *G. flanaganii* 261 contigs (N50 = 5.7 Mb) (Table 1). To generate chromosome-scale assemblies, 27.5 Gb (152×) of Hi-C data were produced for *G. radlkoferi*, enabling scaffolding into 16 pseudomolecules with ∼5 Mb remaining unplaced (Figure 1d). Using *G. radlkoferi* as a reference, genome-guided scaffolding was applied to *G. sutherlandii* and *G. flanaganii* (Figure S2). All three assemblies were resolved into 16 pseudomolecules, anchoring 97.7–98.9% of contigs (173–174 Mb) and achieving scaffold N50 values of 9.9–11.2 Mb (Table 1). Assembly quality was further validated by mapping RNA-seq reads from multiple tissues, with 96.4 – 98.6% alignment (Table S1). Benchmarking Universal Single-Copy Orthologs (BUSCO) analysis using the eudicots_odb10 dataset confirmed high completeness (98.5–98.6%, Table 1). Together, these results demonstrate the high quality of the three *Greyia* genome assemblies.

**Table 1.** Statistics of the three *Greyia* genome assemblies and annotations.

| Assembly features |  |  |  |
| --- | --- | --- | --- |
| Species | *G. radlkoferi* | *G. sutherlandii* | *G. flanaganii* |
| Estimated genome size (Mb)* | 178 | / | 192 |
| Number of contigs | 314 | 394 | 261 |
| N50 contig length (Mb) | 5.9 | 2.5 | 4.7 |
| Total size of contigs (Mb) | 175 | 177 | 178 |
| Number of pseudo-chromosomes | 16 | 16 | 16 |
| Total size of pseudo-chromosomes (Mb) | 173 | 173 | 174 |
| N50 scaffold length (Mb) | 11.2 | 9.9 | 9.9 |
| BUSCO (complete) | 98.6% | 98.6% | 98.5% |
| TE annotation features |  |  |  |
| Repeat content | 27.0% | 28.5% | 29.1% |
| LTR elements | 11.8 % | 12.2 % | 11.6 % |
| Copia | 4.01 % | 4.25 % | 3.89 % |
| Gypsy | 5.3 % | 5.1 % | 4.7 % |
| DNA transposons | 1.6 % | 1.5 % | 2.1 % |
| Gene annotation features |  |  |  |
| Number of gene models | 27,222 | 26,878 | 27,861 |
| Mean gene length (bp) | 2848 | 2883 | 2832 |
| Mean coding sequence length (bp) | 1231 | 1258 | 1221 |
| Mean exons number per gene | 5.39 | 5.43 | 5.36 |
| Mean exon length (bp) | 228 | 231 | 228 |
| Mean intron length (bp) | 369 | 368 | 370 |
| BUSCO (complete) | 99.6% | 99.6% | 99.3% |
| Complete and single-copy | 94.2% | 93.8% | 93.5% |
| Complete and duplicated | 5.3% | 5.8% | 5.8% |
| Fragmented | 0.1% | 0.1% | 0.4% |
| Missing | 0.3% | 0.3% | 0.3% |
| OMArk (complete) | 94.1% | 94.1% | 94.3% |

\* Genome size was estimated using K-mer analysis of Illumina reads (Figure S1).

Repetitive elements account for 27.0%, 28.5%, and 29.1% of the *G. radlkoferi*, *G. sutherlandii*, and *G. flanaganii* genomes, respectively. Long terminal repeat retrotransposons (LTR-RTs) dominate as the major transposable element class, accounting for ∼12% of each genome, with Gypsy (∼5%) and Copia (∼4%) being the most abundant LTR families (Table 1). As observed in other plant genomes, transposable element density is inversely correlated with gene density across the chromosomes (Figure 1e). Following transposable element prediction and masking, protein-coding genes were annotated through an integrated approach combining evidence-based and *ab initio* methods (see Methods). The high gene density in these compact genomes led to the frequent merging of adjacent genes in initial computational annotations. We markedly improved genome annotation quality through extensive manual curation (see Methods), with BUSCO completeness increasing from 90% to >99%. The final annotations comprise 27,222 genes in *G. radlkoferi*, 26,878 in *G. sutherlandii*, and 27,861 in *G. flanaganii* with transcriptome support (TPM >1) for 86.7%, 82.5%, and 80.2% of predicted genes, respectively. BUSCO (v5.7.1) confirmed these as near-complete genome annotations (99.3–99.6% complete; missing; eudicots_odb10), with comparable gene structures among species (Table 1; Figure 1f). Independent evaluation with OMArk (v0.3.1) yielded slightly different but consistent quality metrics (Table 1). Functional annotation using eggNOG-mapper and InterProScan assigned putative roles to 93.1-94.2% of all genes.

### Phylogenomics of Geraniales

Previous phylogenetic frameworks have placed Geraniales either as sister to Myrtales based on plastid data or as sister to Crossosomatales based on nuclear gene analyses (Byng *et al*., 2016, Li *et al*., 2021, Zuntini *et al*., 2024), highlighting uncertainty in the placement of this order within rosids. To clarify the phylogenetic position of Geraniales, we conducted a genome-based phylogenomic analysis using *Greyia* as a representative. We analyzed 17 genomes spanning 14 orders within the superrosids – Brassicales, Malvales, Sapindales, Oxalidales, Malpighiales, Celastrales, Fagales, Rosales, Cucurbitales, Fabales, Geraniales, Crossosomatales, Zygophyllales, and Myrtales – along with two monocot outgroups. This analysis assigned 520,016 genes (92.4% of 563,009 total) to 43,164 orthogroups, including 401 highly conserved single-copy genes used for phylogenetic reconstruction. Among the three *Greyia* species analyzed, we identified 17,048 conserved orthogroups (89.5%) present in all three species, 1,488 orthogroups (7.8%) shared by two species, and 519 species-specific orthogroups (2.7%), indicating a high degree of genomic conservation.

Phylogenomic reconstruction based on the 401 single-copy genes recovered a strongly supported sister relationship between *Greyia* and *Euscaphis japonica* (Crossosomatales), with an estimated divergence time of ∼106 Mya (96.05-110.43 Mya; Figure 2a). This topology places the Geraniales–Crossosomatales clade as sister to the Fabids and Malvids, whereas Myrtales occupies an early-diverging position among the analyzed clades, a relationship that is consistent with phylogenomic inferences from the *E. japonica* genome analysis (Sun *et al*., 2021) and with recent studies proposing a Crossosomatales-Geraniales sister relationship (Zuntini *et al*., 2024). Comparison of our 401-gene dataset with the Angiosperms353 reference set (Johnson *et al*., 2019) revealed that only 52 of these orthogroups show clear homologs with Angiosperms353 loci, indicating that the majority of loci used here are distinct from the universal angiosperm marker set. Consistent with this difference in gene sampling, our topology contrasts with large-scale angiosperm phylogenomic analyses based on Angiosperms353 nuclear genes, which recovered Crossosomatales–Geraniales as early-diverging, with Myrtales and Zygophyllales forming a collective sister group to Fabids and Malvids (Figure 2b). This discrepancy indicates how different datasets can influence the inferred deep evolutionary relationships among superrosids. Within *Greyia*, divergence time estimates indicate that *G. flanaganii* represents the earliest split, occurring approximately 1.39 Mya (0.85-2.03 Mya), followed by the split between *G. radlkoferi* and *G. sutherlandii* at 0.64 Mya (0.33-1.01 Mya) (Figure 2a). These results suggest that diversification within the genus *Greyia* took place during the Quaternary period.

**Figure 2.**
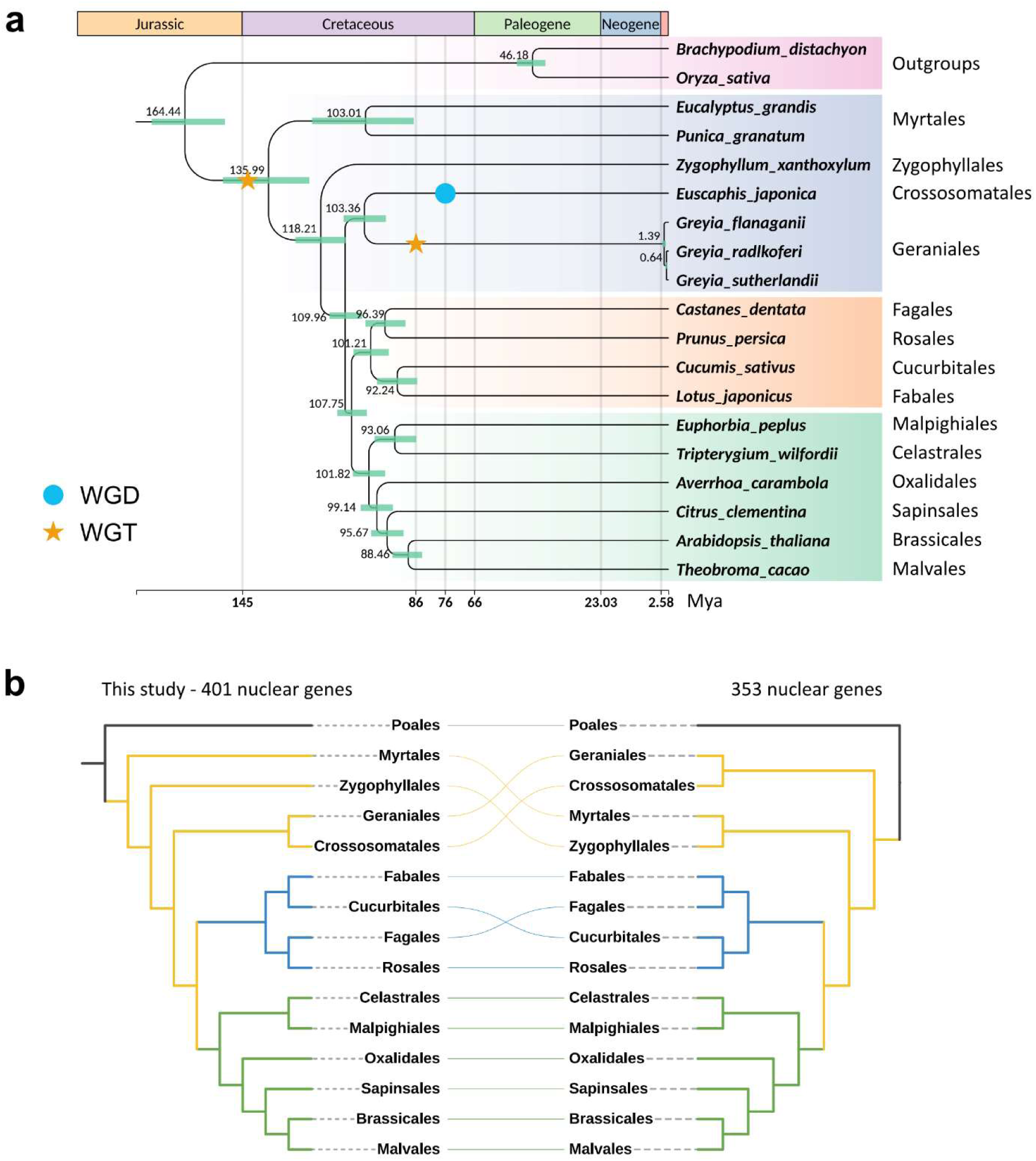
Time-calibrated phylogeny of *Greyia* and corresponding tanglegram showing topological discrepancies between the current study (left) and a schematic tree based on 353 nuclear genes (Angisoperm353 dataset) published previously (right) (Zuntini *et al*., 2024). **a**. Time-calibrated tree using 401 single-copy genes based on genomic data. Only the WGD and WGT events for the groups under study are indicated; **b.** Tanglegram at the ordinal level between the tree of the current study(left) and the schematic tree based on 353 nuclear genes. In the tanglegram, identical orders are connected across the two trees, enabling visual assessment of topological congruence and incongruence.

To further resolve relationships within Geraniales, we incorporated transcriptomic data from six species with BUSCO completeness greater than 65% (Table S1). This analysis recovered two well-supported clades corresponding to Geraniaceae and Francoaceae. The Geraniaceae clade included *Pelargonium dichondrifolium*, *Monsonia marlothii*, *California macrophylla*, and *Erodium chrysanthum* whereas the Francoaceae clade comprised *Greyia*, *Melianthus villosus*, and *Francoa sonchifolia* (Figure S3). In Francoaceae, despite their geographic separation, *F. sonchifolia* from South America shows the closest affinity to the South African genus *Greyia*, with *M. villosus* forming a sister group to the *Francoa*-*Greyia* lineage (Figure S3). Our findings are consistent with previous phylogenetic analyses based on ITS and trnL-F sequences (Palazzesi *et al*., 2012, Sytsma *et al*., 2014), providing additional support for the robustness and stability of evolutionary relationships among genera within Geraniales.

### A *Greyia*-specific whole-genome triplication

To reconstruct the polyploidization history of *Greyia*, we applied multi-method approaches (see Methods). Analysis of synonymous substitution rates (*K_S_*) using syntenic paralogs revealed two clear peaks (∼0.52 and ∼1.34, Figure 3a) in all *Greyia* species, indicative of two polyploidization events. The sister lineage *E. japonica* similarly exhibited two *K_S_* peaks (∼0.34 and ∼1.21), consistent with previously reported whole-genome duplication (WGD) event (Sun *et al*., 2021). The older *K_S_* peaks in *Greyia* (∼1.34) and *E. japonica* (∼1.21) exceed the orthologous divergence with *Vitis vinifera* (∼0.89) (Figure 3a), indicating that they correspond to the shared Gamma event in core eudicots (Chanderbali *et al*., 2022). To determine whether the younger polyploidization events were shared, we compared orthologous *K_S_* values between *Greyia* and *E. japonica*. The ortholog *K_S_* peak between *G. radlkoferi* and *E. japonica* (∼0.85) is higher than the lineage-specific younger peaks (∼0.52 in *Greyia* and ∼0.34 in *E. japonica*), indicating that these events occurred independently after lineage divergence (Figure 3a). Consistent with this inference, synteny analyses revealed a 3:1 relationship between *Greyia* and *V. vinifera*, and a 3:2 relationship between *Greyia* and *E. japonica* (Figure 3c), supporting a *Greyia*-specific whole-genome triplication (WGT) and a *E. japonica*-specific WGD (Sun *et al*., 2021). This WGT event was also supported by intra-genomic collinearity analysis (Figure 1e). Absolute dating (see Methods) estimates the *Greyia* WGT event at ∼86.97 (59.86-102.76) Mya (Figure 3b) and the *E. japonica* WGD at ∼76.18 (48.99-99.66) Mya (Figure S4).

**Figure 3.**
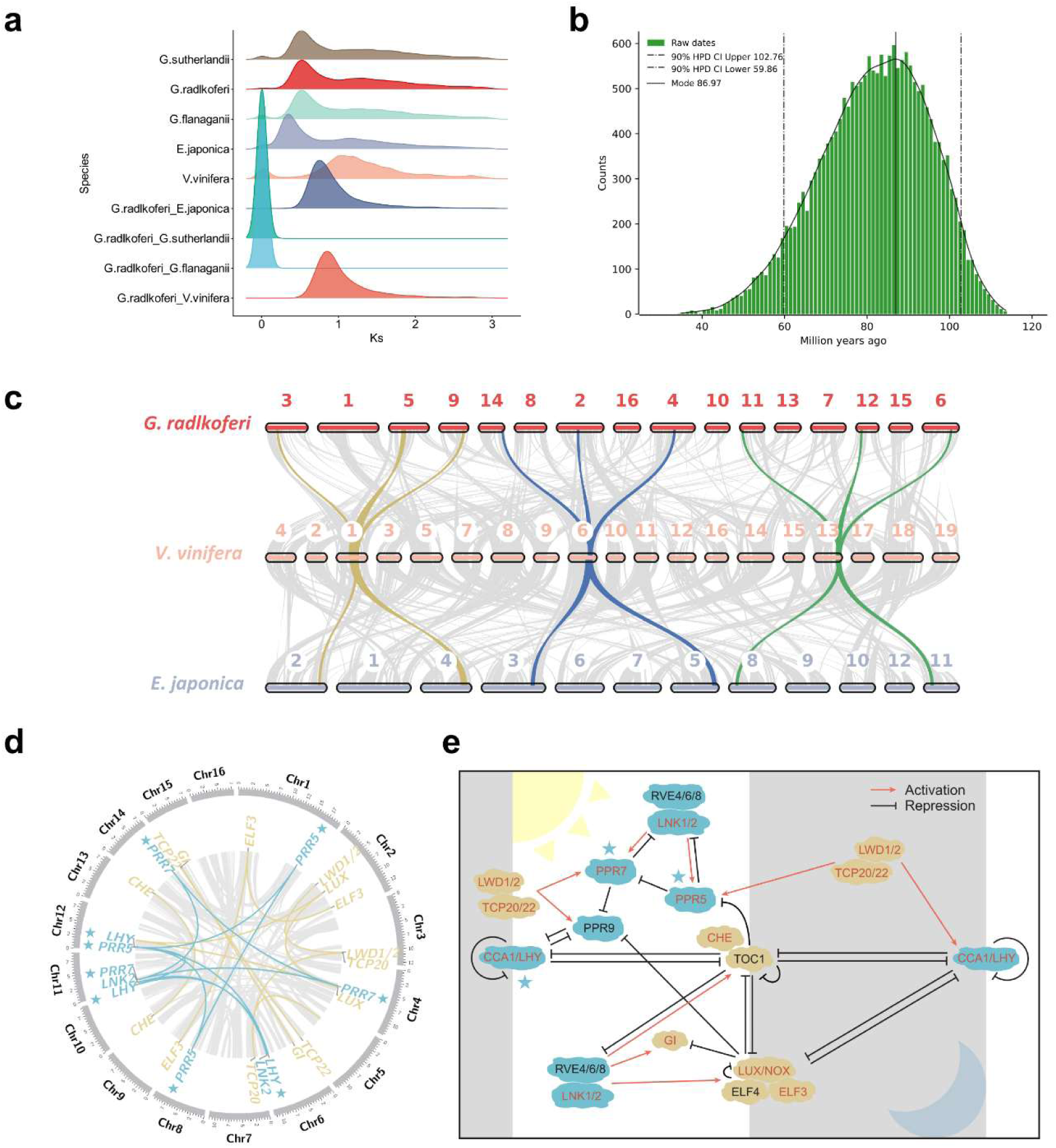
Paleohexaploidy in *Greyia* species. **a.** *K_S_* distributions for anchor pairs (retained paralogs from the WGT and WGD events) of *G. radlkoferi*, *G. sutherlandii*, *G. flanaganii*, and *E. japonica* genomes, and for orthologs of *Greyia* and related species. **b.** Absolute dating (see Methods) of the *G. radlkoferi* WGT event. The age distribution was obtained by phylogenomic dating of *G. radlkoferi* paralogs. The solid black line represents the Kernel Density Estimation (KDE) of the dated paralogs, while the vertical solid black line represents its peak at 86.97 MYA, which was used as the consensus WGD age estimate. The vertical black dotted lines represent the corresponding 90% confidence interval (CI) for the WGD age estimate, 59.86–102.76 MYA. The green histogram shows the raw distribution of dated paralogs. **c.** Synteny between *E. japonica*, *V. vinifera*, and *G. radlkoferi*. The yellow, blue, and green lines highlight the 3:1:2 syntenic relationship among these three species. **d.** Genes related to the photoperiod transduction pathway adapted from Li et al., 2025 (Li *et al*., 2025) are shown in their syntenic relationships derived from the WGT event. Blue lines represent transcription factors; yellow lines represent clock proteins. The blue star represents the genes present as triplicates. Duplicated genes include the transcriptional repressor *CCA1 HIKING EXPEDITION (CHE)*, clock-associated factors *LIGHT-INDUCIBLE AND CLOCK-REGULATED GENE 2 (LNK2)*, *LIGHT-REGULATED WD 1 (LWD1)*, and *TCP-DOMAIN FAMILY PROTEIN 20/22* (*TCP20/22*), the evening complex gene *LUX*, and *GIGANTEA (GI)*; Triplicated genes include *PSEUDO-RESPONSE REGULATOR 5 (PRR5)*, *PSEUDO-RESPONSE REGULATOR 7 (PRR7)*, *CIRCADIAN CLOCK ASSOCIATED 1/LATE ELONGATED HYPOCOTYL 1 (CCA1/LHY1)*, a blue light photoreceptor *CRYPTOCHROME 1 (CRY1)* and a repressor of flowering *EARLY BOLTING IN SHORT DAYS (EBS).* **e.** The core photoperiod transduction pathway adapted from Li et al., 2025 (Li *et al*., 2025). Genes highlighted in red are retained from the WGT event. The blue stars indicate genes retained as triplicates, while the remaining red-highlighted genes are retained as duplicates.

Macrosynteny analysis revealed extensive structural conservation among the three *Greyia* genomes with few large-scale rearrangements (Figure S5). Approximately 25,000 collinear genes were identified in each pairwise comparison and only three inversions were detected (Figure S6). Intragenomic synteny analyses further showed that ∼40% of genes (∼11,000 genes) are retained within ∼500 well-conserved duplicated blocks (Table S3). Among these retained duplicated genes (RDGs), approximately 52% derive from the *Greyia*-specific WGT, representing ∼3,000 gene families per species and a conserved core of 2,518 families shared by all three species (Table S3 and Figure S7). These shared RDGs are significantly enriched (p < 0.05) for functions related to organophosphate and thioester biosynthesis, signal transduction, stomatal movement, and pyruvate metabolism (Figure S7).

In addition to duplicated regions, several genomic regions remain preserved in three copies, notably in Chromosome 1 (syntenic with Chr 10/13), Chromosome 3 (Chr 5/9), and Chromosome 6 (Chr 11/12) (Figure 1). These regions comprise 1,080 genes (360 retained triplicated genes, RTGs) conserved across all three species. RTGs are significantly enriched (p < 0.05) for functions associated with protein metabolism regulation, circadian activity, rhythmic processes, and long-day photoperiodism (Figure S8). Nearly all copies of the core components involved in the photoperiod pathway (Li *et al*., 2025) were located in the syntenic blocks retained following the WGT (Figure 3d, 3e). Notably, the core circadian clock transcription factors (*PRR5*, *PRR7*, *CCA1/LHY1*, *CRY1*, and *EBS*) are retained as triplicates, whereas several clock-associated proteins present as duplicated copies (Figure 3d, 3e). Expression profiling in *G. radlkoferi* revealed that most RDGs and RTGs display tissue-biased expression patterns, whereas a subset of genes, including multiple *PRR7* and *GI* copies, show broad expression across flowers, leaves, and xylem (Figure S9).

### Trichome development

Our current knowledge on trichome development is mostly derived from studies of *Arabidopsis thaliana*, which possesses non-glandular, unicellular trichomes. The C2H2 zinc-finger transcription factor *GLABROUS INFLORESCENCE STEMS* (*GIS*) and its subfamily members act upstream of the MYB-bHLH-WD40 (MBW) transcriptional complex – comprising *GLABRA 1* (*GL1*), *GLABRA 3* (*GL3*)/*ENHANCER OF GLABRA 3* (*EGL3*), and *TRANSPARENT TESTA GLABRA 1* (*TTG1*) – to regulate trichome initiation (Oppenheimer *et al*., 1991, Walker *et al*., 1999, Zhang *et al*., 2003, Grebe, 2012). However, multicellular trichomes in plant species such as tobacco exhibit distinct regulatory mechanisms, and for instance, *GL1* orthologs do not appear to regulate trichome development (Payne *et al*., 1999).

In *Greyia*, leaf surface morphology differs strikingly between species, particularly between *G. radlkoferi* and *G. sutherlandii* (Figure 1 and Figure 4a, 4b). Owing to the lack of a complete RNA-seq dataset for *G. flanaganii*, which lacks xylem tissue (Table S1), our comparative analysis focused primarily on *G. radlkoferi* and *G. sutherlandii*. High-resolution anatomical scans showed that *G. radlkoferi* leaves are densely covered with branched trichomes, whereas *G. sutherlandii* leaves are nearly glabrous (Figure 4a and 4b). These observations are consistent with previous reports describing uniseriate branched hairs in *G. radlkoferi* and predominantly glands in mature *G. sutherlandii* leaves (Steyn, 1974).

**Figure 4.**
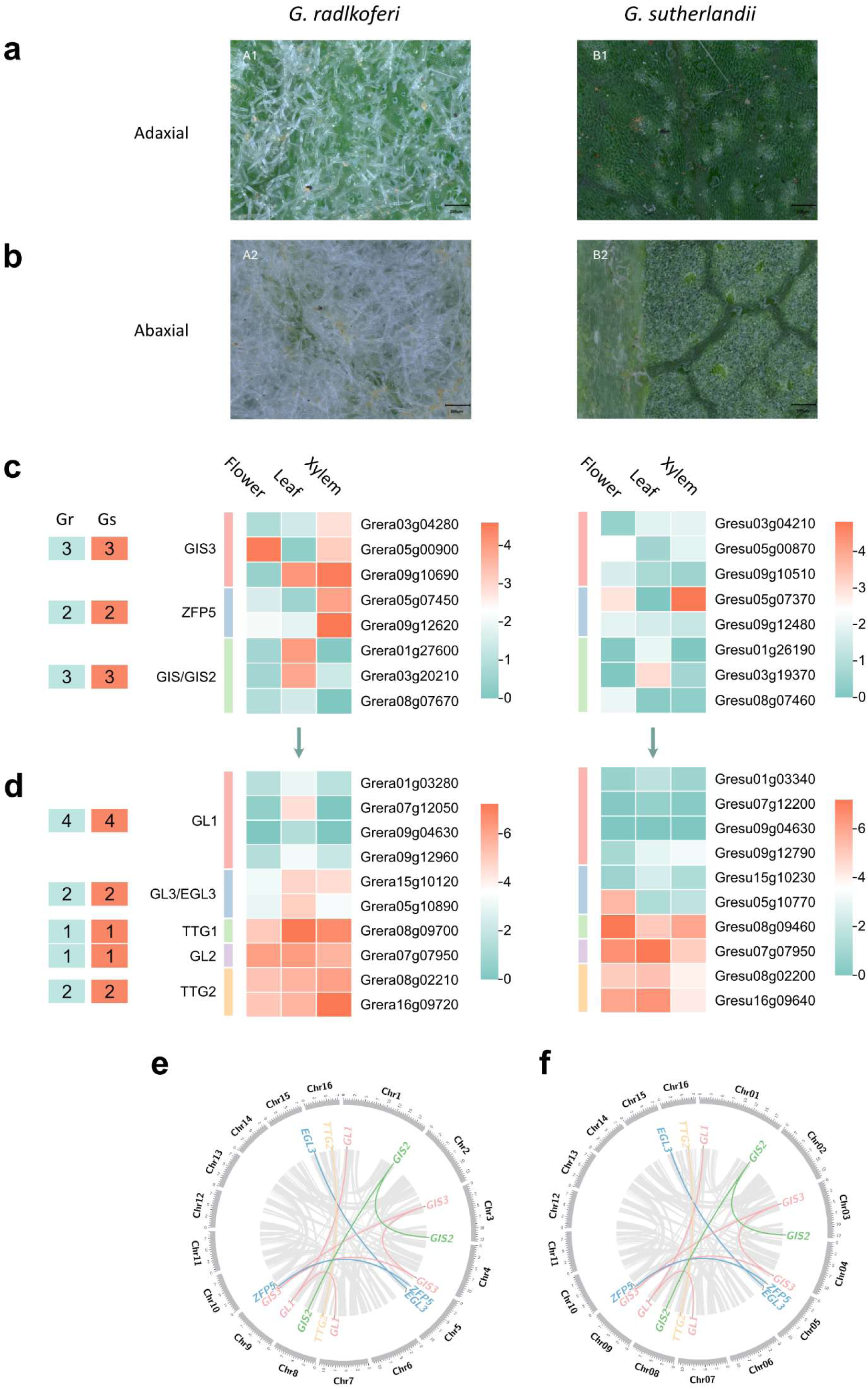
The MBW complex in *Greyia*. **a.** High-resolution anatomical scans of the adaxial (top) leaf surface at 200 µm resolution in *G. radlkoferi* (left) and *G. sutherlandii* (right). **b.** High-resolution anatomical scans of the abaxial (bottom) leaf surface at 200 µm resolution in *G. radlkoferi* (left) and *G. sutherlandii* (right). **c.** Gene copy number and expression patterns of upstream MBW regulators (GIS3, ZFP5, GIS, GIS2) in *G. radlkoferi* (Gr) and *G. sutherlandii* (Gs). Expression values are scaled by log2(TPM +1). TPM is transcripts per million. In the expression profile, the prefix of Grera represents genes in *G. radlkoferi*; Gresu represents genes in *G. sutherlandii*; **d.** Gene copy number and expression patterns of the core MBW complex (*GL1-GL3/EGL3-TTG1*) and downstream targets (*GL2*, *TTG2*). **e.** Genes related to trichome development are shown in their syntenic relationships derived from the WGT event in G. *radlkoferi*. Lines with different colors represent different gene families. **f.** Genes related to trichome development are shown in their syntenic relationships derived from the WGT event in G. *sutherlandii*. Lines with different colors represent different gene families.

To dissect the genetic basis of their divergent trichome phenotypes and to identify candidate regulators, we first identified 311 and 304 MYB transcription factors (TFs) and 39 and 38 bHLH TFs in *G. radlkoferi* and *G. sutherlandii*, respectively. We then screened for these TFs with higher expression in leaves relative to flower and xylem. Comparative transcriptomic analysis highlighted 13 MYB genes with strong leaf-specific expression in *G. radlkoferi*, including a homolog of *GL1* (Figure S10). Although the *GL1* orthogroup is represented by four copies in *Greyia*, only a single copy shows high expression in *G. radlkoferi* leaves (Figure 4d). In contrast, the corresponding *GL1* homolog in *G. sutherlandii* exhibited no detectable expression in any sampled tissue (Figure 4d), paralleling the absence of non-glandular trichomes in this species (Figure 4d). Analysis of the bHLH family further identified two *GL3/EGL3* homologs in *Greyia.* In *G. radlkoferi*, both copies are highly expressed in leaves, whereas in *G. sutherlandii*, one copy showed low expression across all sampled tissues and the other was predominantly expressed in flowers (Figure 4d), indicating marked tissue-specific divergence. We next examined the broader MBW regulatory network, including upstream regulators (*GIS3*, *ZFP5*, *GIS*, *GIS2*) and downstream targets (*GL2*, *TTG2*). All three *Greyia* species retain identical gene complements for these components (Figure 4c, 4d), indicating structural conservation. However, expression patterns differed substantially. Notably, one copy of *GIS3* showed high expression in *G. radlkoferi* leaves but low expression in *G. sutherlandii* leaves (Figure 4c), closely mirroring the expression pattern of *GL1*. Reduced expression of *GIS/GIS2* was also observed in *G. sutherlandii* leaves (Figure 4c). These observations suggest that although the MBW complex and its regulatory genes are structurally conserved in *Greyia*, divergence in their expression likely underlies the species-specific variation in trichome development.

To explore potential regulatory mechanisms underlying these expression differences, we analyzed DNA methylation profiles from ONT reads. In *G. sutherlandii*, CpG methylation was detected in promoter regions upstream of *GL1* and *GIS3*, whereas no such methylation was detected in the corresponding promoters in *G. radlkoferi* (Table S4 and Table S5). These methylation patterns are consistent with the observed transcriptional repression of trichome-related genes in *G. sutherlandii* and with previously reported variation in leaf hairiness within this species.

Finally, key trichome regulators – including *GL1*, *EGL3*, *GIS3*, and *GIS/GIS2* – are present as multi-copy genes derived from the *Greyia*-specific WGT event in both *Greyia* species (Figure 4e and f), demonstrating again how polyploidization provides genetic raw material for potential diversification. Furthermore, expression divergence among these paralogs reveals clear subfunctionalization patterns post-WGT, with different copies preferentially expressed in distinct tissues (e.g., floral vs. vegetative) or developmental stages, suggesting subfunctionalization has contributed to phenotypic innovation in *Greyia*.

### Flower development

Greyia species produce conspicuous red flowers that differ markedly in morphology at anthesis (Figure 1a, b vs. 1c). *G. sutherlandii* and *G. radlkoferi* display open, valvate corollas, whereas *G. flanaganii* forms urceolate (urn-shaped), imbricate flowers with overlapping petals (Figure 1c vs 1a, b). In all three species, flowers bear exserted stamens with red anthers prior to opening (Figure 1b). Floral organ development is described here, whereas inflorescence morphology (Figure 6A) will be discussed in the section “inflorescence development”.

The development of floral structures in most Angiosperm plant species is regulated by MADS-box transcription factors (Soltis *et al*., 2007, O’Maoileidigh *et al*., 2014). Among them, type II MADS-box genes are well-known for their conserved functions in floral organ specification and distinct expression patterns (Becker and Theissen, 2003, Smaczniak *et al*., 2012). Genome-wide analysis identified 87, 89, and 88 putative MADS-box genes in *G. radlkoferi*, *G. sutherlandii*, and *G. flanaganii*, respectively. All three species shared an identical complement of 47 type II MADS-box genes (Table S6) – an expansion compared to *A. thaliana*, including homologous genes for the ABCE model of floral organ identities: five *APETALA1*s (*AP1*s) (A function for sepals and petals), two *APETALA3*s (*AP3*s) and two *PISTILLATA*s (*PI*s) (B function for petals and stamen), four *AGAMOUS*s (*AG*s) (C function for stamen and carpel) and six SEPALLATAs (*SEP*s) (E function for interacting with ABC function proteins) (Figure 5a). Collinearity analyses indicated that the expansion of type II MADS-box genes in *Greyia* resulted largely from the WGT event, including *AP1*, *SEP*, *SVP*, *AGL6*, and *SOC1* (Figure 5b-5d). For instance, *SEP-1*, *SEP-2*, and *SEP-3* are retained as triplicates and SEP-4 and SEP-5 present as duplicates following the WGT event (Figure 5b-5d and 5g-5i). In addition to WGT-derived duplications, tandem duplication has also contributed to MADS-box family expansion, as illustrated by paired *SEP* and *AP1* clusters. *SEP-4* (Grera02g02950) and *AP1-4* (Grera02g02960) originated from a tandem duplication on chromosome 2, which is syntenic with another tandem duplication on chromosome 7 – *SEP-5* (Grera07g01690) and *AP1-5* (Grera07g01680) (Figure 5b-5d). Sequence alignment showed only 52% identity between *SEP-4* and *AP1-4*, compared with 84% for *SEP-4*–*SEP-5* and 76% for *AP1-4*–*AP1-5* identity (Figure 5e), supporting tandem duplication predating the WGT (Figure 5f). Interestingly, the tandem-derived clusters exhibit distinct expression profiles: *SEP-4* and *SEP-5* are highly expressed in flowers, *AP1-4* is expressed in flowers and xylem, and *AP1-5* in flowers and leaves (Figure 5g–5i), suggesting functional subfunctionalization among paralogs. These findings suggest that successive gene expansion in *Greyia* – first via tandem duplication and later through WGT – has generated a rich repertoire of MADS-box genes, likely contributing to the evolution of this genus’s showy floral morphology.

**Figure 5.**
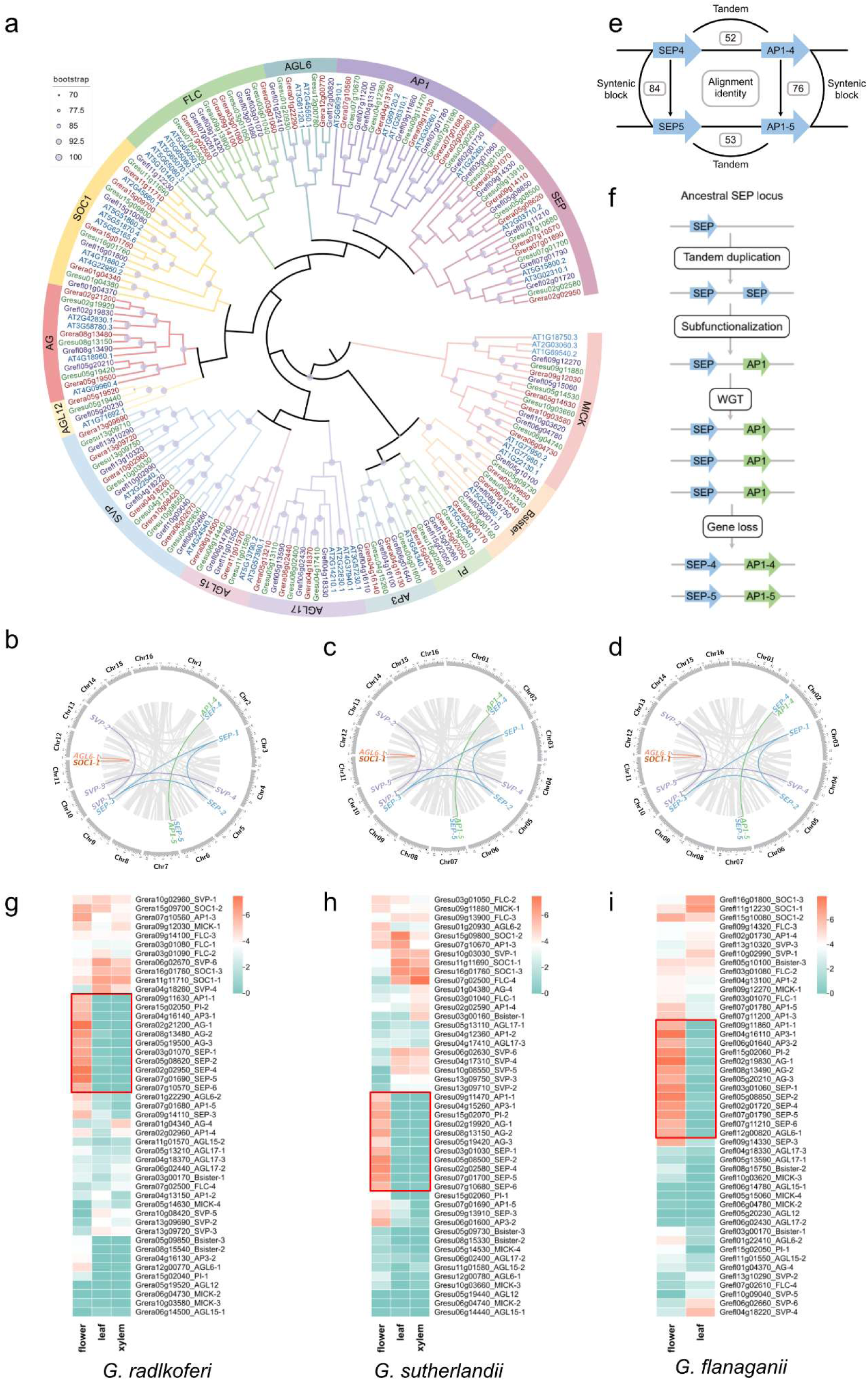
MADS-box genes in *Greyia* species. **a.** The phylogenetic tree of type II MADS-box genes in three *Greyia* and *Arabidopsis thaliana*. **b-d**. The syntenic relationships retained from the WGT event. Lines with different colors represent different gene families. **e**. Sequence alignment identity between two gene clusters (SEP – AP1-4 and SEP-5 – AP1-5). **f**. Reconstruction of the origin of gene clusters of SEP and AP1 (see Text for details). **g-i**. Gene expression patterns of type II MADS-box genes from various organs in the three *Greyia* species. The expression values are scaled by log2(TPM +1). TPM is transcripts per million. The red box highlights genes that are highly expressed in flowers, indicating the ABCE homologs in *Greyia*. The numbers following the gene are only used to distinguish different copies, the same as in Table S6

**Figure 6.**
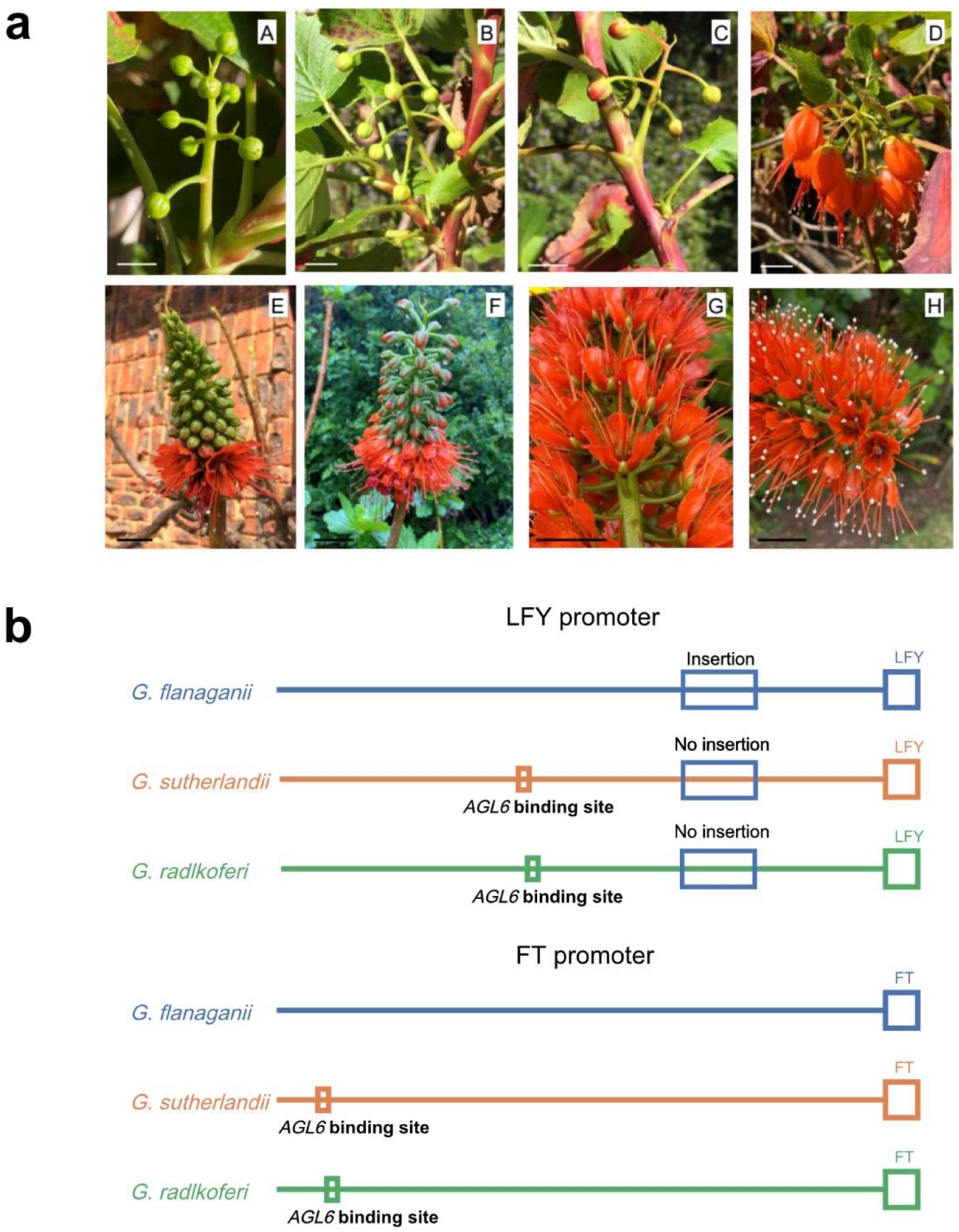
Potential regulation of inflorescence genes in *Greyia*. **a**. Inflorescence development from bud to anthesis in Greyia flanaganii and Greyia sutherlandii. a-c: G.flanaganii bud development, illustrating the long pedicels (1.5 – 3 cm). d: G. flanaganii inflorescence at anthesis, illustrating the typical number of flowers per inflorescence (<12), and the pendulous urceolate flower structure with unfused petals in imbricate formation. e-f: G. sutherlandii acropetal bud development, illustrating the many flowers per inflorescence (30-120). g-h: G. sutherlandii at anthesis, with short erect pedicels (< 1.2 cm), and valvate flowers. The panels represent stages 11-13 of flower development as defined for the model plant Arabidopsis in Alvarez-Buylla et al. (2010). G.radlkoferi is not shown since its inflorescence development is similar to G.sutherlandii. White scale bars = 1 cm; black scale bars = 2 cm. **b**. Cis-regulatory divergence of LFY and FT among Greyia species. Promoter lengths are scaled based on sequence alignments. For LFY, the promoter spans 2 kb upstream of the translation start site (ATG) in G. flanaganii, but is shorter in G. sutherlandii (1769 bp) and G. radlkoferi (1772 bp) due to the absence of a lineage-specific insertion. In G. flanaganii, this insertion is located 341–566 bp upstream of the LFY start codon. Predicted AGL6 binding sites are present in the LFY promoters of G. sutherlandii (–1026 to –1044 bp) and G. radlkoferi (–1030 to –1048 bp), based on motif similarity (p-value = 9.92e-06; q-value = 0.0237), but are absent in G. flanaganii. For FT, predicted AGL6 binding sites were detected in the FT promoters of G. sutherlandii (–1826 to –1844 bp) and G. radlkoferi (–1828 to –1846 bp), based on motif similarity (p-value = 2.09e-06; q-value = 0.00684), but not in G. flanaganii.

We next examined the expression profiles of type II MADS-box genes across flower, leaf, and xylem tissues in *Greyia* to identify candidate regulators associated with floral development. Among the five *AP1* paralogs, only *AP1-1*, the homolog of *APETALA1*, showed consistent, flower-predominant expression in all three species (Figure 5a, Figure 5g-5i), supporting its conserved association with floral tissues. For B-class genes, both *PI* and *AP3* have two homologs each, with *PI-2* and *AP3-1* showing high expression in flowers, suggesting that they likely cooperate to perform the B function (Figure 5g-5i). In addition, three C-function *AG* homologs and five E-function *SEP* homologs exhibited predominant floral expression, collectively contributing to floral organ identity, together with *API-1*, *PI-2*, and *AP3-1* (Figure 5g-5i). The expanded set of *AG* homologs in *Greyia spp.* contrasts with many angiosperms that retain only one or two highly expressed copies (Zhang *et al*., 2020, Chang *et al*., 2023, Ma *et al*., 2024). This may underlie the development of the characteristic exserted stamens observed in the *Greyia* genus, paralleling functional evidence from rice *AG* homologs *OsMADS3* and *OsMADS58*, where *OsMADS3* knockout results in stamen-to-lodicule transformations and ectopic lodicule formation (Dreni *et al*., 2011, Castelan-Munoz *et al*., 2019).

Within this conserved framework, we further examined *AGL6*, a close relative of *AP1* and *SEP*. Previous studies have shown that *AGL6*-like genes display diverse expression patterns and functional associations across angiosperms: in water lily, *AGL6* is expressed in sepals and petals, suggesting A-function substitution for *AP1*, whereas in petunia and rice, *AGL6-like* genes show *SEP*-like expression patterns and participate in higher-order MADS-box complexes (Rijpkema *et al*., 2009, Seok *et al*., 2010). In *Greyia*, two *AGL6* paralogs were identified in all three species. *AGL6-1* exhibited flower-specific expression, whereas *AGL6-2* showed broader expression across flower, leaf, and xylem tissues (Figure 5g–5i). Quantitative differences of *AGL6-1* floral expression were observed among species. *AGL6-1* showed lower floral expression compared to *AP1-1* in *G. radlkoferi* and *G. sutherlandii*, however, in *G. flanaganii*, *AGL6-1* is highly co-expressed with *AP1-1* in flowers (Figure 5g–5i). Phylogenetically, *AGL6* is closely related to *AP1*, and together with *SEP*, these subfamilies form a monophyletic superclade, suggesting a common ancestral origin predating the angiosperm/gymnosperm divergence (Becker and Theissen, 2003, Morel *et al*., 2019). Although the precise relationships among the subfamilies are not fully resolved, our analysis places *SEP* and *AP1* as sister groups, with *AGL6* sister to this clade (Figure 5a). These expressions and close phylogenetic patterns identify *AGL6-1* as a flower-associated MADS-box gene in Greyia and highlight it as a candidate for future functional investigation, especially in *G. flanaganii*.

### Inflorescence development

*Greyia* species are characterized by sympodial branching of shoots with inflorescences forming at the apex but a single side branch developing, causing a layered architecture (Steyn *et al*., 1987). The inflorescences are unbranched, indeterminate racemes bearing conspicuous red, nectariferous flowers (Figure 1a-c). While *G. radlkoferi*, *G. sutherlandii*, and *G. flanaganii* share a similar raceme structure, they differ markedly in flower density. *G. radlkoferi* and *G. sutherlandii* produce densely packed racemes with 30-120 flowers and short pedicels (<1.2 cm) (Figure 6e-h), whereas *G. flanaganii* bears sparser racemes typically with fewer than 12 flowers and conspicuously greater inter-floral spacing (1.5-3 cm) (Figure 6a-d). This architecture resembles simple indeterminate inflorescences in *A. thaliana*, where inflorescence and floral meristem identity are governed by a conserved regulatory framework involving *TERMINAL FLOWER 1* (*TFL1*), *LEAFY* (*LFY*), and *FLOWERING LOCUS T* (*FT*) (Andres *et al*., 2015).

To investigate whether similar regulators underlie inflorescence development in *Greyia*, we analyzed *LFY*, *TFL1*, and *FT* across species. Comparative analyses revealed that all three genes display highly conserved coding sequences and gene structures (Figure S11). However, *LFY* and *FT* show clear species-specific divergence at the promoter level, whereas no such divergence was detected for *TFL1*, indicating regulatory rather than structural variation. *LFY* is retained as a single-copy gene and exhibits extensive regulatory potential at the promoter level. Promoter analyses of LFY (see Methods) predicted 58 putative transcription factor (TF) binding sites representing 30 TFs in *G. flanaganii*, compared with 53 sites for 26 TFs in *G. radlkoferi* and 46 sites for 25 TFs in *G. sutherlandii* (Table S7-S9). Multiple sequence alignments identified a 226-bp insertion in the *G. flanaganii LFY* promoter, which is absent in *G. radlkoferi* and *G. sutherlandii* (Figure 6b and Figure S12). Motif prediction using the JASPAR 2026 database (Ovek Baydar *et al*., 2025) indicated that this insertion harbors a putative *PHL12* binding site (Table S7), although the functional relevance of this motif remains unclear, as *PHL12* has not previously been implicated in flowering regulation. Moreover, a predicted *AGL6* binding site was detected in the *LFY* promoters of *G. radlkoferi* and *G. sutherlandii* but was absent in *G. flanaganii* (Figure 6b and Table S7-S9). Despite these cis-regulatory differences, *LFY* transcript abundance was extremely low across all sampled tissues, consistent with its restriction to meristematic domains and suggesting that *LFY* activity at early inflorescence stages is not captured by the present RNA-seq dataset. Although direct binding of *AGL6* to the *LFY* promoter has not been experimentally demonstrated, the known role of *AGL6* in promoting floral transition through activation of *FT* and repression of *FLC/FLM* suggests a potential contribution to species-specific modulation of LFY activity in Greyia (Yoo *et al*., 2011, Yu *et al*., 2017, Kong *et al*., 2022).

Each *Greyia* genome contains two *FT* paralogs from the *Greyia*-specific WGT enent; however, only one copy showed detectable expression in the sampled tissues and was therefore retained for further analysis, consistent with functional divergence, as reported for *FT* paralogs in poplar (Bohlenius *et al*., 2006). Promoter analysis of this expressed *FT* copy revealed 73 predicted TF binding sites corresponding to 18 TFs in *G. flanaganii*, and 86 sites representing 26 TFs in both *G. radlkoferi* and *G. sutherlandii* (Table S10-S12). Similar to *LFY*, a predicted *AGL6* binding site was detected in the *FT* promoters of *G. radlkoferi* and *G. sutherlandii* but was absent in *G. flanaganii* (Figure 6b and Table S10-S12), indicating species-specific divergence in the cis-regulatory landscape. Expression profiling showed that in the deciduous species *G. radlkoferi* and *G. sutherlandii*, *FT* transcripts were predominantly detected in mature flowers at anthesis, whereas no expression was observed in their leaf samples. By contrast, the evergreen species *G. flanaganii* exhibited *FT* expression in both flower and leaf tissues (Figure 6 and Figure S13). These spatial expression patterns are consistent with differences in growth habit and phenology, with evergreen species retaining leaves during flowering and deciduous species flowering prior to leaf flush, as described for other woody perennials (Bohlenius *et al*., 2006, Wilkie *et al*., 2008, Hsu *et al*., 2011). The detection of *FT* expression in mature floral tissues is also noteworthy, as transcriptomic studies in *Arabidopsis* typically report very low FT expression in flowers, reflecting its primary role during early floral transition rather than late floral development (Zeevaart, 2008, Liu *et al*., 2013). In *Greyia*, floral *FT* expression coincides with elevated floral expression of *AGL6-2* and the presence of predicted *AGL6* binding sites in *G. radlkoferi* and *G. sutherlandii*, compared to *G. flanaganii* (Figure S14). Together, these observations highlight coordinated expression and cis-regulatory features associated with floral tissues, but do not by themselves establish a direct regulatory relationship. Clarifying the regulatory relationships among *AGL6*, *FT*, and *LFY*, and evaluating their potential contribution to variation in inflorescence architecture, will require transcriptomic and spatial expression analyses encompassing shoot apical meristems, early floral transition stages, and multiple seasonal time points.

## Discussion

Greyia is a small genus comprising three species, and the three chromosome-scale genomes reported here constitute an important genomic resource for elucidating the molecular mechanisms underlying trichome and floral differentiation. Representing all currently recognized species – *G. radlkoferi*, *G. sutherlandii*, and *G. flanaganii* – these assemblies provide the first high-quality reference genomes for the order Geraniales and establish a valuable foundation for comparative genomics and evolutionary studies.

*Greyia* species possess relatively small genomes (∼180 Mb; Table 1) with low heterozygosity, posing little challenge for genome assembly. However, achieving a high-quality assembly does not automatically guarantee accurate annotation (Wu *et al*., 2024). For instance, while BUSCO assessments of recently published telomere-to-telomere (T2T) and gapless genomes often achieve impressive completeness rates of ∼98%, the evaluation of predicted genes typically falls slightly below this benchmark (Detcharoen *et al*., 2023, Nie *et al*., 2023, Xia *et al*., 2023). Indeed, the compact genome structure of *Greyia* genomes frequently led to adjacent genes being linked together. Here, we took advantage of comprehensive gene predictions and extensive manual curations of gene models (see Methods), achieving a BUSCO completeness score exceeding 99.3%, and over 80% of genes that were supported by transcriptomic evidence – an improvement over many previously reported plant genomes.

The phylogenetic placement of Geraniales has long remained uncertain, largely due to the limited availability of genomic resources from key lineages. By leveraging phylogenomic analyses based on 401 single-copy genes identified across our sampled taxa, we recovered a well-supported sister-group relationship between *Greyia* and *E. japonica*, thereby refining the placement of Geraniales within the superrosids. This topology is consistent with recent studies proposing a Crossosomatales-Geraniales sister relationship (Zuntini *et al*., 2024) and is further supported by phylogenomic inferences from the *E. japonica* genome. Notably, the superrosid topology inferred here differs from that recovered in previous large-scale angiosperm phylogenomic analyses based on the Angiosperms353 nuclear gene set (Zuntini *et al*., 2024). Comparison between our dataset and Angiosperms353 revealed that only a limited subset of loci is shared between the two, reflecting their distinct design principles. Whereas Angiosperms353 loci were selected for broad applicability and phylogenetic informativeness across the entire angiosperm radiation (Zuntini *et al*., 2024), the 401 genes analyzed here represent lineage-specific single-copy markers tailored to resolving relationships within our focal clade. Such differences in gene sampling, together with variation in taxon coverage and analytical strategies, likely contribute to the observed topological discrepancies and underscore the sensitivity of deep superrosid phylogenetic inference to dataset choice. Resolving these deep relationships will therefore require the integration of additional genomic resources and the application of complementary phylogenomic frameworks (Chen *et al*., 2019, Stull *et al*., 2020, Sanderson *et al*., 2023, Huang *et al*., 2025).

Within Geraniales, the combined genomic and transcriptomic evidence supports two major families, consistent with the previous result (Palazzesi *et al*., 2012). Within Francoaceae, the closest relationship between *Greyia* (South Africa) and *Francoa* (South America) points to a historical biogeographic connection between these lineages across Gondwana. This pattern may reflect either ancient geographic separation following the breakup of Gondwana, with descendants persisting on different continents, or from later long-distance dispersal across the South Atlantic, followed by diversification in isolation. Divergence dating further indicates that *Greyia* diversified relatively recently during the Quaternary (≤ 1.4 Mya), likely influenced by climatic fluctuations that promoted ecological specialization across southern Africa (Fletcher and Thomas, 2010, Paillard, 2015).

Polyploidization plays a crucial role in plant genome evolution and adaptation to environmental change (Van de Peer *et al*., 2017). Using multiple complementary approaches, we identified a *Greyia*-specific WGT event dated to ∼86.97 Mya, in addition to the core-eudicot Gamma event. This WGT has had a profound impact on *Greyia* genome architecture, with approximately 40% of duplicated genomic regions retained, representing an unusually high level of preservation. Among the retained duplicated and triplicated genes, we observed significant enrichment for functions associated with rhythmic and circadian processes. Notably, WGT-derived genes include key components of the photoperiod and circadian clock pathways involved in light perception (*CRY1*), core oscillation (*CCA1/LHY1*, *PRR5*, *PRR7*), and downstream regulation of flowering (*GI*, *EBS*). Their coordinated retention within WGT-derived syntenic blocks suggests that polyploidy preserved an intact regulatory context rather than isolated gene copies. Photoperiodic regulation is a key mechanism by which plants align developmental transitions with seasonal environmental cues, particularly in temperate and subtropical regions. Although the present data do not establish a direct functional role for WGT-derived photoperiod genes in shaping *Greyia* phenotypes, their retention patterns are consistent with a potential contribution to the regulation of seasonal growth habit between the evergreen *G. flanaganii* and the other two deciduous species. Tissue-specific expression of these RTGs further indicate functional specialization, implying that duplicated genes have been co-opted for distinct roles in different organs or developmental stages. Preferential retention of circadian and photoperiod regulators following genome duplication has also been reported in other angiosperm lineages, supporting the idea that these pathways may be particularly dosage-sensitive or constrained by network connectivity (Adams and Wendel, 2005, Soltis *et al*., 2015, Van de Peer *et al*., 2017, Van de Peer *et al*., 2021). Together, these findings indicate that the *Greyia*-specific WGT provided a genomic substrate upon which regulatory diversification could occur, with possible consequences for life-history variation among species. Functional validation across developmental stages and seasonal contexts will be required to determine the extent to which retained photoperiod genes contribute to species-specific traits in *Greyia*.

Leaf and floral morphology represent the primary traits distinguishing *Greyia* species, yet their molecular bases have remained poorly understood. Our analyses indicate that contrasting leaf trichome phenotypes between *G. radlkoferi* and *G. sutherlandii* are not explained by differences in gene content, but instead reflect regulatory divergence within a structurally conserved trichome initiation network. A central feature of this divergence is the differential expression of *GL1*. Although *GL1* is retained as four paralogs following the *Greyia*-specific WGT event, only a single copy exhibits strong leaf-specific expression in *G. radlkoferi*, consistent with a conserved role in trichome initiation, whereas the corresponding *GL1* homolog in *G. sutherlandii* is transcriptionally silent across all examined tissues. Regulatory divergence extends to *GL3/EGL3*, another core component of the MBW complex, which shows leaf-enriched expression in *G. radlkoferi* but flower-biased expression in *G. sutherlandii*. This shift in tissue-specific expression is consistent with the context-dependent roles of *GL3/EGL3* described in *Arabidopsis*, where the same bHLH factors participate in distinct epidermal or pigmentation pathways depending on their MYB partners (Oppenheimer *et al*., 1991, Nesi *et al*., 2001, Gonzalez *et al*., 2008). Coordinated down-regulation of upstream regulators, including *GIS3* and *GIS/GIS2*, further suggests attenuation of the trichome initiation module at the network level rather than isolated gene-specific effects. Importantly, polyploidization has expanded the copy number of key MBW-associated genes in *Greyia*, providing a substrate for differential expression and partitioning of regulatory roles among paralogs. Previous studies have demonstrated that promoter DNA methylation is generally associated with transcriptional repression in plants (Dubin *et al*., 2015, Zhang *et al*., 2018). Together with differential promoter methylation detected in *G. sutherlandii*, these patterns suggest that regulatory divergence operating at both transcriptional and epigenetic levels, facilitated by post-WGT gene retention, underlies species-specific variation in trichome development.

In contrast to the pronounced regulatory divergence underlying leaf traits, floral morphological differences among *Greyia* species appear to arise without major changes to the core floral identity gene network. All three species share an identical complement of type II MADS-box genes, including conserved ABCE-class homologs with largely similar floral expression profiles, indicating strong conservation of the genetic framework specifying floral organ identity. Despite the striking contrast between the urceolate, imbricate flowers of *G. flanaganii* and the open, valvate flowers of *G. sutherlandii* and *G. radlkoferi*, we detected no species-specific gains, losses, or qualitative expression shifts among canonical ABCE genes, suggesting that floral diversification did not arise from rewiring of the core MADS-box program. Instead, finer-scale regulatory modulation within this conserved framework is likely responsible. Expansion of *SEP* genes following the Greyia-specific WGT provides a robust genetic substrate for floral organ patterning, while *AG* gene expansion may underlie the characteristic exserted stamens observed across the genus (Dreni *et al*., 2011, Castelan-Munoz *et al*., 2019).

In the inflorescence development, both *LFY* and *FT* show promoter-level differentiation among species, despite conservation of gene copy number and coding sequence, indicating that regulatory rather than structural evolution underlies phenotypic diversification in *Greyia* inflorescences. In particular, the presence of predicted *AGL6* binding sites in the promoters of *G. radlkoferi* and *G. sutherlandii*, but not *G. flanaganii*, points to lineage-specific modification of upstream regulatory context. In evergreen *G. flanaganii*, retention of leaves during flowering is compatible with a regulatory mode resembling canonical leaf-derived *FT* signaling, whereas deciduous species that flower prior to leaf flush are more consistent with perennial tree strategies in which floral induction and initiation are temporally separated (Wilkie *et al*., 2008, Hsu *et al*., 2011). Interpretation of these patterns must consider the temporal constraints of transcriptomic sampling, as *FT* activity typically precedes anthesis and is restricted to narrow inductive windows. In this context, it is notable that the leaves of *G. radlkoferi* used for RNA-seq were collected in late summer when the tree had not flowered for several years and did not flower again until spring of the next year, suggesting that the sampled tissues were unlikely to represent a floral inductive state. Thus, the absence of detectable *FT* expression in leaves of deciduous *Greyia* species is most parsimoniously explained by sampling context rather than by a lack of *FT* involvement in flowering regulation. The detection of *FT* transcripts in mature floral tissues of all three species is nevertheless intriguing and may reflect diversification of *FT* spatial or temporal roles, as reported for woody perennials (Wilkie *et al*., 2008). Overall, our results suggest that diversification of inflorescence architecture in *Greyia* has occurred primarily through regulatory divergence and life-history–dependent modulation of conserved flowering pathways, rather than through changes to core meristem identity genes. Dissecting the precise roles of *FT*, *LFY*, and *AGL6* during early floral induction versus later stages of floral development will require transcriptomic and spatial expression profiling of shoot apical meristems, combined with seasonal sampling across multiple developmental stages.

### Conclusion

This study presents chromosome-scale genome assemblies for all three extant *Greyia* species, representing the first high-quality genomic resources for the order Geraniales. Integrating comparative genomics with transcriptomic analyses, we show that polyploidy and regulatory diversification, rather than changes in gene content, underlie phenotypic variation in leaf trichomes as well as floral and inflorescence architecture. The retention of duplicated regulators following a *Greyia*-specific WGT provided the genetic substrate for subfunctionalization and regulatory reprogramming of conserved pathways. These high-quality genomic resources establish a framework for functional dissection of developmental gene networks in Geraniales using Greyia as a model system. They can be deployed to elucidate the molecular basis of morphological differentiation in *Greyia*, for example adaptation to pollinators through floral architecture and nectary diversification (Jeiter *et al*., 2017b). Furthermore, they will provide the foundation for studies of adaptation to pests, pathogens, and abiotic stresses through chemical defences (Bohm and Chan, 1992), which will underpin exploitation of the medicinal properties of the *Greyia* genus (Botha *et al*., 2026). Additionally, these genomic resources provide the resolution necessary to clarify the molecular systematics of *Greyia*, surpassing the limitations of standard DNA barcoding (Botha *et al*., 2026). The establishment of reference genomes for each species facilitates robust population genomics across the genus’s 1,000 km range in southern Africa, offering a critical framework for evaluating species concepts in the face of recent divergence and the sympatric ranges of *G. radlkoferi* and *G. sutherlandii*. By securing the taxonomic identity of this keystone grassland genus, this would directly inform conservation strategies. On a macro-evolutionary scale, these data will help resolve a long-standing phytogeographic conundrum, namely that *Greyia* has closest phylogenetic affinity with *Francoa*, a genus endemic to Chile, and not other African taxa in the Geraniales (*Melianthus, Bersama*) (Sytsma *et al*., 2014).

## EXPERIMENTAL PROCEDURES

### Plant materials

*Greyia* trees sampled in this study are part of a larger study where each individual tree was given a unique “P” number. Plant samples for gDNA and RNA extraction were obtained from the following individual trees. *Greyia sutherlandii* Hook. & Harv. tree P245 growing at Monks Cowl Nature Reserve, Maloti-Drakensberg Park World Heritage Site, KwaZulu-Natal Province, South Africa. Sampling was conducted under Research Permits OP34/2022, OP737/2023, and OP2781/2024 issued by Ezemvelo KZN Wildlife, South Africa. *Greyia flanaganii* Bolus tree P261 at the Manie Van der Schijff Botanical Garden (accession number 4631), University of Pretoria, South Africa. *Greyia radlkoferi* Szyszyl. tree P262 at the Manie Van der Schijff Botanical Garden (accession number 4632), University of Pretoria, South Africa. *Greyia* species identities were determined based on flower and leaf morphological differences (Figure 1) verified against herbarium specimens at the HGWJ Schweickerdt Herbarium (PRU) at the University of Pretoria, South Africa. Representative voucher specimens of *G. sutherlandii* P245, *G. radlkoferi* P262 and *G. flanaganii* P261 have been deposited at this herbarium with accession numbers PRU0132900, PRU0133318, PRU0133319, respectively. For high molecular weight DNA, fully expanded leaves at leaf flush stage were collected from *G. sutherlandii* P245, *G. flanaganii* P261, and *G. radlkoferi* P262, flash frozen in liquid nitrogen, and stored at –80°C. The protocol described by Zerpa-Catanho et al. (Zerpa-Catanho *et al*., 2021) was followed to obtain high-quality high molecular weight intact DNA: in short, nuclei were first isolated after thoroughly pulverising 5 g leaf material in liquid nitrogen using a nuclei isolation buffer (10 mM Tris-HCl, 10 mM EDTA, 100 mM KCl, 0.5 M sucrose, 4 mM spermidine, 1 mM spermine) to extract (supplemented with 10% Triton X-102) and wash (supplemented with 0.1% β-mercaptoethanol) the nuclei, from which DNA was extracted using a DNA extraction buffer (100 mM Tris-HCl, 50 mM EDTA, 500 mM NaCl, 1.25% SDS). The DNA was isolated from the extract by performing proteinase K and RNase A treatments, and using one volume phenol:chloroform: isoamyl alcohol (15:14:1) and chloroform: isoamyl alcohol (14:1) in subsequent steps to separate out cellular debris and other macromolecules. An equal volume of isopropanol was used to precipitate the DNA from solution, which was purified using a QIAGEN Genomic Tip 20/G (QIAGEN, Cat no. / ID. 10223). The DNA was resuspended in TE buffer and stored at 4°C prior to shipping to Macrogen Europe B.V., Amsterdam for sequencing.

RNA was extracted using a modified CTAB protocol with LiCl precipitation (Chang *et al*., 1993) – in short, 2g of flash-frozen tissue was ground to a fine powder in liquid nitrogen, to which pre-warmed CTAB buffer (2 M NaCl, 25 mM EDTA, 100 mM Tris-HCl (pH 8.0), 2% PVP-10, 2% CTAB, 3.44 mM spermidine, 2% ß-mercaptoethanol) was added. Samples were incubated for 10 minutes at 65°C before an equal volume of chloroform:isoamyl alcohol (14:1) was added, and shaken vigorously. Centrifugation was performed at 8500 *g* at 4°C for 15 minutes, and the upper aqueous phase was transferred to a new, pre-chilled Falcon tube. The chloroform:isoamyl alcohol extraction was repeated two more times. A ¼ volume 10 M LiCl was then added to precipitate the RNA overnight at 4°C, which was pelleted by centrifugation at 8500 *g* at 4°C for 90 minutes. Following the removal of the LiCl solution, the pellet was washed with ice-cold 70% ethanol twice, with centrifugation performed at 8500 *g* at 4°C for 15 minutes. The pellet was air-dried at room temperature and resuspended in RNase-free water.

For DNase treatment, the RNA solution was first diluted to 0.2 μg/μl before 10X DNase I buffer (100 mM Tris (pH7.5), 25 mM MgCl_2_, 5 mM CaCl_2_) and 2 U Ambion DNase I was added and incubated for 30 minutes at 37°C. The Monarch® Spin RNA Cleanup Kit (50 μg (New England Biolabs®, T2040) was then used to purify the RNA, and quality was assessed through Qubit assay (Invitrogen) and NanoDrop spectrophotometry (ThermoFischer Scientific). RNA was precipitated from solution with 0.1 volumes 3 M NaOAc (pH 5.5) and 2 volumes 100% ethanol for shipping to Macrogen Europe B.V. on ice.

### Genome sequencing

Long-read nanopore sequencing was carried out on the PromethION sequencer (Oxford Nanopore Technologies, Oxford, UK) to assemble the genome. Genomic DNA by Ligation (SQK-LSK110) kit was used to generate sequencing libraries following the manufacturer’s protocol, and nanopore sequencing was performed on the R10.4.1 flow cells. Raw data was obtained from all PromethION runs and basecalling was performed with Dorado (v0.5.3).

Libraries were prepared for short-read Illumina whole-genome sequencing using the TruSeq® Nano DNA Library Prep Kit according to the protocol guide (Part # 15041110 Rev. D). The DNA was first fragmented to 550 bp fragments through Covaris shearing, leaving 3’ and 5’ overhangs, which were subsequently removed and repaired to produce blunt ends. The fragment 3’ ends are then adenylated to prevent ligation between fragments and provide a complementary overhang for the adaptors to anneal. Following ligation of the indexing adaptors, the fragments are selectively enriched through PCR amplification with primers complementary to the adaptors. The library concentrations are then normalised to 10 nM and pooled in equal volumes in the Pooled Diluted Cluster Template Plate for cluster generation. Paired end sequencing was performed using the Illumina NovaSeq platform of Macrogen Europe B.V.

Hi-C sequencing was conducted on the Illumina NovaSeq X plus system using the 150-bp pair-end reads by Biomarker Technologies (BMK) GmbH, Münster, Germany. The Hi-C libraries were constructed to anchor scaffolds onto chromosomes (Burton *et al*., 2013). DNA isolated from *G. radlkoferi* leaves was digested with HindIII overnight. Sticky ends were biotinylated and proximity-ligated, and then physically sheared to a size of 300-700 bp and enriched to make chimeric junctions. The cross-linked long-distance physical connections were next processed into chimeric fragments, followed by reverse cross-linking, purification, and PCR amplification, and were subsequently used to create paired-end sequencing libraries.

For transcriptome sequencing, sequencing libraries were prepared using the TruSeq® Stranded mRNA LT Sample Prep Kit according to the protocol guide (#15031047 Rev. E) – in short, mRNA was first purified with oligo-dT attached magnetic beads, fragmented, and used for cDNA synthesis. The 3’ ends of the cDNA were then adenylated to allow for adaptor ligation, following which the fragments are enriched through PCR amplification with primers specific to the adaptor sequences. Finally, the libraries are validated through quantification and quality control. The libraries were then loaded onto flow cells, with surface-bound oligonucleotides capturing the complementary adaptors, and clonal clusters were produced from the captured fragments with bridge amplification. Paired end sequencing was performed using the Illumina NovaSeq platform of Macrogen Europe B.V. Genome size and heterozygosity estimation

Genome size and heterozygosity were estimated by K-mer frequency distribution analysis. Initially, the short reads were trimmed using Trimmomatic (v0.39) (Bolger *et al*., 2014) with the parameters “ILLUMINACLIP: TruSeq3-PE-2.fa:2:30:10 LEADING:20 TRAILING:20 SLIDINGWINDOW:10:20 MINLEN:50”. K-mers were then counted using Jellyfish (v2.3.0) (Marcais and Kingsford, 2011) with the parameters “-C –m 21 –s 4G –t 5”. The resulting output file was subsequently used as input for GenomeScope2.0 (Ranallo-Benavidez *et al*., 2020), which estimated the genome size and heterozygosity using default parameters.

### Genome assembly and evaluation

Prior to the assembly, we performed quality trimming on the Nanopore data using Porechop (v0.2.4; https://github.com/rrwick/Porechop). Draft contig-level assemblies were then generated with Flye (v2.9.3) using Nanopore data under optimized parameters (--asm-coverage 50; –-nano-hq).

Hi-C sequencing reads were processed by removing adapters and filtering low-quality sequences. These processed reads were then aligned to the contig-level assembly using BWA (v0.7.17) (Li and Durbin, 2009), with the resulting BAM file serving as input for Juicer (v1.9.8) (Durand *et al*., 2016b). Scaffolding was performed using 3D-DNA (v180922) (Dudchenko *et al*., 2017) to cluster, order, and orient the contigs. Finally, we manually corrected misassembled scaffolds in Juicerbox (v1.11.08) (Durand *et al*., 2016a) by examining Hi-C contact frequency patterns. Genome completeness of the final assemblies was assessed using Benchmarking Universal Single-Copy Orthologs (BUSCO, v5.7.1) (Manni *et al*., 2021) with the eudicots_odb10 dataset.

### Repeat prediction

A preliminary list of candidate LTR retrotransposons (LTR-RT) was generated using LTR_FINDER_parallel (Ou and Jiang, 2019) and LTR_harvest (v1.5.9) (Ellinghaus *et al*., 2008). A repeats library was constructed for each *Greyia* species to include LTR-RT and other TE elements identified by LTR_retriever (v3.0.1) (Ou and Jiang, 2018) and RepeatModeler (v2.0.1) (Flynn *et al*., 2020), respectively. The repeats library was used as input for RepeatMasker (v4.1.1; http://www.repeatmasker.org/RepeatMasker/) to predict and mask repeats in three *Greyia* genomes.

### Structural and functional annotation

Gene structure annotation in the three *Greyia* genomes was performed by integrating RNA-based, homology-based, and ab initio predictions. For RNA-based prediction, RNA-seq data were first mapped to the assemblies using HISAT2 (v2.1.0; parameter: –dta) (Kim *et al*., 2019), and subsequently assembled into transcripts by StringTie2 (v1.3.6) (Kovaka *et al*., 2019). All transcripts from RNA-seq were combined using Cuffcompare (v.2.2.1) (Trapnell *et al*., 2010) and subsequently merged with Stringtie2 to remove fragments and redundant structures. TransDecoder (v5.0.2; https://github.com/TransDecoder/TransDecoder) was then used to predict open reading frames. *Arabidopsis thaliana* and *Euscaphis japonica* genomes and annotations were used for homology-based prediction using GeMoMa (v1.9) (Keilwagen *et al*., 2016, Keilwagen *et al*., 2018). BRAKER3 (v3.0.8) was used for *ab initio* prediction using model training based on RNA-seq data.

All structural gene annotations were joined with EvidenceModeller (v1.1.1) (Haas *et al*., 2008), and BUSCO (v5.7.1) and OMArk (v0.3.1) (Nevers *et al*., 2024) were used to assess the quality of the annotation results. Finally, we used GenomeView (Abeel *et al*., 2012) to perform the gene curation manually based on RNA-seq data alignments, including (1) split incorrectly linked gene models, (2) added missing genes supported by RNA-seq evidence, and (3) merged fragmented annotations where RNA-seq indicated single transcripts. Putative gene functions were identified using InterProScan (v5.65) (Jones *et al*., 2014) and eggNOG-mapper (v2.1.12) (Cantalapiedra *et al*., 2021).

### Whole-genome duplication identification and dating

*K_S_* age distribution analysis was performed using the wgd v2 package (Chen *et al*., 2024). Anchor pairs (that is, paralogous genes lying in collinear or syntenic regions of the genome) were identified using i-ADHoRe (Proost *et al*., 2012). To account for substitution rate heterogeneity among lineages, *K_S_* distribution analysis among species was also corrected using the KSRATES software (Sensalari *et al*., 2022), which locates ancient polyploidization events with respect to speciation events within a phylogeny, comparing paralog and ortholog *K_S_* distributions, while correcting for substitution rate differences across the involved lineages. Visualization of the *K_S_* distribution was performed with Hiplot (https://hiplot.cn/).

The absolute age (in geological time) of the WGT event in *Greyia* and the WGD event in Euscaphis was also dated using the wgd v2 package(Chen *et al*., 2024). In brief, paralogous gene pairs located in duplicated segments (so-called anchor genes, anchor pairs, or anchors) and duplicated pairs lying under the WGD peak (so-called peak-based duplicates) were collected for phylogenetic dating. Anchors, assumed to be corresponding to the most recent WGD, were detected using i-ADHoRe (Fostier et al., 2011; Proost et al., 2012; v3.0). We selected anchor pairs and peak-based duplicates present under the *Greyia* WGD peak and with *K_S_* values between 0.3 and 0.8 for absolute dating. For each supposed WGD paralogous pair, an orthogroup was created that included the two paralogs plus several orthologs from other plant species as identified by InParanoid (Ostlund et al., 2010; v4.1) using a broad taxonomic sampling: *Lotus japonicus*, *Vigna mungo*, *Tripterygium wilfordii*, *Poncirus trifoliata*, *Mangifera indica*, *Pistacia vera*, *Theobroma cacao*, *Durio zibethinus*, *Gossypium raimondii*, *Arabidopsis thaliana*, *Tarenaya hassleriana*, *Cleome violacea*, *Greyia radlkoferi*, *Paeonia ostii*, and *Kalanchoe fedtschenkoi*. In total, 780 orthogroups based on anchors and peak-based duplicates were collected for the phylogenetic dating. The pipeline can be divided into three main steps. 1) The construction of anchor *K_S_* distribution and the delineation of credible *K_S_* range adopted for phylogenetic dating, using “wgd dmd”, “wgd ksd”, “wgd syn”, and “wgd peak”. 2) The formulation of a starting tree used in the phylogenetic dating, composed of a broad taxonomic species with annotated fossil calibration information, using “wgd dmd –-globalmrbh”; 3) The construction of orthogroups consisting of collinear duplicates of *Greyia* species and their reciprocal best hits (RBHs) against other species in the starting tree and the phylogenetic dating using a molecular dating program MCMCtree with the parameters “burnin = 2000 –ds sampfreq = 1000’ –ds nsample = 20000” via wgd focus. Finally, 780 orthogroups were accepted, and absolute age estimates of the node uniting the WGD paralogous pairs based on both anchor pairs and peak-based duplicates were grouped into one absolute age distribution, for which kernel density estimation and a bootstrapping procedure were used to find the peak consensus WGD age estimate and its 90% CI boundaries, respectively.

### Phylogenetic tree construction and estimation of divergence times

To retrieve the evolutionary history of Geraniales, we chose one or two representative species in the close orders: Brassicales, Malvales, Sapindales, Oxalidales, Malpighiales, Celastrales, Fagales, Rosales, Cucurbitales, Fabales, Geraniales, Crossosomatales, Zygophyllales, and Myrtales, as well as two monocot outgroups (*Oryza sativa* and *Brachypodium distachyon*). Orthologous groups were identified using OrthoFinder (v2.5) (Emms and Kelly, 2019). From these results, single-copy orthologous genes were extracted. Protein sequences of each single-copy ortholog were aligned with MAFFT (Katoh and Standley, 2013) using default parameters, and the resulting protein alignments were converted into codon alignments. Poorly aligned or divergent regions were removed using trimAl (Capella-Gutierrez *et al*., 2009). The filtered codon alignments from all single-copy orthologs were then concatenated into a supergene matrix for species phylogenetic reconstruction. A maximum-likelihood phylogenetic tree was inferred from both the protein and codon alignments using IQ-TREE (Nguyen *et al*., 2015) under the GTR+G model with 1,000 bootstrap replicates. Divergence times among 19 plant species were estimated with MCMCtree in the PAML package (Yang, 2007), employing the GTR+G substitution model. Fossil-based calibration points included: the divergence between monocots and eudicots (125-247 Mya; based on tricolpate pollen grain)(Dilcher *et al*., 2007), the split between Brachypodium distachyon and Oryza sativa (42-52 Mya)(Catalan *et al*., 2012), and the divergence between Fabids and Malvids (102-125 Mya)(Wang *et al*., 2009). For Bayesian inference, MCMCtree was run to generate 10,000 posterior trees, sampling every 150 iterations after a burn-in of 500,000 iterations. Two independent runs were compared to assess convergence, and the posterior distribution of node ages was also compared with the effective prior implied by fossil calibrations.

### Genome and gene family evolution

Synteny analyses were conducted using MCScanX with the parameter “-s 10.” Genes duplicated or triplicated as a result of whole-genome triplication (WGT) events were visualized with Circos (Krzywinski *et al*., 2009). Syntenic relationships among *Euscaphis*, and *Greyia* were further identified using JCVI (https://github.com/tanghaibao/jcvi). Gene Ontology (GO) enrichment analyses were performed with the R package clusterProfiler (Wu *et al*., 2021).

For expression analysis, RNA-seq data were first mapped to the assemblies using HISAT2 (v2.1.0; parameter: –dta), and then raw reads counts were obtained using featureCounts (v2.0.1) (Liao *et al*., 2014), TPM was calculated with a custom script.

For MADS-box gene identification, a hidden Markov model (HMM) approach was applied to the three *Greyia* genomes. The HMM profile for the SRF-TF domain (PF00319) was retrieved from the Pfam database (Mistry *et al*., 2021). Candidate MADS-box genes were identified by searching the local protein databases with HMMER (Eddy, 2011) using an *E*-value cutoff of <1e-5. MADS-box proteins of Arabidopsis thaliana were identified using the same pipeline. Candidate sequences from all four species were then aligned with MAFFT and concatenated for phylogenetic reconstruction, and the resulting gene tree was visualized with iTOL (https://itol.embl.de/login.cgi). Trichome-related genes in *Greyia* were identified via homology searches against *A. thaliana*. RNA expression profiles of MADS-box and trichome-related genes were visualized using Chiplot (https://www.chiplot.online/).

For promoter analysis, promoter regions were operationally defined as the 2-kb sequence upstream of the translation start site (ATG) (or upstream of the annotated TSS when available) for cis-element/TFBS scanning, following common practice in plant promoter studies (Thibaud-Nissen *et al*., 2006, Baek *et al*., 2011, Brooks *et al*., 2023). Thus, the 2-kb sequence upstream of each target gene was extracted as the putative promoter region. Motif discovery was performed with MEME (http://meme-suite.org) to identify statistically overrepresented sequence patterns. The predicted motifs were subsequently annotated and compared against transcription factor binding site databases using Tomtom (https://meme-suite.org/meme/tools/tomtom), thereby enabling functional inference of *cis*-regulatory elements.

For DNA methylation analysis, Oxford Nanopore reads were basecalled using Dorado with modified base detection enabled (--modified-bases-models), allowing the identification of methylated bases directly from raw signal data. Basecalled reads were aligned to the reference genome using the Dorado aligner, and methylation calls were extracted from the resulting BAM files using modkit and converted to BED format for downstream analyses.

## Acknowledgements

Funding for this research was provided by the National Research Foundation South Africa (NRF) Foundational Biodiversity Information Programme grant to DKB (FBIS2204041924), an Oppenheimer Memorial Trust (OMT) Sabbatical Award to DKB (OMT Ref. 2023-1431), a University of Pretoria Staff Travel Abroad grant to DKB, and OMT MSc Scholarship to JB (OMT Ref. 2024-5181). The authors wish to acknowledge the Manie van der Schijff Botanical Garden, Department of Plant and Soil Sciences, University of Pretoria for providing *Greyia* plant material, T. Duong for assistance with *Greyia* high molecular weight gDNA extraction, and M. Robertson, Ezemvelo KZN Wildlife, for sampling support. YVdP acknowledges funding from the European Research Council (ERC) under the European Union’s Horizon 2020 research and innovation program (No. 833522) and from Ghent University (Methusalem funding, BOF.MET.2021.0005.01).

## Author contributions

Xiao Ma: data curation, formal analysis, software, visualization, writing – original draft (lead), methodology (equal). Jiyang Chang: data curation, formal analysis, software, visualization (lead), methodology, writing – review & editing (equal). Jana Botes: investigation (lead), methodology, visualization, writing – original draft (supporting). Yves Van de Peer: resources, funding acquisition (lead), conceptualization, project administration, supervision, writing – review & editing (equal). Dave K. Berger: conceptualization, project administration, supervision, writing – review & editing (equal), resources, funding acquisition (supporting).

## Data availability statement

All genome assemblies, structural and functional annotations, and protein-coding genes are deposited in the ORCAE database at https://bioinformatics.psb.ugent.be/gdb/Greyia/.

## Supplementary Figures

**Figure S1.**
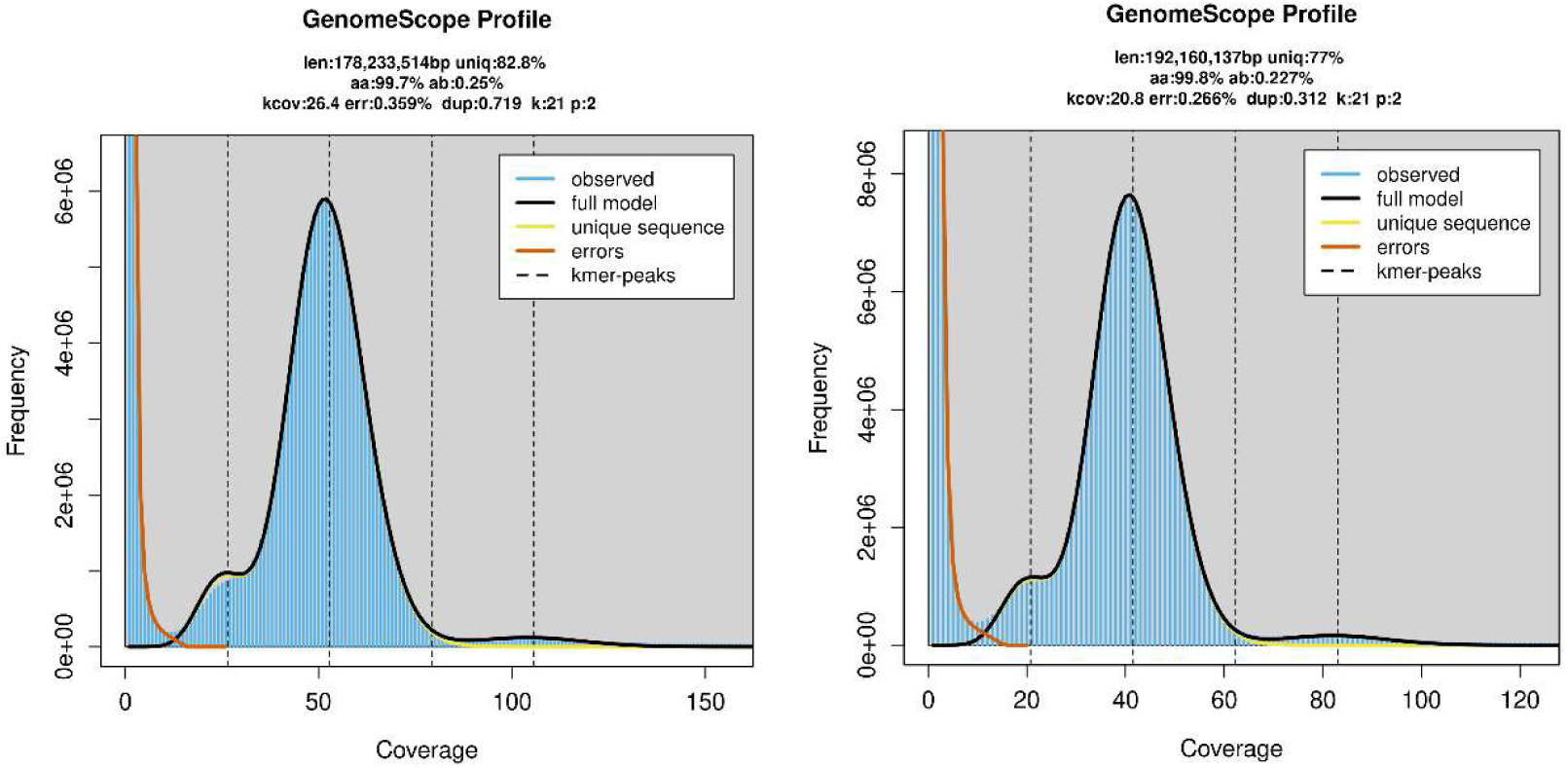
Genome size and heterozygosity of *G. radlkoferi* (Left) and *G. flanaganii* (right) based on 21-mer analysis using Illumina short reads.

**Figure S2.**
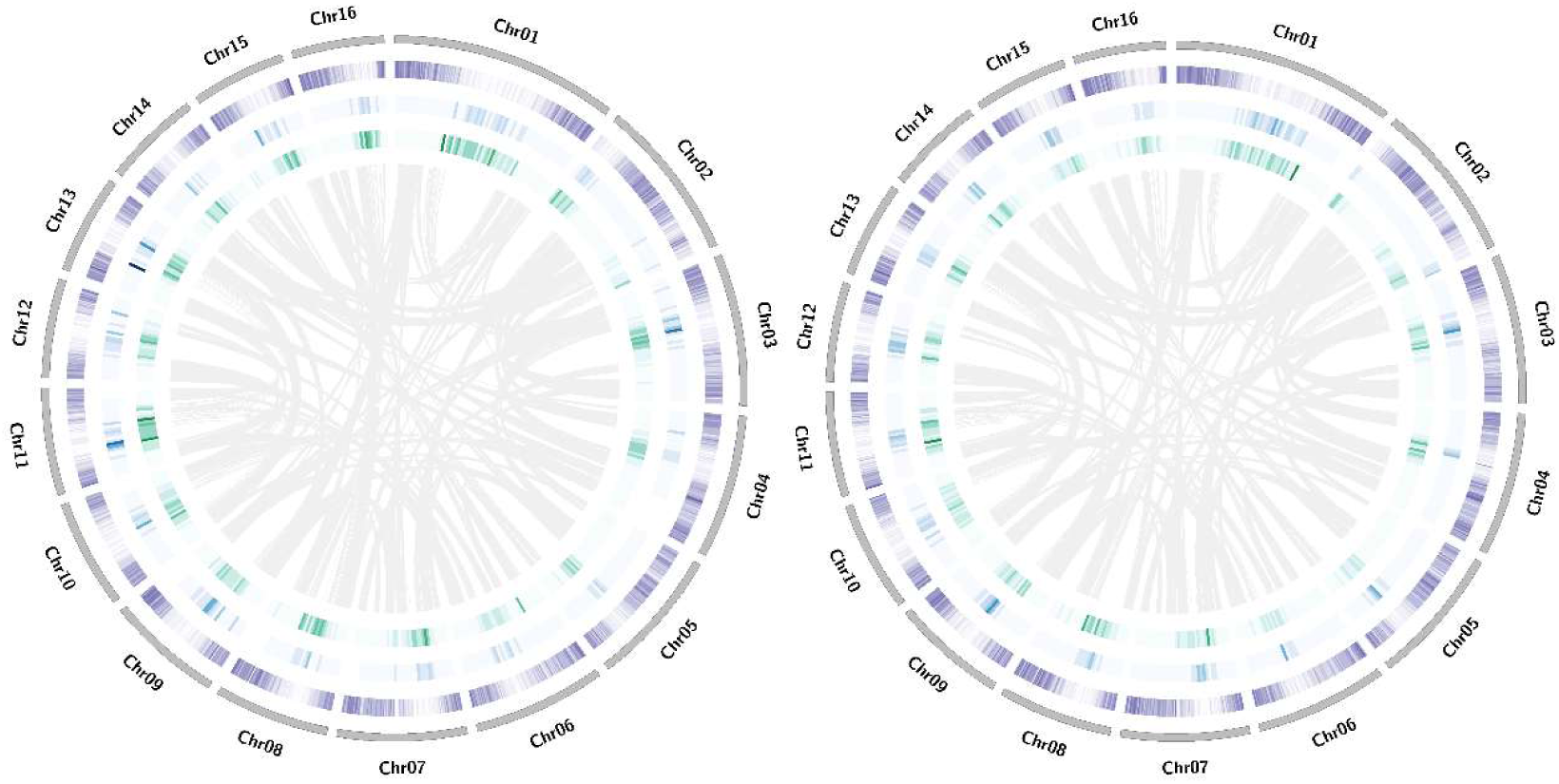
The circos figures of *G. sutherlandii* (left panel) and *G. flanaganii* (right panel) using *G. radlkoferi* as a reference, to do the genome-guided scaffolding.

**Figure S3.**
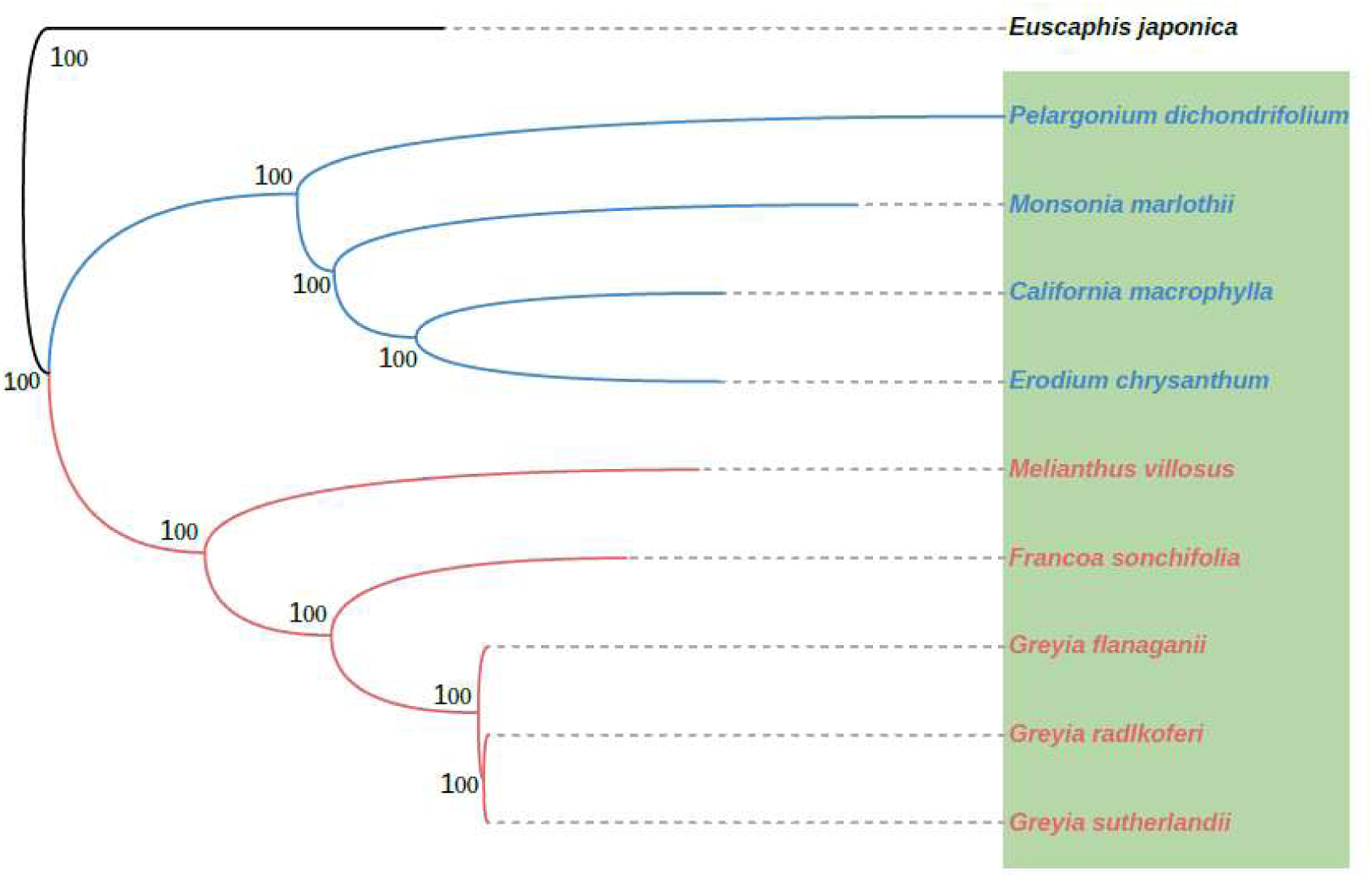
The phylogenetic relationships within Geraniales using *Euscaphis* as the outgroup. This tree utilized 2,479 orthogroups, with a minimum of 58.3% of species having single-copy genes in any orthogroup. Species in Geraniales are highlighted in the green box. The blue and red lines indicate two major clades: one comprising *Pelargonium dichondrifolium, Monsonia marlothii, California macrophylla, and Erodium chrysanthum*; and the other including *Greyia, Melianthus villosus, and Francoa sonchifolia*. The number on the branch represents the bootstrap values

**Figure S4.**
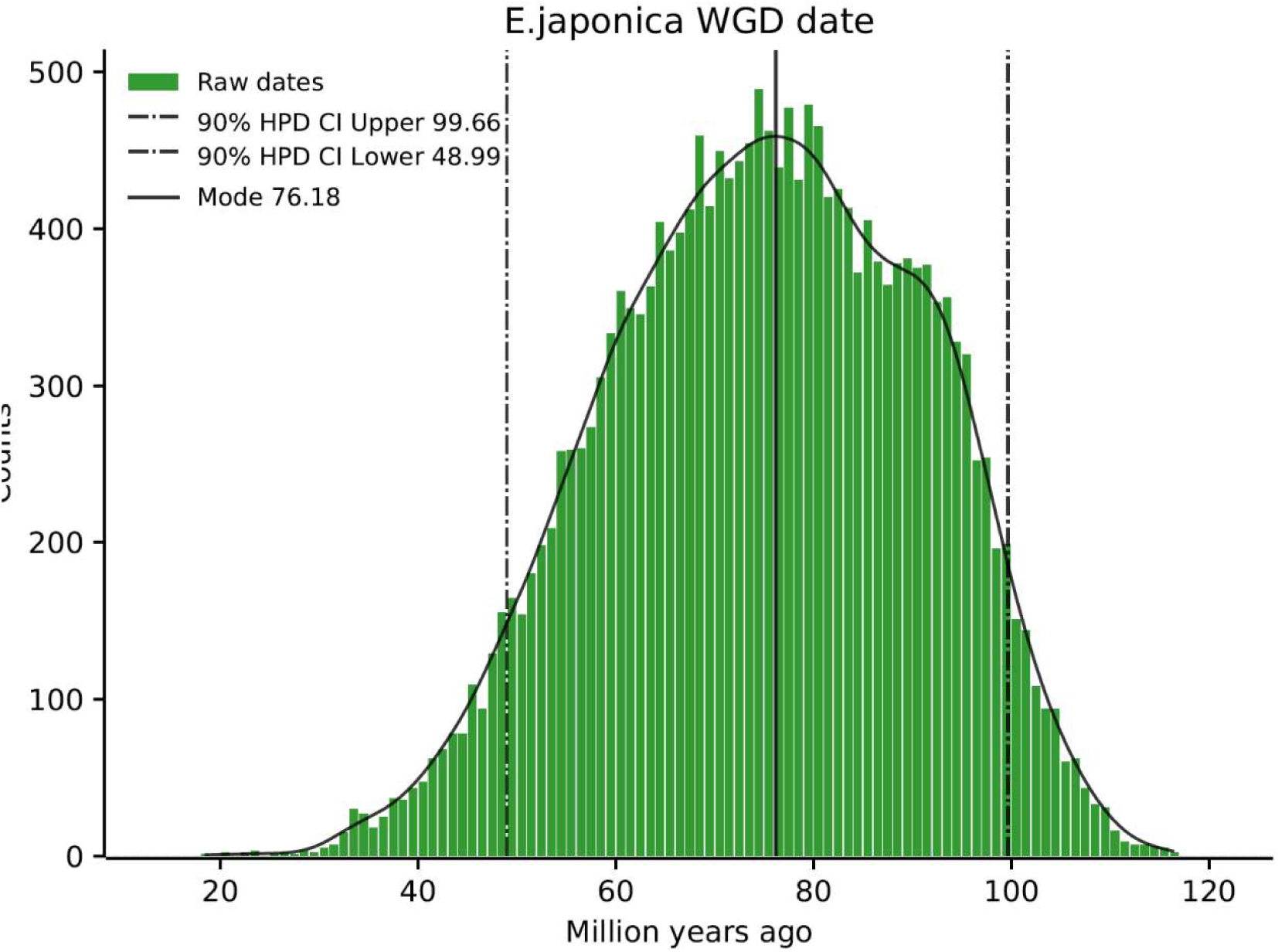
Absolute dating (see Methods) of the *E. japonica* WGD event. The age distribution was obtained by phylogenomic dating of *E. japonica* paralogs. The solid black line represents the Kernel Density Estimation (KDE) of the dated paralogs, while the vertical solid black line represents its peak at 76.18 MYA, which was used as the consensus WGD age estimate. The vertical black dotted lines represent the corresponding 90% confidence interval (CI) for the WGD age estimate, 48.99-99.66 MYA. The green histogram shows the raw distribution of dated paralogs.

**Figure S5.**
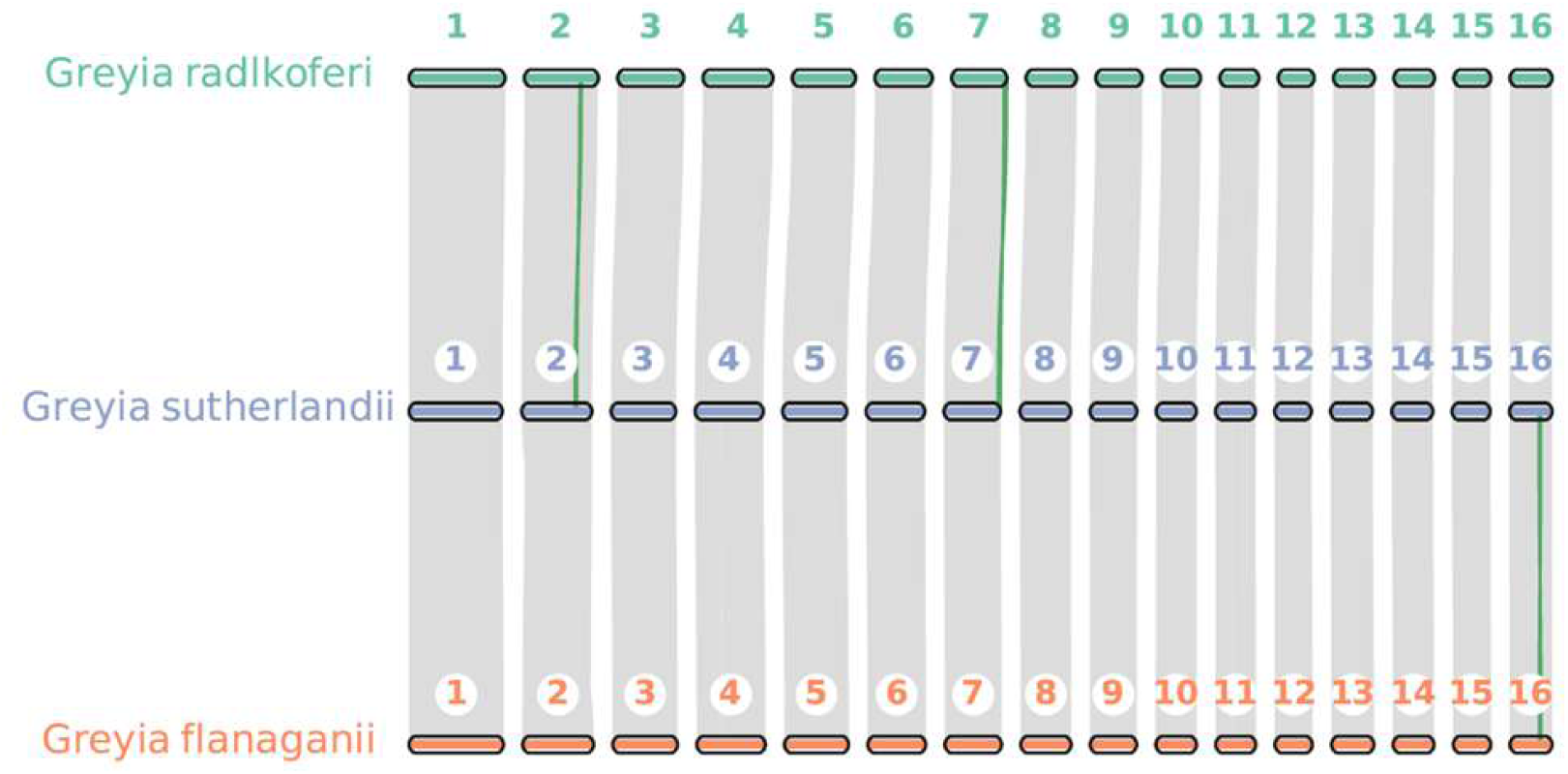
The macrosynteny among *G. radlkoferi*, *G. sutherlandii*, and *G. flanaganii* using MCScanX. The line highlighted in green represent the inversions.

**Figure S6.**
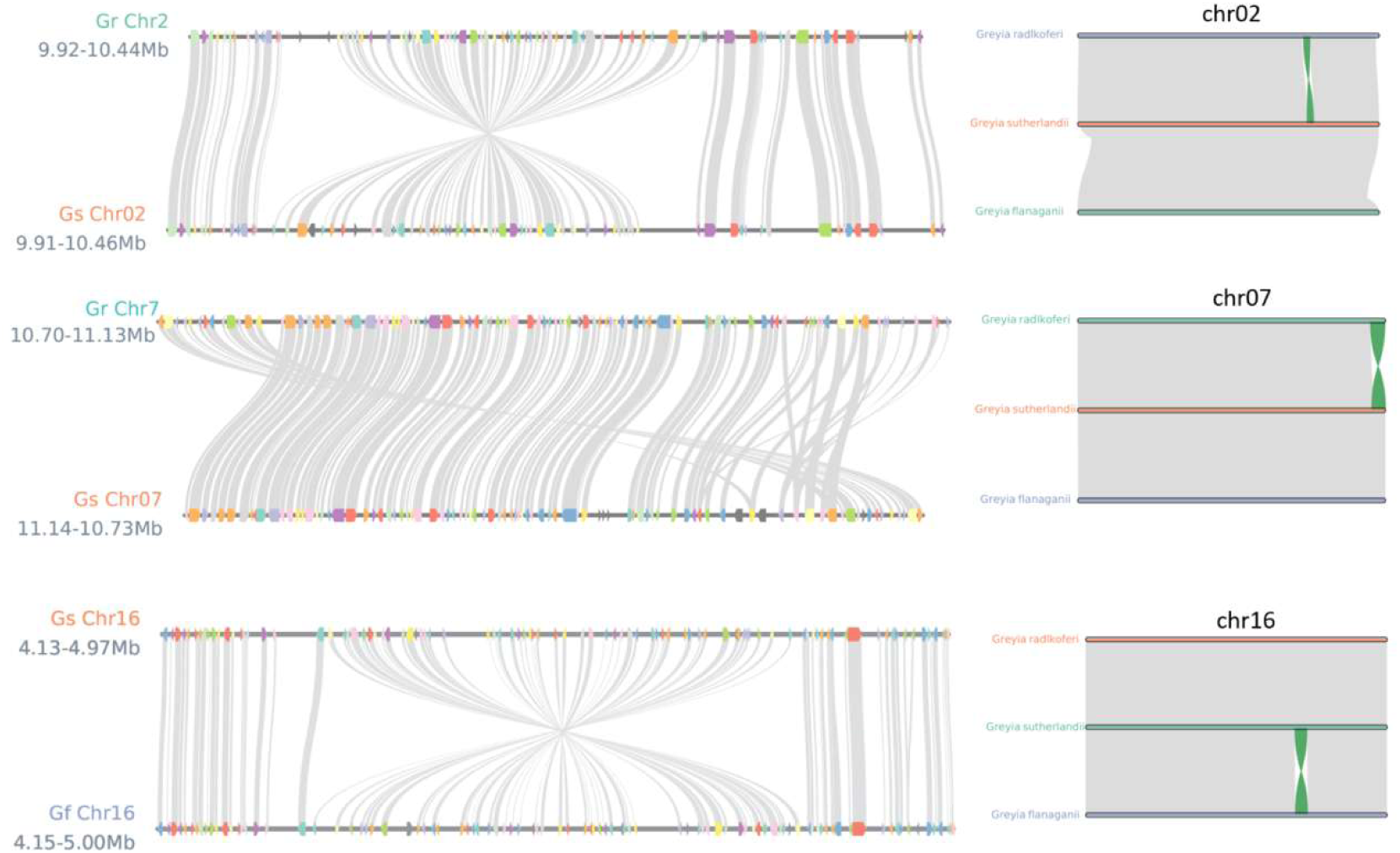
The micro-synteny of three inversions found among *G. radlkoferi*, *G. sutherlandii,* and *G. flanaganii* using MCScanX.

**Figure S7.**
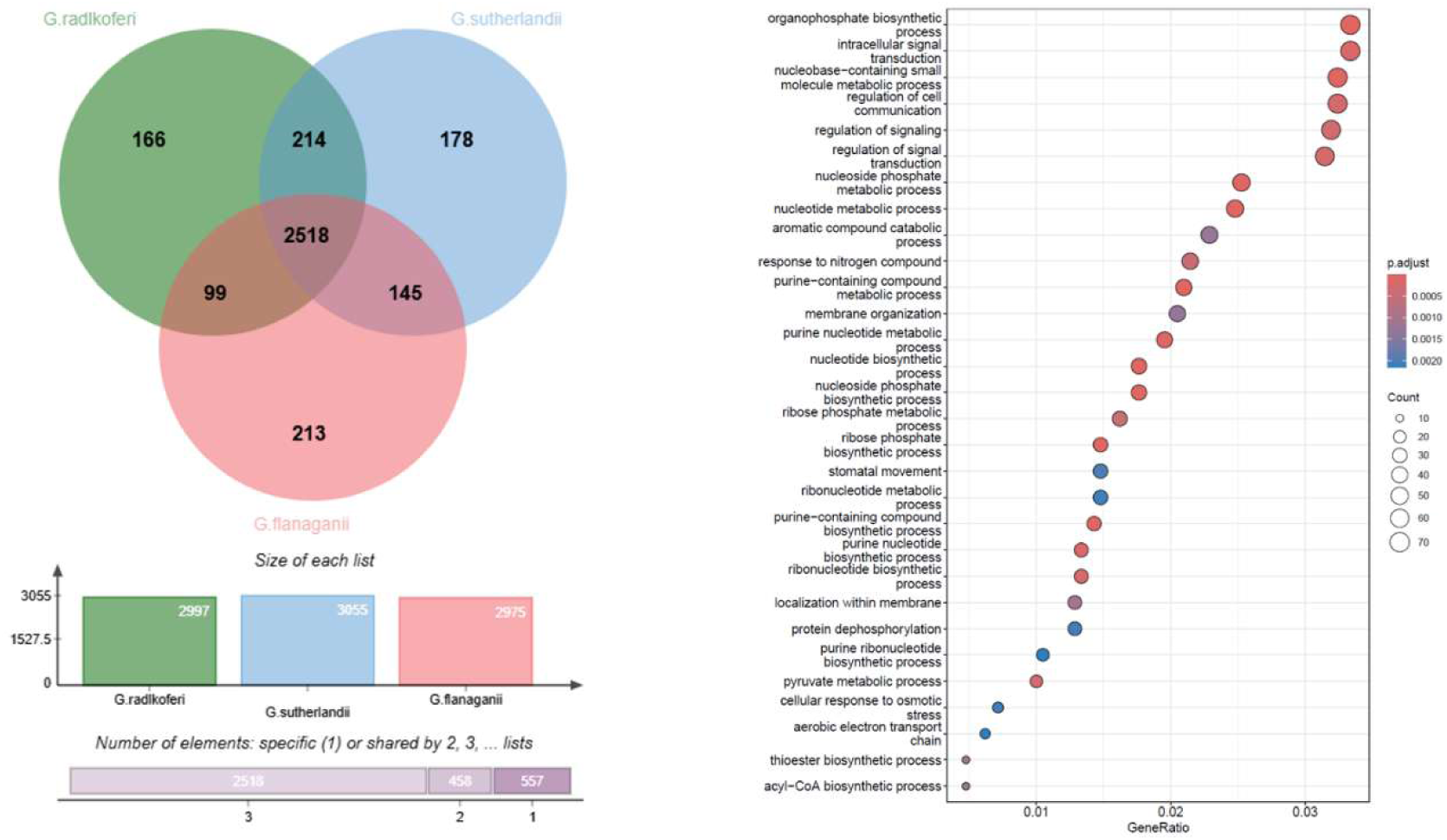
The GO enrichment (p<0.05) of duplicated gene families retained from the WGT event in Greyia. The left panel showed that 2518 families are common in three Greyia species. The right panel is GO enrichment of shared WGT-derived duplicated genes in Greyia (2518 gene families).

**Figure S8.**
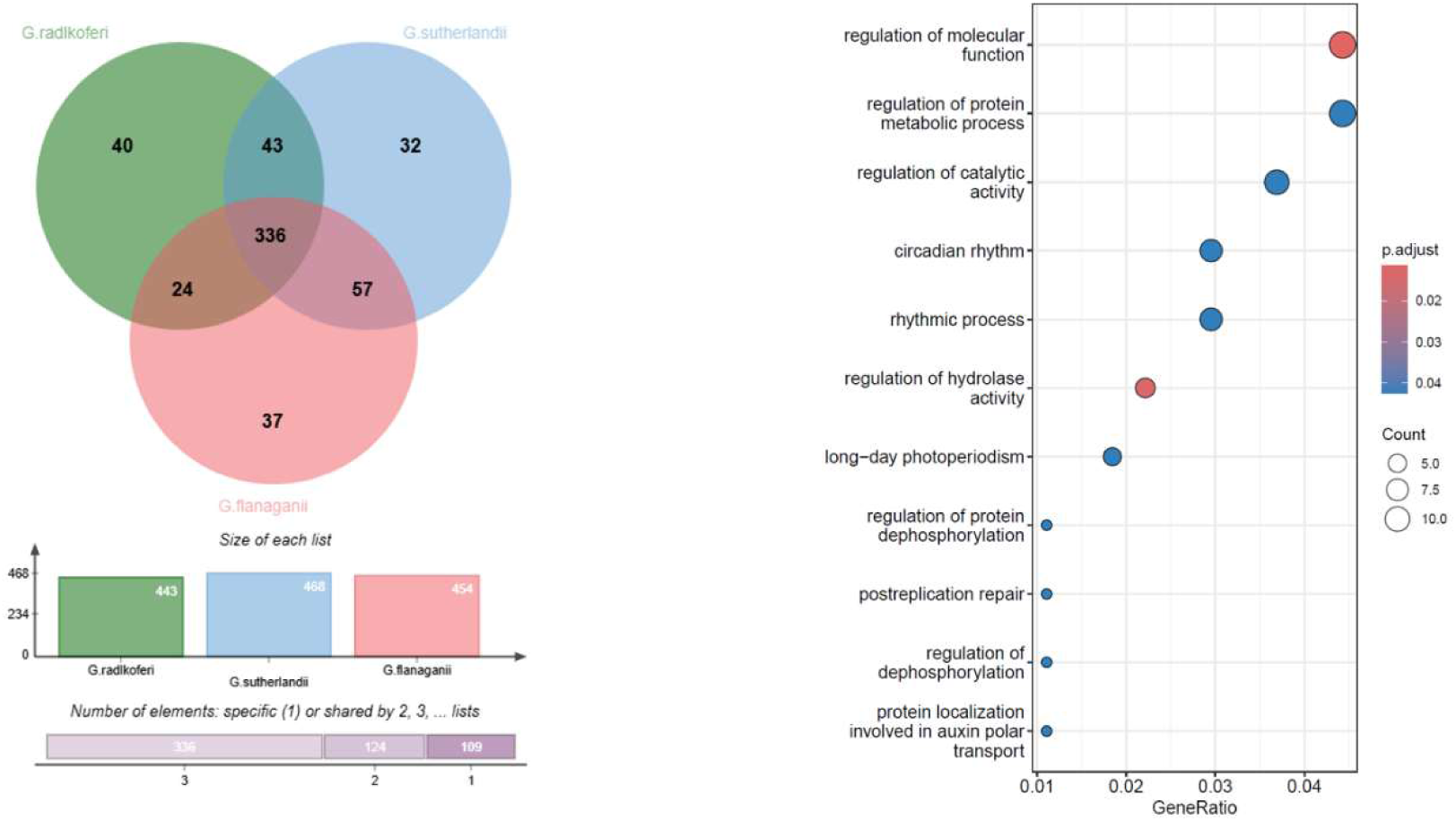
The GO enrichment (p<0.05) of triplicated gene families retained from the WGT event in Greyia. The left panel showed that 1080 genes (360 triple-copy paleoparalogs) consistently retained across all Greyia species belonged to 336 families. The right panel is GO enrichment of shared WGT-derived triplicated genes (three copies) in Greyia (336 gene families).

**Figure S9.**
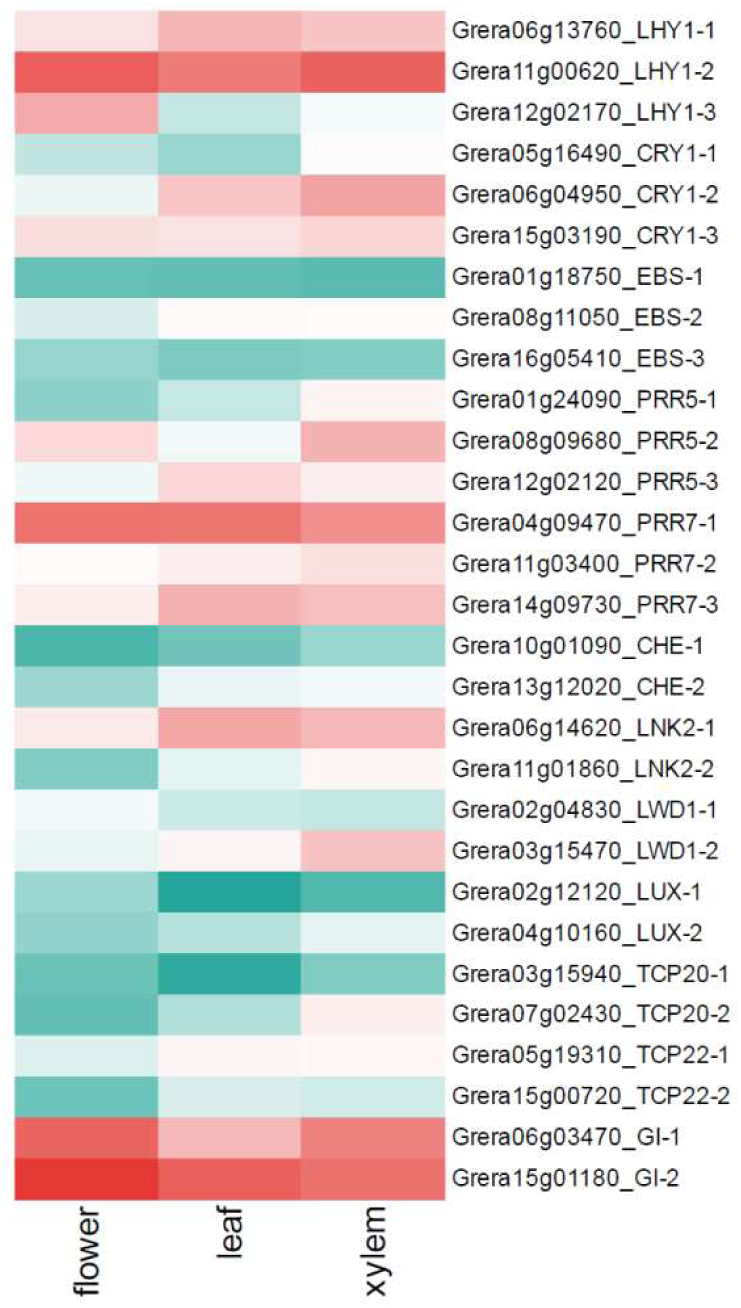
The expression profile of duplicated and triplicated genes involved in photoperiodic module.

**Figure S10.**
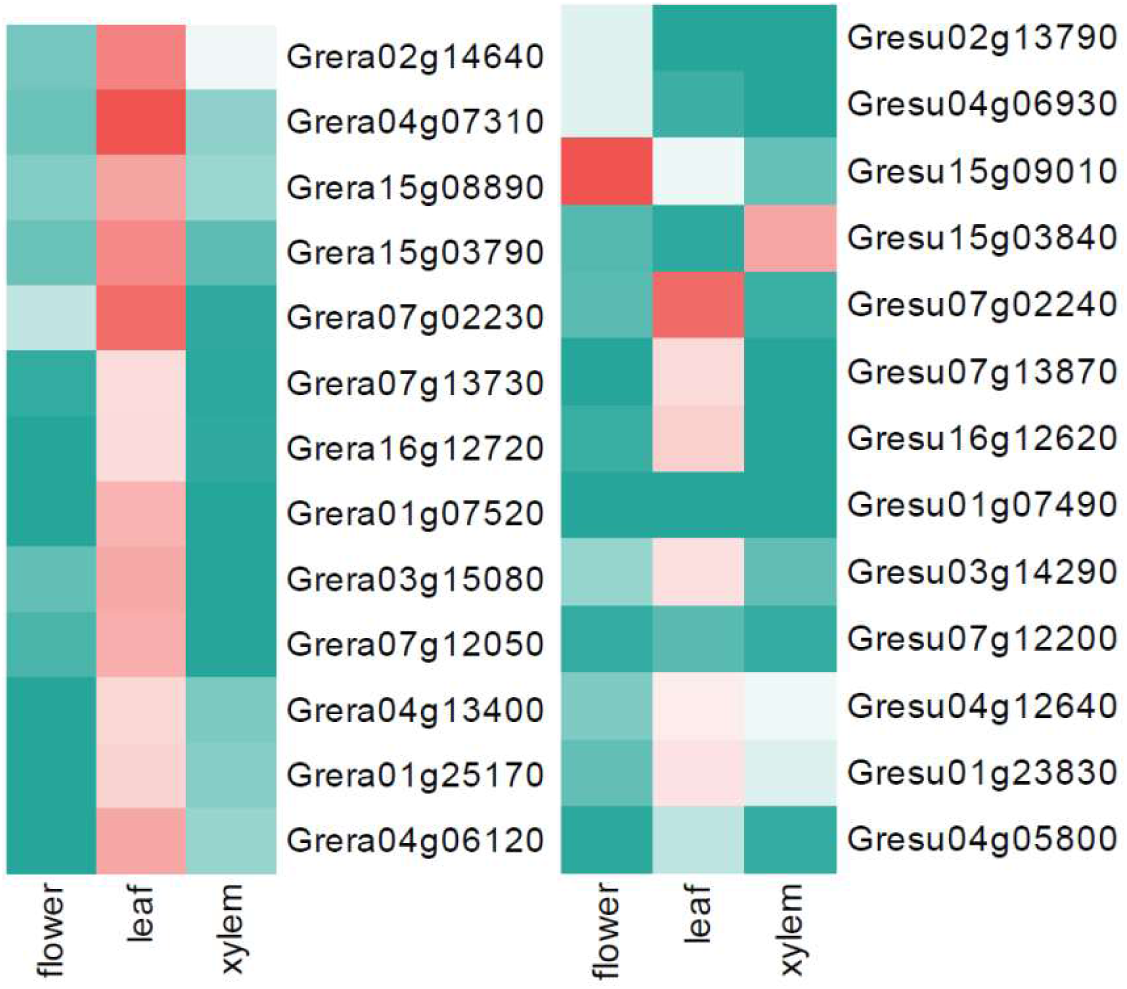
13 MYB transcription factors with significantly higher expression in leaves than in flowers and xylem tissues of *G. radlkoferi,* together with their homologs in *G. sutherlandii*. Each row represents the homologous genes in two Greyia species. Among these genes, Grera0712050 with leaf-specific expression and Gresu07g12200 with no detectable expression across all tissues are the homologs of GL1 in *G. radlkoferi* and *G. sutherlandii,* respectively.

**Figure S11.**
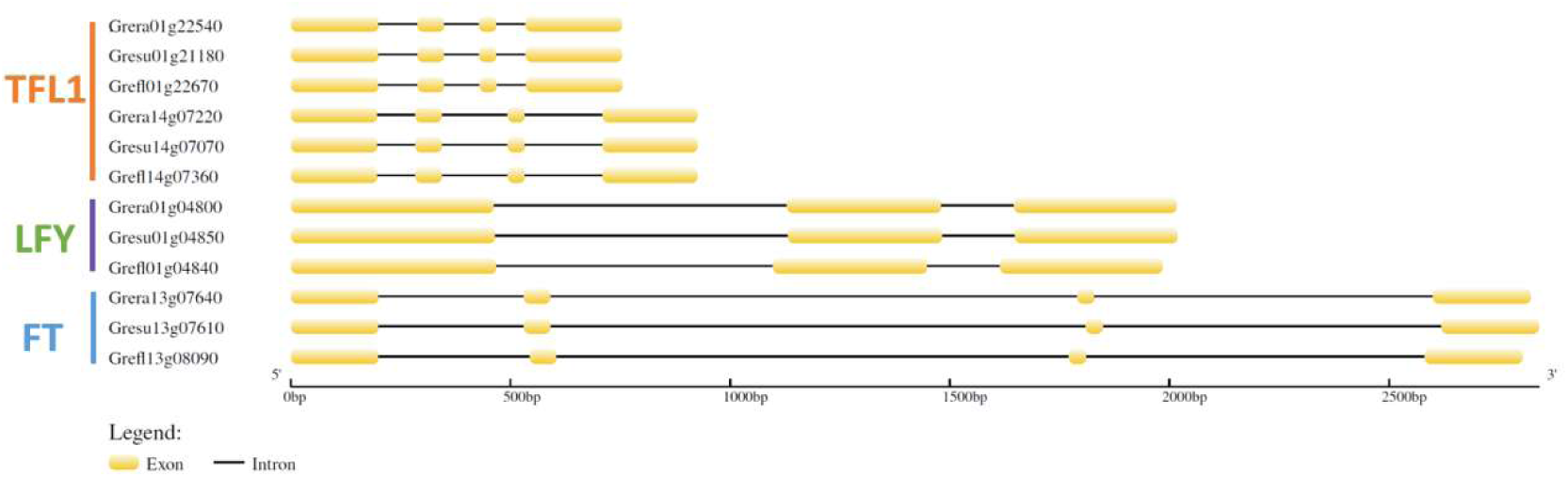
The gene structures and domain regions of *TFL1*, *LFY*, and *FT* in three Greyia.

**Figure S12.**
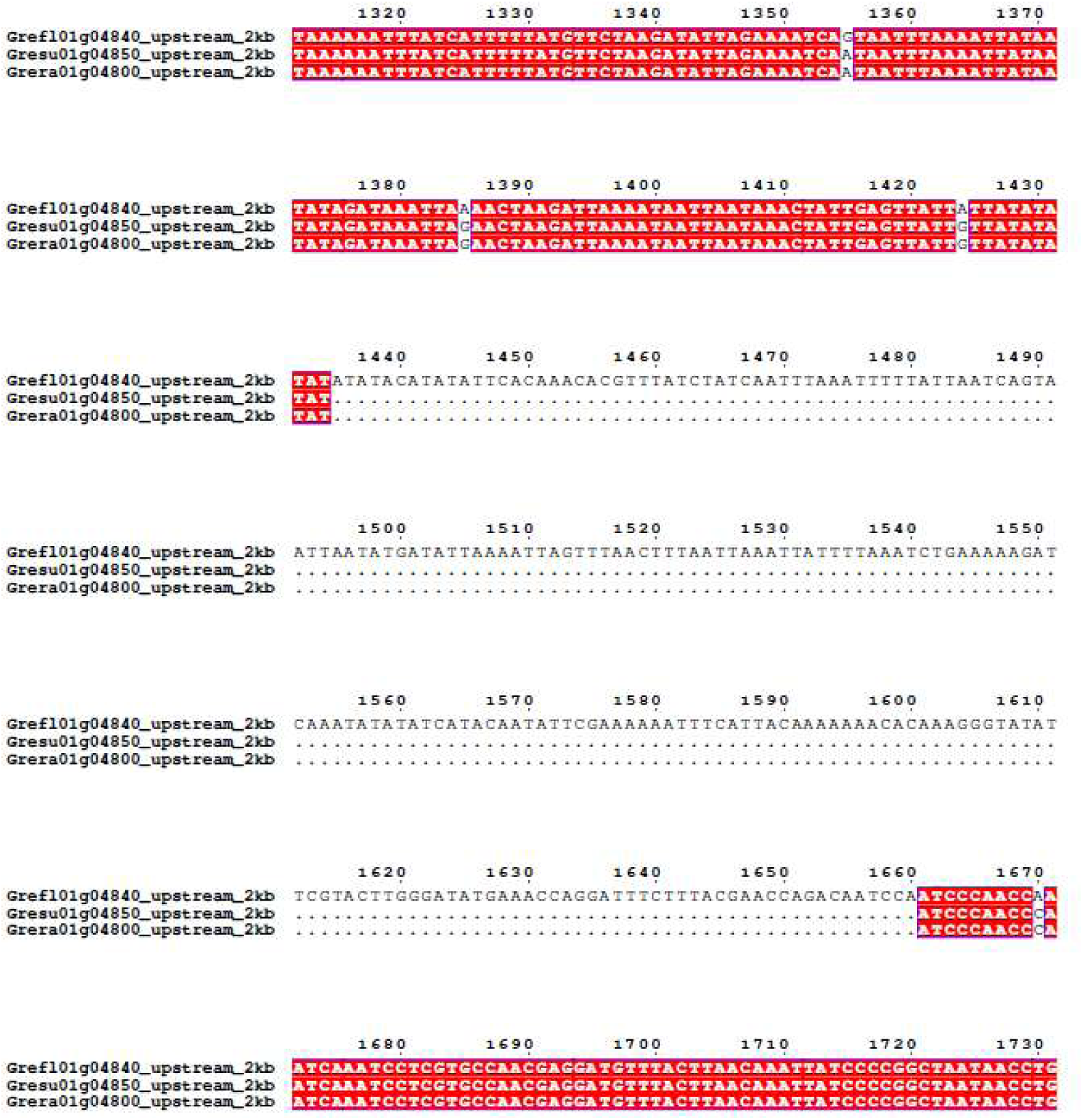
A 226-bp insertion was identified in the LFY promoter of *G. flanaganii*. Multiple sequence alignment of the 2-kb upstream regions from the transcription start site (TSS) revealed that this insertion is located between 1335 and 1660 bp in the alignment, corresponding to 341–566 bp upstream of the TSS. This insertion is absent in *G. radlkoferi* and *G. sutherlandii*.

**Figure S13.**
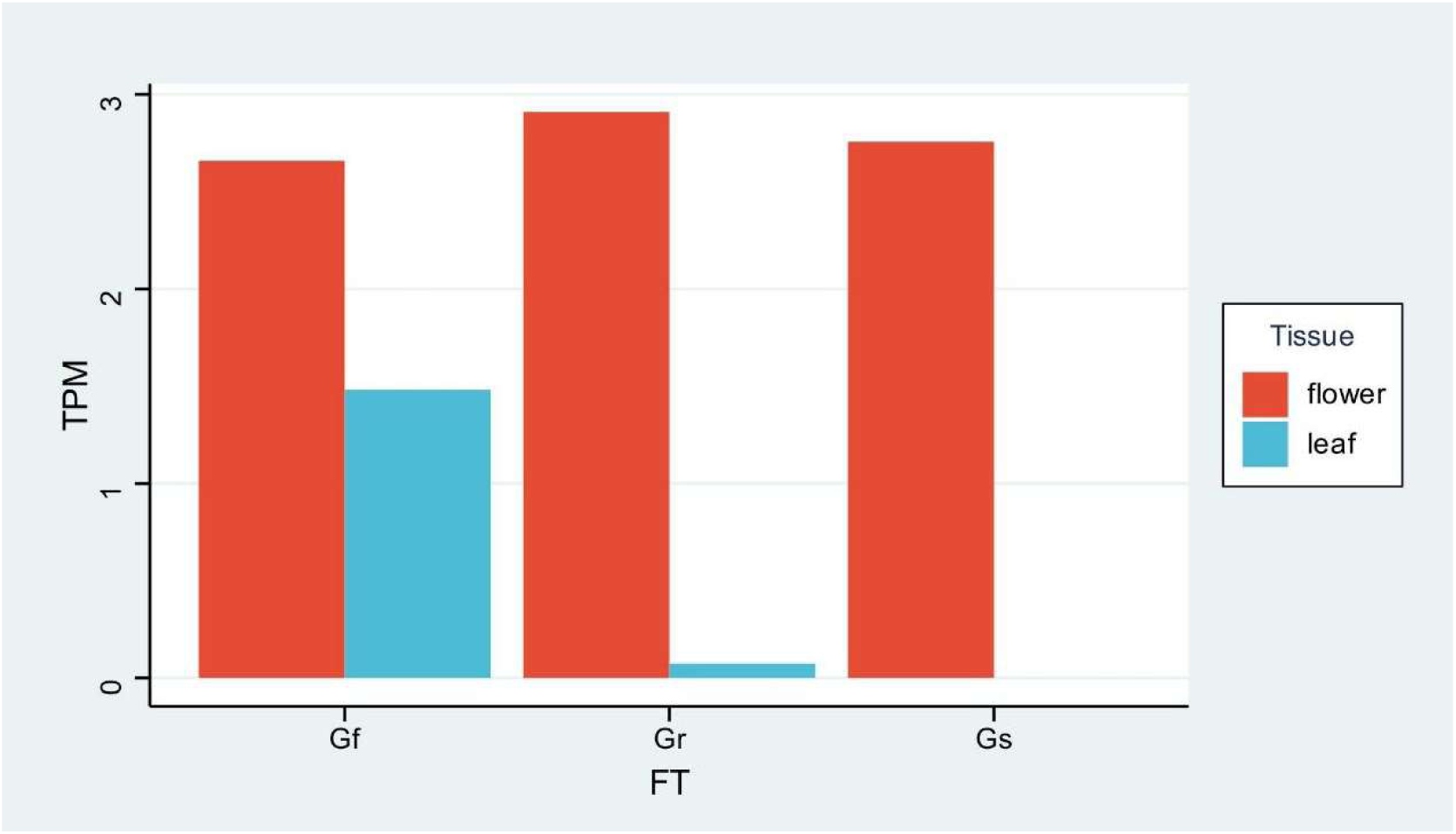
Comparison of FT expression levels in flower and leaf of *G. radlkoferi*, *G. sutherlandii*, and *G. flanaganii*.

**Figure S14.**
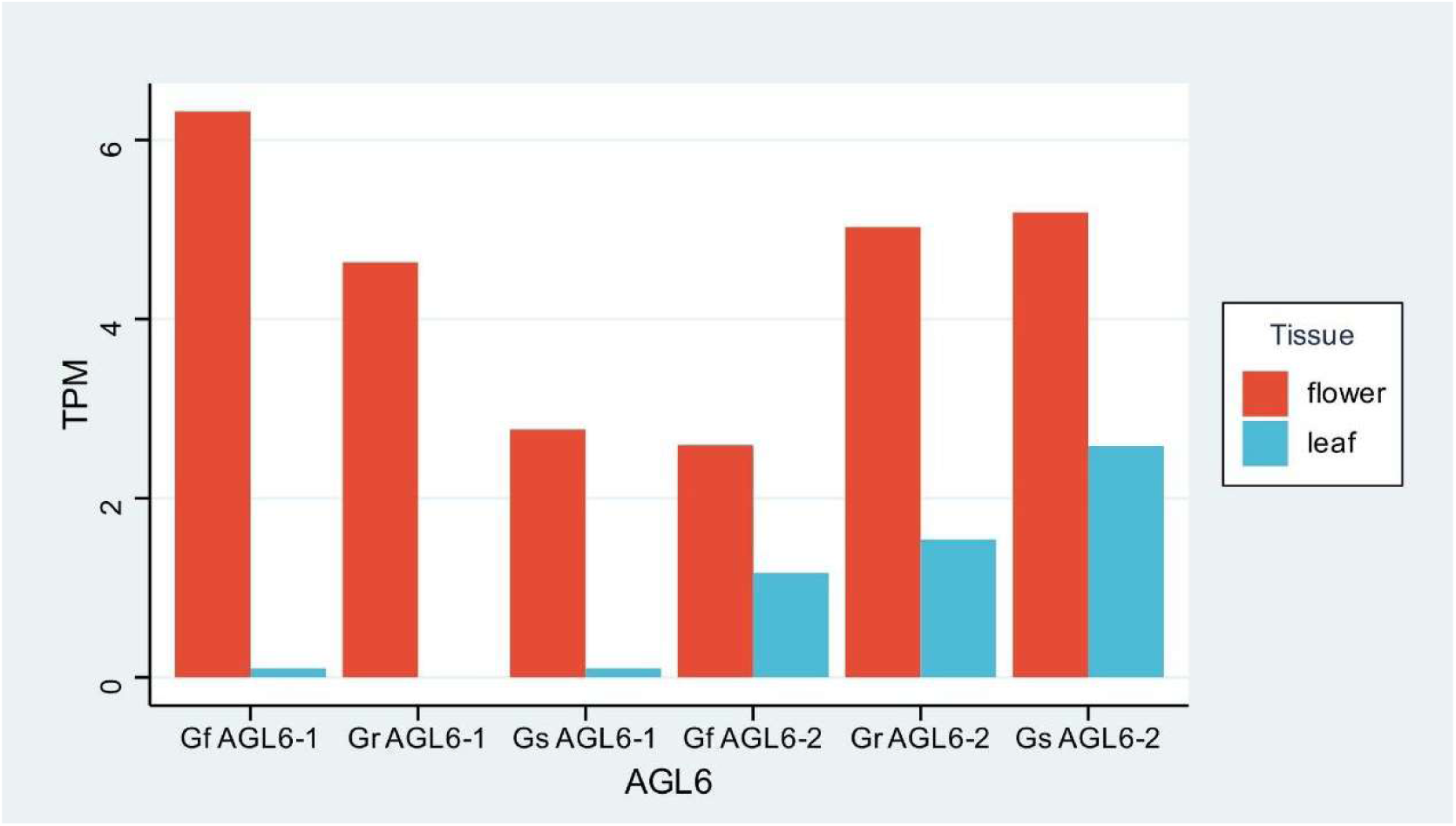
Comparison of AGL6 expression levels in flower and leaf of *G. radlkoferi*, *G. sutherlandii*, and *G. flanaganii*.

**Table S1.**
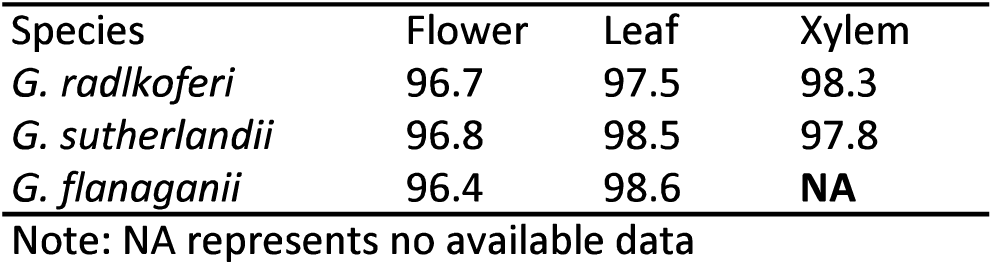
Mapping rates of RNA-seq reads from multiple tissues.

**Table S2.**
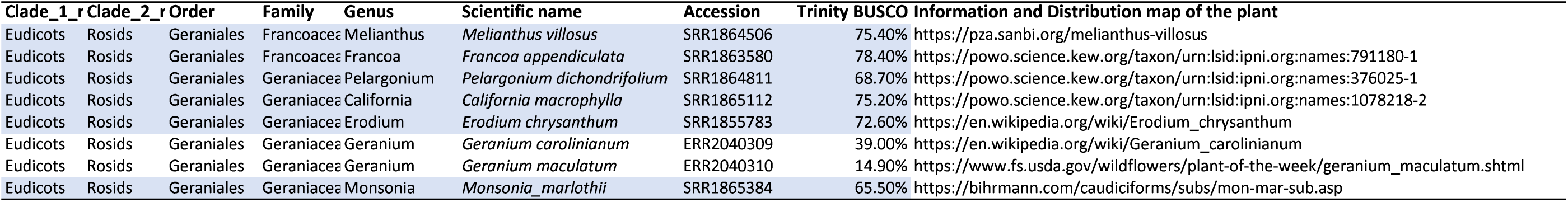
Transcriptomic data for species within Geraniales. Species highlighted in the blue box were included in our study, as their BUSCO completeness scores exceeded 65%.

**Table S3.** Retention of duplicated genes in three *Greyia* spp.

|  | *G. radlkoferi* | *G. sutherlandii* | *G. flanaganii* |
| --- | --- | --- | --- |
| Number of collinear genes | 11250 | 11070 | 11010 |
| Percentage of collinear genes | 41.33% | 41.19% | 39.52% |
| Number of syntenic blocks | 525 | 492 | 509 |
| Pair number from the Greyia-specific WGT | 3442 | 3566 | 3448 |
| Gene numbers from the Greyia-specific WGT | 5910 | 6057 | 5892 |
| Percentage of genes from the Greyia-specific | 52.50% | 54.70% | 53.50% |
| number of orthogroups with duplicated genes | 2997 | 3055 | 2975 |
| Triplicated copies retained from the WGT event | 1097 | 1201 | 1139 |
| number of orthogroups with triplicated genes | 443 | 468 | 454 |

**Table S4.**
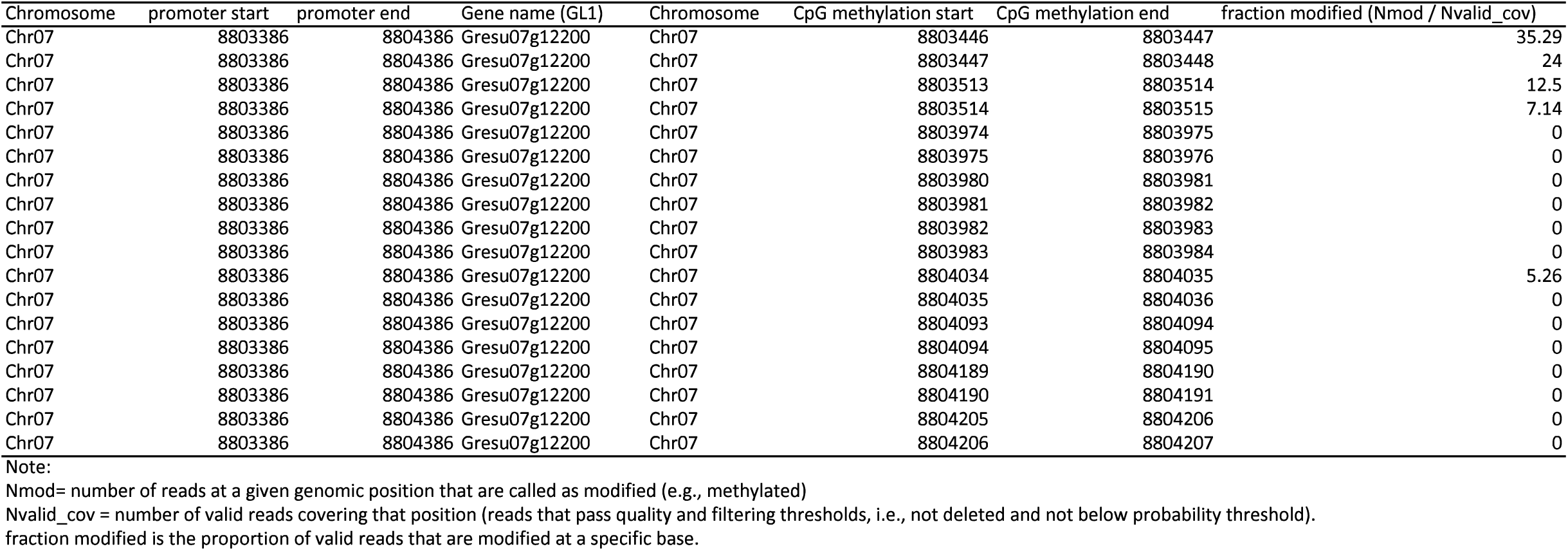
Methylated sites detected in the promoter of *GL1* in *G. sutherlandii*.

| Chromosome | promoter start | promoter end | Gene name (GL1) | Chromosome | CpG methylation start | CpG methylation end | fraction modified (Nmod / Nvalid_cov) |
| --- | --- | --- | --- | --- | --- | --- | --- |
| Chr07 | 8803386 | 8804386 | Gresu07g12200 | Chr07 | 8803446 | 8803447 | 35.29 |
| Chr07 | 8803386 | 8804386 | Gresu07g12200 | Chr07 | 8803447 | 8803448 | 24 |
| Chr07 | 8803386 | 8804386 | Gresu07g12200 | Chr07 | 8803513 | 8803514 | 12.5 |
| Chr07 | 8803386 | 8804386 | Gresu07g12200 | Chr07 | 8803514 | 8803515 | 7.14 |
| Chr07 | 8803386 | 8804386 | Gresu07g12200 | Chr07 | 8803974 | 8803975 | 0 |
| Chr07 | 8803386 | 8804386 | Gresu07g12200 | Chr07 | 8803975 | 8803976 | 0 |
| Chr07 | 8803386 | 8804386 | Gresu07g12200 | Chr07 | 8803980 | 8803981 | 0 |
| Chr07 | 8803386 | 8804386 | Gresu07g12200 | Chr07 | 8803981 | 8803982 | 0 |
| Chr07 | 8803386 | 8804386 | Gresu07g12200 | Chr07 | 8803982 | 8803983 | 0 |
| Chr07 | 8803386 | 8804386 | Gresu07g12200 | Chr07 | 8803983 | 8803984 | 0 |
| Chr07 | 8803386 | 8804386 | Gresu07g12200 | Chr07 | 8804034 | 8804035 | 5.26 |
| Chr07 | 8803386 | 8804386 | Gresu07g12200 | Chr07 | 8804035 | 8804036 | 0 |
| Chr07 | 8803386 | 8804386 | Gresu07g12200 | Chr07 | 8804093 | 8804094 | 0 |
| Chr07 | 8803386 | 8804386 | Gresu07g12200 | Chr07 | 8804094 | 8804095 | 0 |
| Chr07 | 8803386 | 8804386 | Gresu07g12200 | Chr07 | 8804189 | 8804190 | 0 |
| Chr07 | 8803386 | 8804386 | Gresu07g12200 | Chr07 | 8804190 | 8804191 | 0 |
| Chr07 | 8803386 | 8804386 | Gresu07g12200 | Chr07 | 8804205 | 8804206 | 0 |
| Chr07 | 8803386 | 8804386 | Gresu07g12200 | Chr07 | 8804206 | 8804207 | 0 |

Note: Nmod= number of reads at a given genomic position that are called as modified (e.g., methylated) Nvalid\_cov = number of valid reads covering that position (reads that pass quality and filtering thresholds, i.e., not deleted and not below probability threshold). fraction modified is the proportion of valid reads that are modified at a specific base.

**Table SS.**
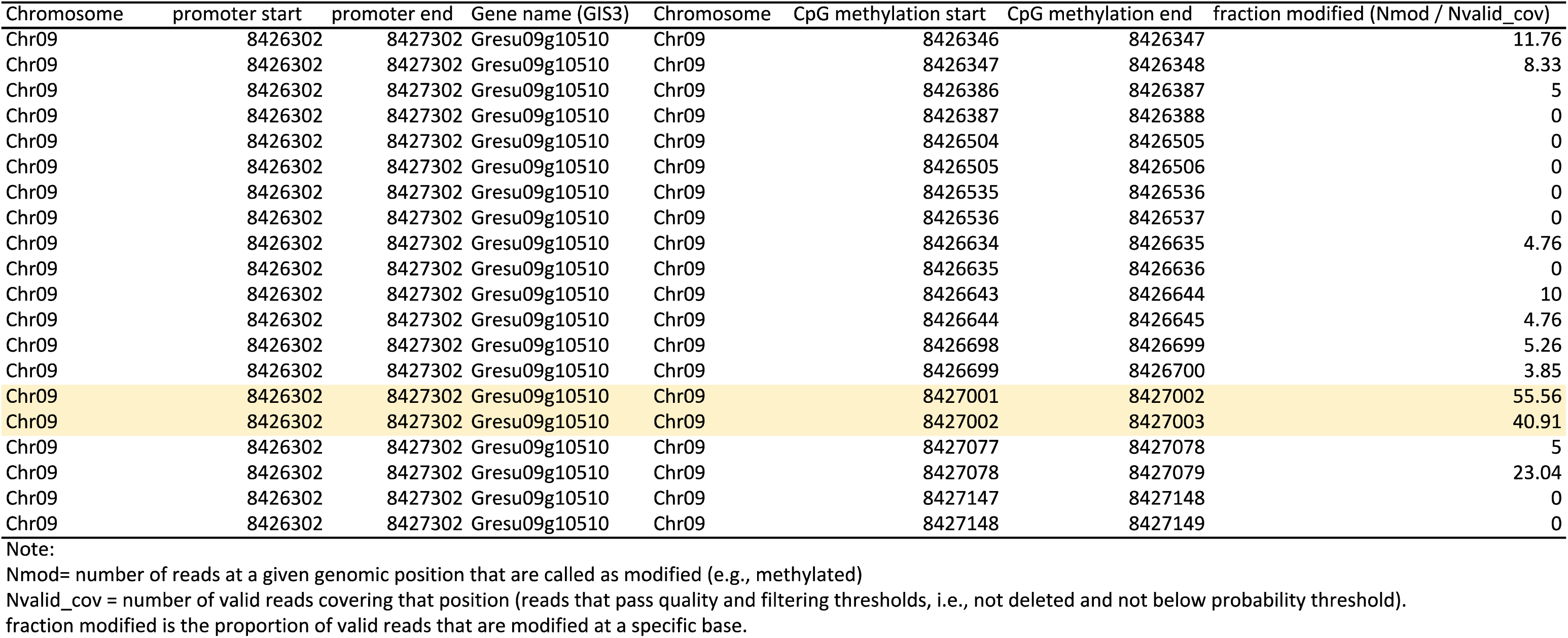
Methylated sites detected in the promoter of *G/S3* in *G. sutherlandii*.

**Table S6a.** Type II MADS-box genes and encoded proteins.

| Genename | Gr | protein length (AA) | Gs | protein length (AA) | Gf | protein length (AA) |
| --- | --- | --- | --- | --- | --- | --- |
| MICK* | Grera09g12030 | 370 | Gresu09g11880 | 371 | Grefl09g12270 | 370 |
| MICK* | Grera06g04730 | 214 | Gresu06g04740 | 211 | Grefl06g04780 | 208 |
| MICK* | Grera10g03580 | 330 | Gresu10g03660 | 331 | Grefl10g03620 | 330 |
| MICK* | Grera05g14630 | 307 | Gresu05g14530 | 308 | Grefl05g15060 | 314 |
| SEPgene-1 | Grera03g01070 | 239 | Gresu03g01030 | 240 | Grefl03g01060 | 239 |
| SEPgene-2 | Grera05g08620 | 242 | Gresu05g08500 | 243 | Grefl05g08850 | 242 |
| SEPgene-3 | Grera09g14110 | 241 | Gresu09g13910 | 242 | Grefl09g14330 | 241 |
| SEPgene-4 | Grera02g02950 | 240 | Gresu02g02580 | 241 | Grefl02g01720 | 240 |
| SEPgene-5 | Grera07g01690 | 244 | Gresu07g01700 | 245 | Grefl07g01790 | 244 |
| SEPgene-6 | Grera07g10570 | 243 | Gresu07g10680 | 244 | Grefl07g11210 | 243 |
| AP1-1 | Grera09g11630 | 242 | Gresu09g11470 | 243 | Grefl09g11860 | 242 |
| AP1-2 | Grera04g13150 | 233 | Gresu04g12360 | 234 | Grefl04g13100 | 233 |
| AP1-3 | Grera07g10560 | 243 | Gresu07g10670 | 244 | Grefl07g11200 | 243 |
| AP1-4 | Grera02g02960 | 241 | Gresu02g02590 | 245 | Grefl02g01730 | 244 |
| AP1-5 | Grera07g01680 | 247 | Gresu07g01690 | 248 | Grefl07g01780 | 247 |
| FLC-1 | Grera03g01080 | 241 | Gresu03g01040 | 242 | Grefl03g01070 | 242 |
| FLC-2 | Grera03g01090 | 186 | Gresu03g01050 | 187 | Grefl03g01080 | 186 |
| FLC-3 | Grera09g14100 | 187 | Gresu09g13900 | 188 | Grefl09g14320 | 187 |
| FLC-4 | Grera07g02500 | 207 | Gresu07g02500 | 207 | Grefl07g02610 | 207 |
| AGL6-1 | Grera12g00770 | 244 | Gresu12g00780 | 245 | Grefl12g00820 | 244 |
| AGL6-2 | Grera01g22290 | 231 | Gresu01g20930 | 260 | Grefl01g22410 | 208 |
| AGL12 | Grera05g19520 | 202 | Gresu05g19440 | 202 | Grefl05g20230 | 202 |
| AG-1 | Grera02g21200 | 251 | Gresu02g19920 | 254 | Grefl02g19830 | 247 |
| AG-2 | Grera08g13480 | 269 | Gresu08g13150 | 270 | Grefl08g13490 | 250 |
| AG-3 | Grera05g19500 | 235 | Gresu05g19420 | 236 | Grefl05g20210 | 235 |
| AG-4 | Grera01g04340 | 191 | Gresu01g04380 | 192 | Grefl01g04370 | 193 |
| AGL17-1 | Grera05g13210 | 226 | Gresu05g13110 | 227 | Grefl05g13590 | 226 |
| AGL17-2 | Grera06g02440 | 292 | Gresu06g02400 | 303 | Grefl06g02430 | 293 |
| AGL17-3 | Grera04g18370 | 272 | Gresu04g17410 | 273 | Grefl04g18330 | 272 |
| SOC1-1 | Grera11g11710 | 218 | Gresu11g11690 | 219 | Grefl11g12230 | 218 |
| SOC1-2 | Grera15g09700 | 210 | Gresu15g09800 | 211 | Grefl15g10080 | 222 |
| SOC1-3 | Grera16g01760 | 205 | Gresu16g01760 | 269 | Grefl16g01800 | 215 |
| SVP-1 | Grera10g02960 | 226 | Gresu10g03030 | 227 | Grefl10g02990 | 226 |
| SVP-2 | Grera13g09690 | 208 | Gresu13g09710 | 209 | Grefl13g10290 | 222 |
| SVP-3 | Grera13g09720 | 225 | Gresu13g09750 | 226 | Grefl13g10320 | 225 |
| SVP-4 | Grera04g18260 | 235 | Gresu04g17310 | 221 | Grefl04g18220 | 220 |
| SVP-5 | Grera10g08420 | 235 | Gresu10g08550 | 236 | Grefl10g09040 | 222 |
| SVP-6 | Grera06g02670 | 226 | Gresu06g02630 | 222 | Grefl06g02660 | 221 |
| AGL15-1 | Grera06g14500 | 246 | Gresu06g14440 | 247 | Grefl06g14780 | 246 |
| AGL15-2 | Grera11g01570 | 264 | Gresu11g01580 | 257 | Grefl11g01550 | 260 |
| AP3-1 | Grera04g16140 | 224 | Gresu04g15260 | 225 | Grefl04g16110 | 224 |
| AP3-2 | Grera04g16130 | 221 | Gresu06g01600 | 221 | Grefl06g01640 | 221 |
| PI-1 | Grera15g02040 | 199 | Gresu15g02060 | 200 | Grefl15g02050 | 199 |
| PI-2 | Grera15g02050 | 211 | Gresu15g02070 | 212 | Grefl15g02060 | 211 |
| Bsister-1 | Grera03g00170 | 237 | Gresu03g00160 | 238 | Grefl03g00170 | 237 |
| Bsister-2 | Grera08g15540 | 237 | Gresu08g15330 | 234 | Grefl08g15750 | 237 |
| Bsister-3 | Grera05g09850 | 275 | Gresu05g09730 | 236 | Grefl05g10100 | 712 |

Note: the number behind the genes only represent the different copies.

**Table S6b.** Count of Type II MADS-box genes and encoded proteins.

| Category | *G. radlkoferi* | *G. sutherlandii* | *G. flanaganii* | *A. thaliana* |
| --- | --- | --- | --- | --- |
| MIKC* | 4 | 4 | 4 | 6 |
| Bsister | 3 | 3 | 3 | 1 |
| PI | 2 | 2 | 2 | 1 |
| AP3 | 2 | 2 | 2 | 1 |
| AGL17 | 3 | 3 | 3 | 4 |
| AGL15 | 2 | 2 | 2 | 2 |
| SVP | 6 | 6 | 6 | 2 |
| AGL12 | 1 | 1 | 1 | 1 |
| AG | 4 | 4 | 4 | 4 |
| SOC1 | 3 | 3 | 3 | 5 |
| FLC | 4 | 4 | 4 | 5 |
| AGL6 | 2 | 2 | 2 | 2 |
| AP1 | 5 | 5 | 5 | 4 |
| SEP | 6 | 6 | 6 | 4 |
| Total | 47 | 47 | 47 | 42 |

**Table S7.**
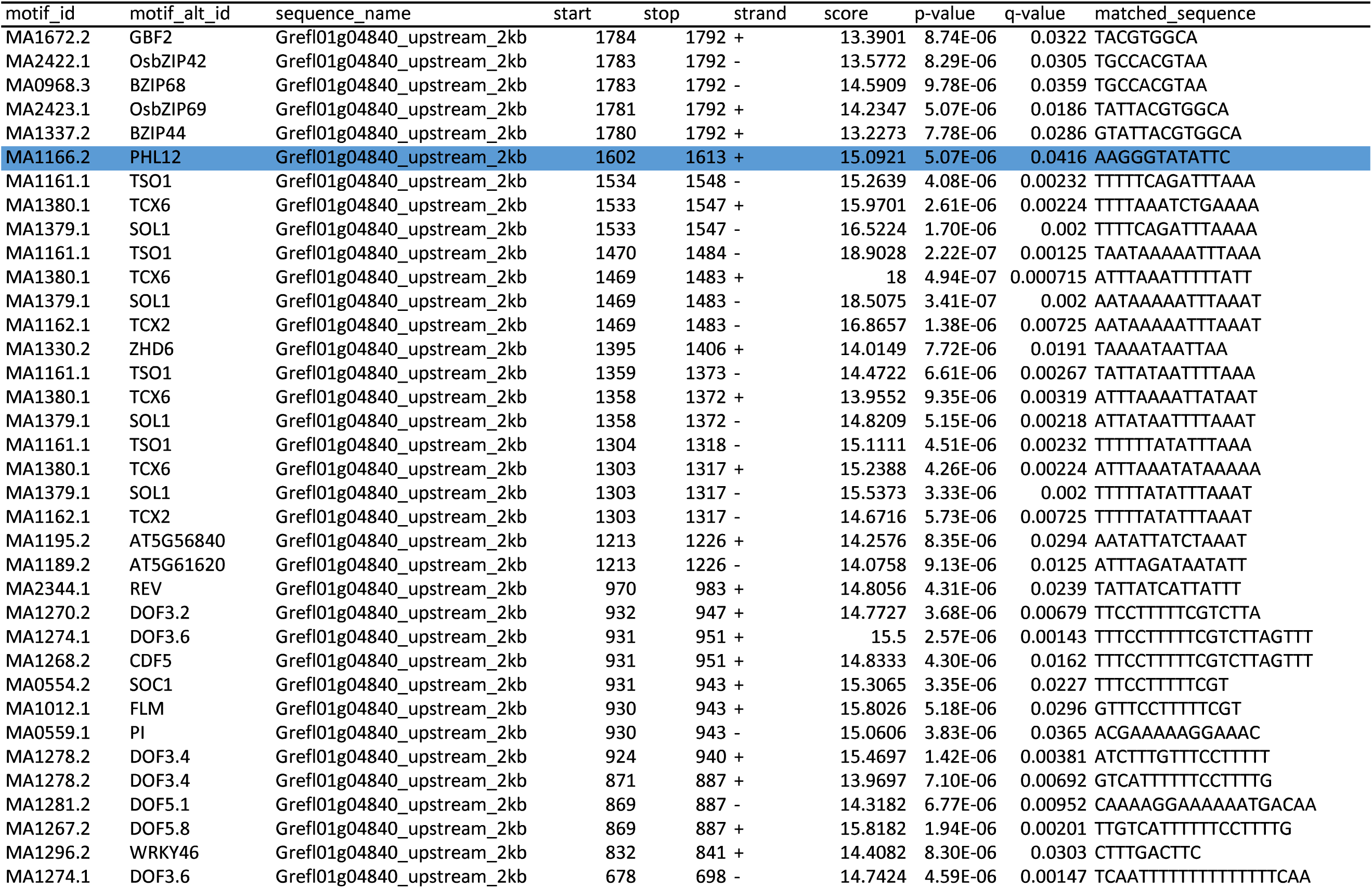
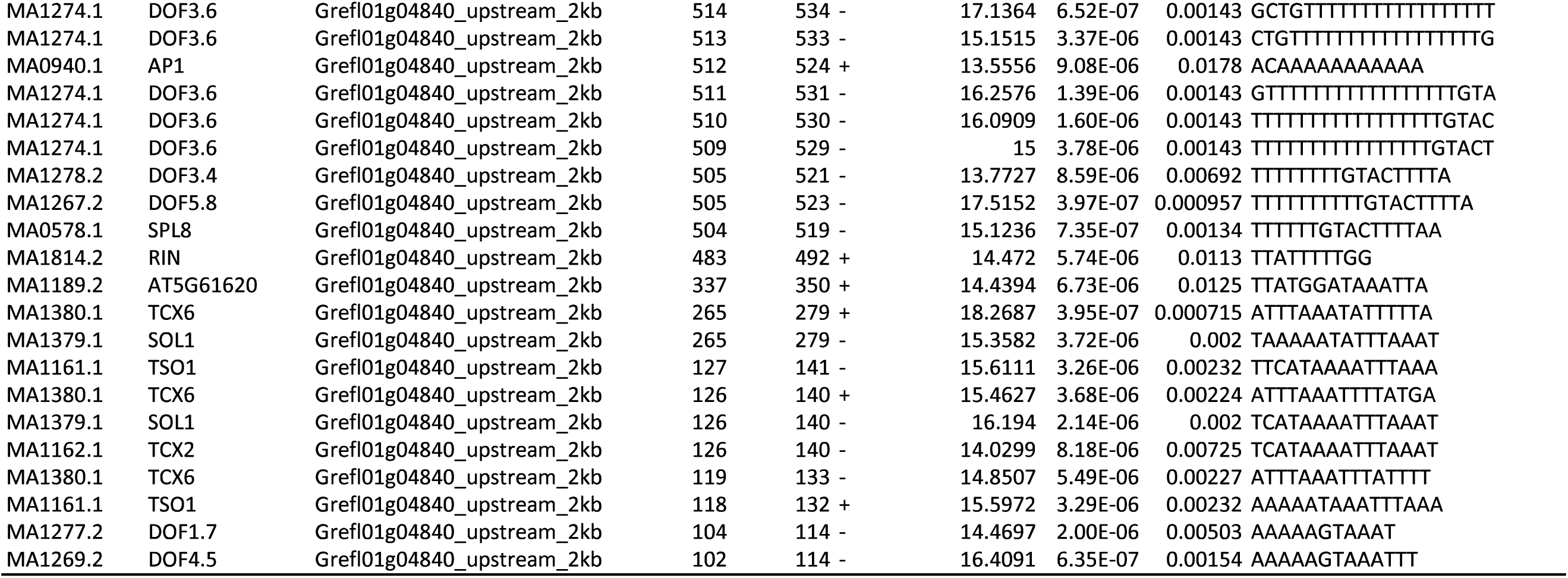
Predicted TFBSs across the 2-kb upstream promoter region of *LFY* in *G. flanaganii* {A of ATG located at position 2000). The PHL12 binding site is highlighted in blue.

**Table S8.**
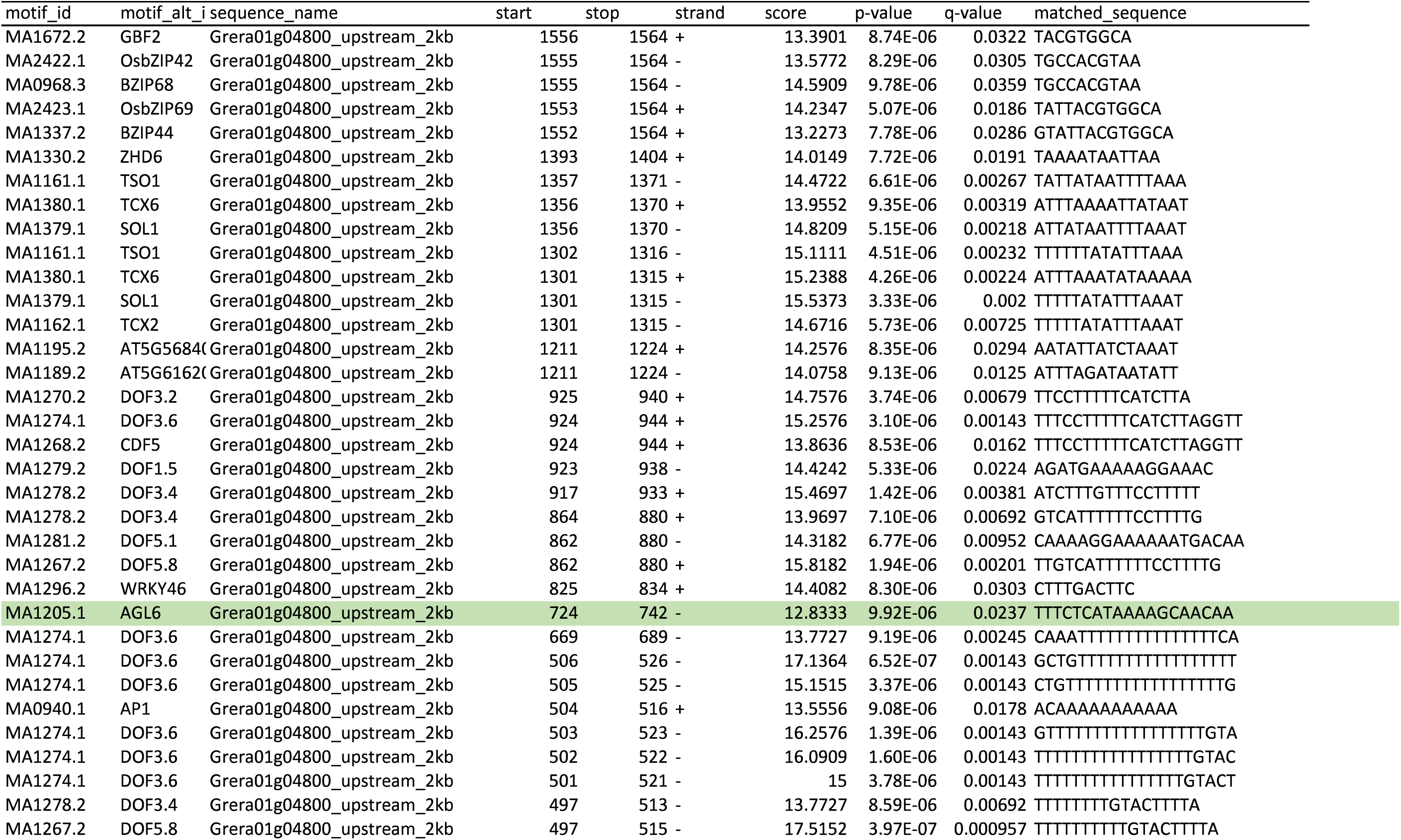
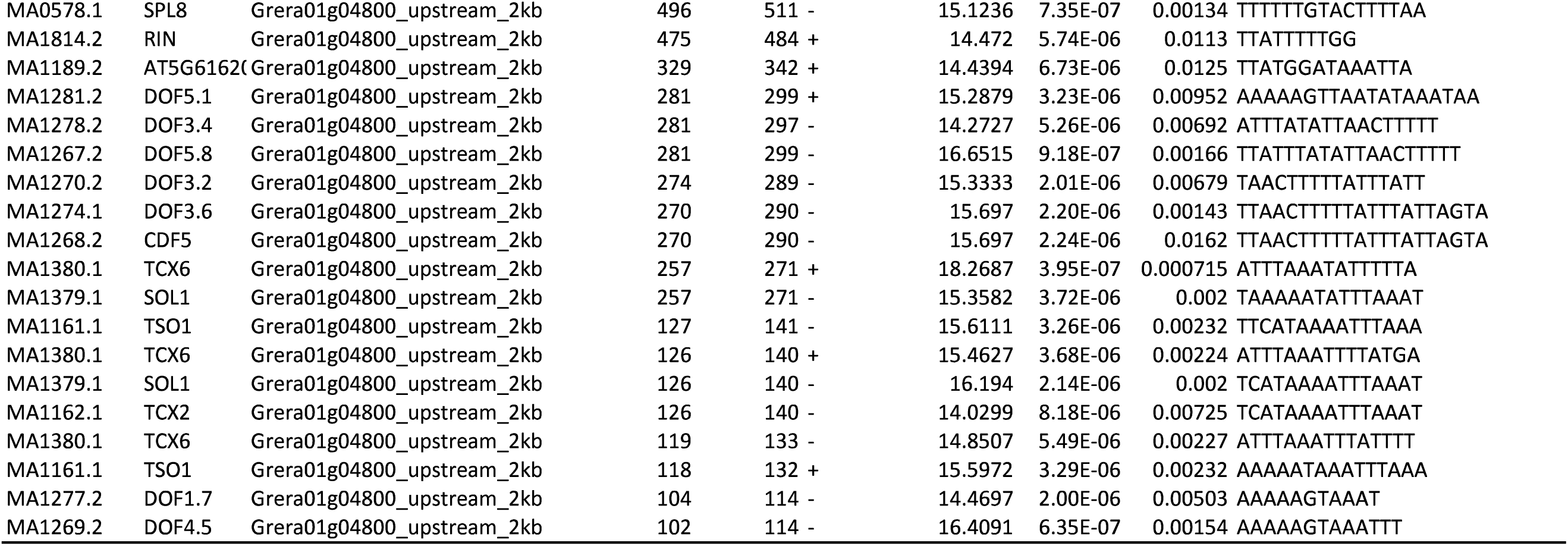
Predicted TFBSs across the 1.77-kb upstream promoter region of *LFY* in *G. radlkoferi* {A of ATG located at position 1772). The AGL6 binding site is highlighted in green.

**Table S9.**
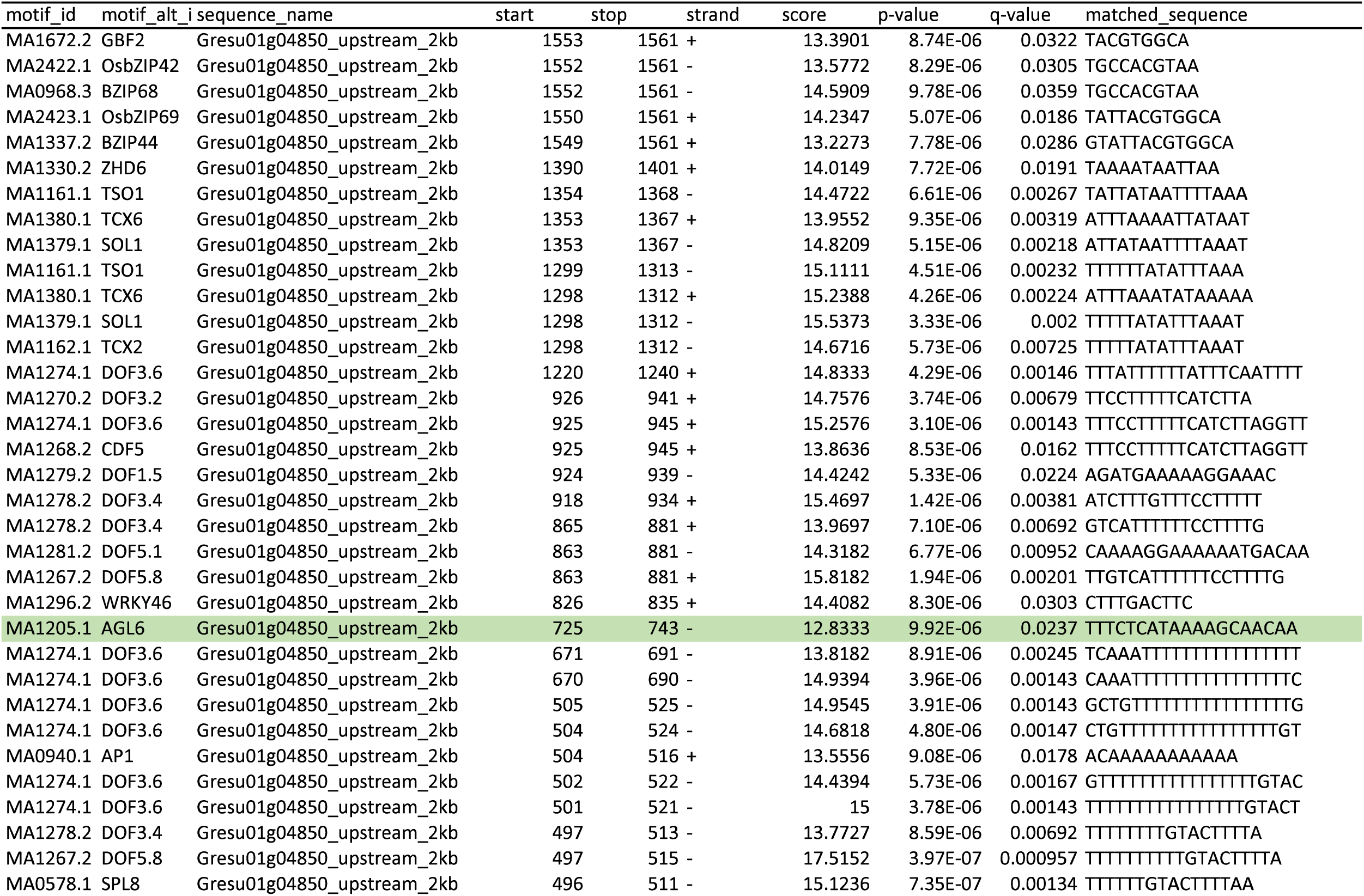
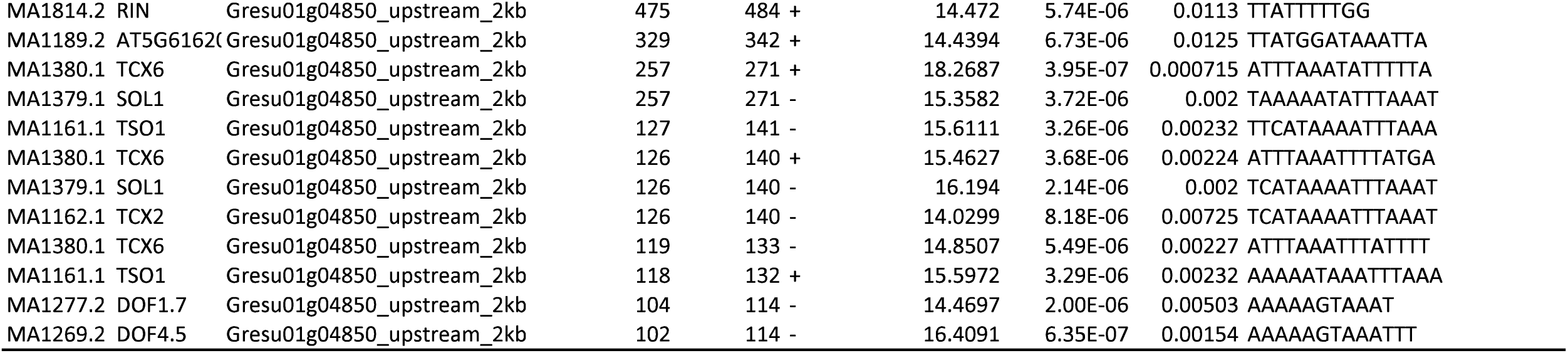
Predicted TFBSs across the 1.77-kb upstream promoter region of *LFYin G. sutherlandii* (A of ATG located at position 1769). The AGL6 binding site is highlighted in green.

**Table S10.**
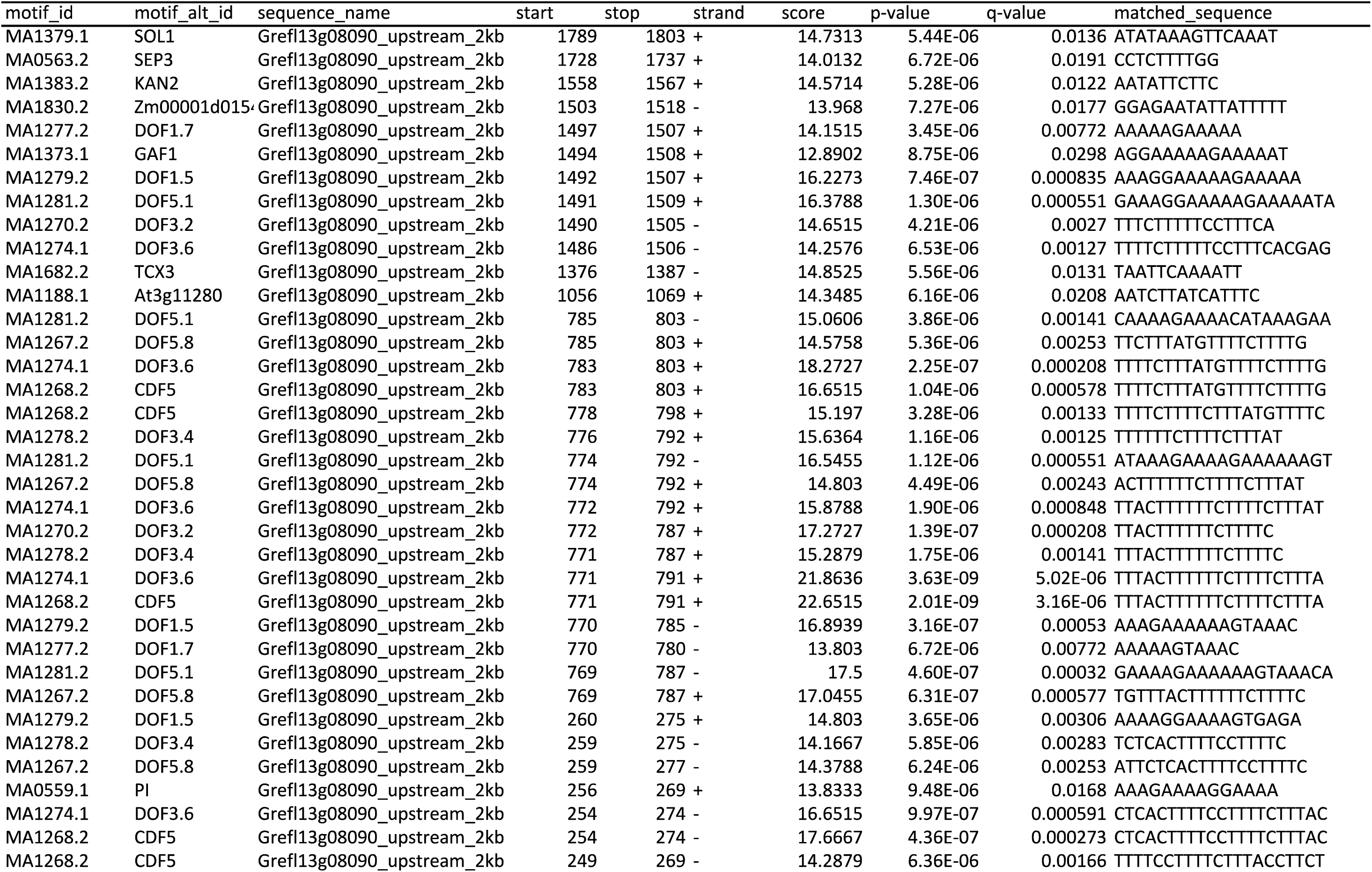
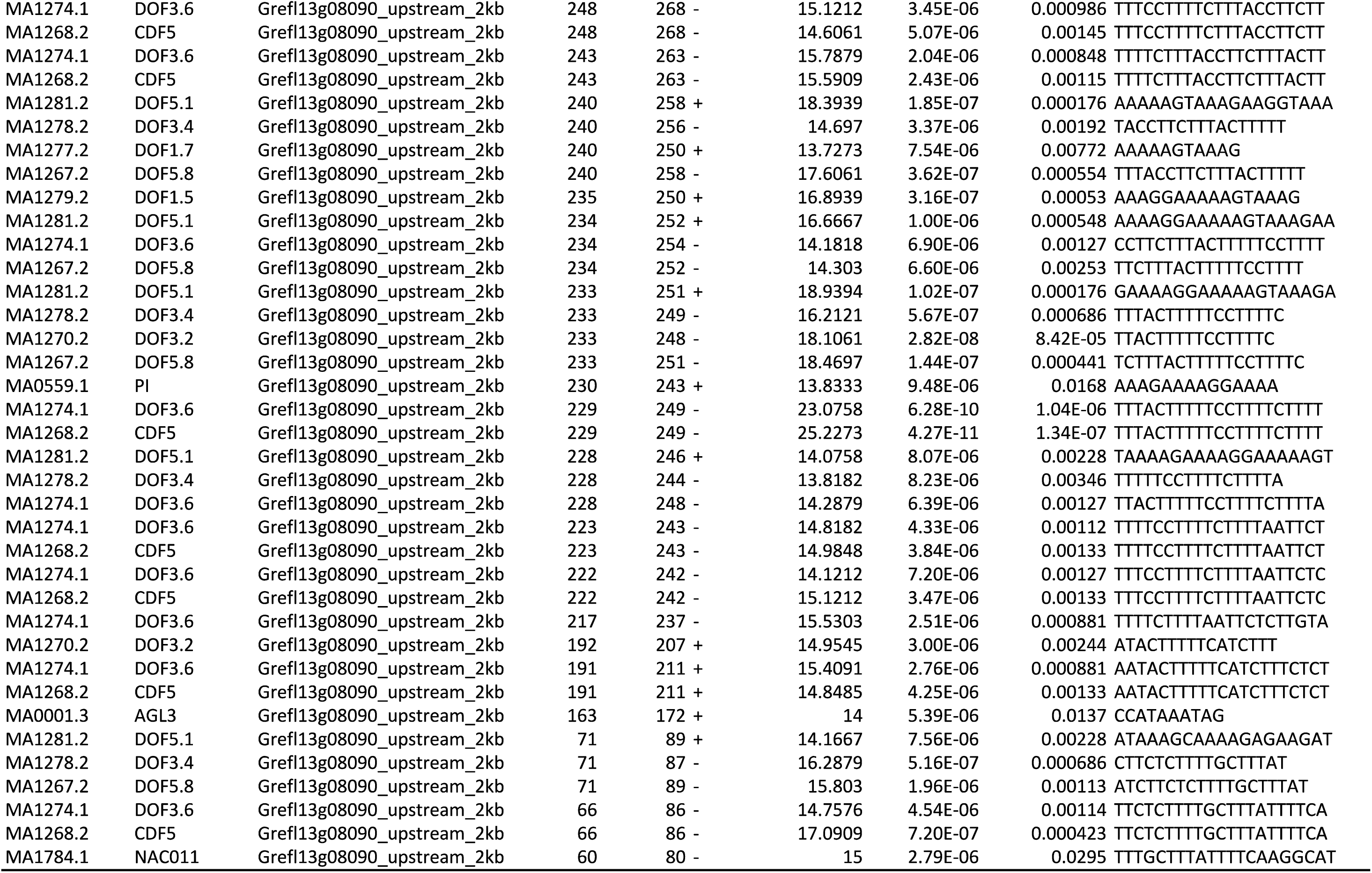
Predicted TFBSs across the 2-kb upstream promoter region of *FT* in *G. flanaganii* (A of ATG located at position 2000).

**Table S11.**
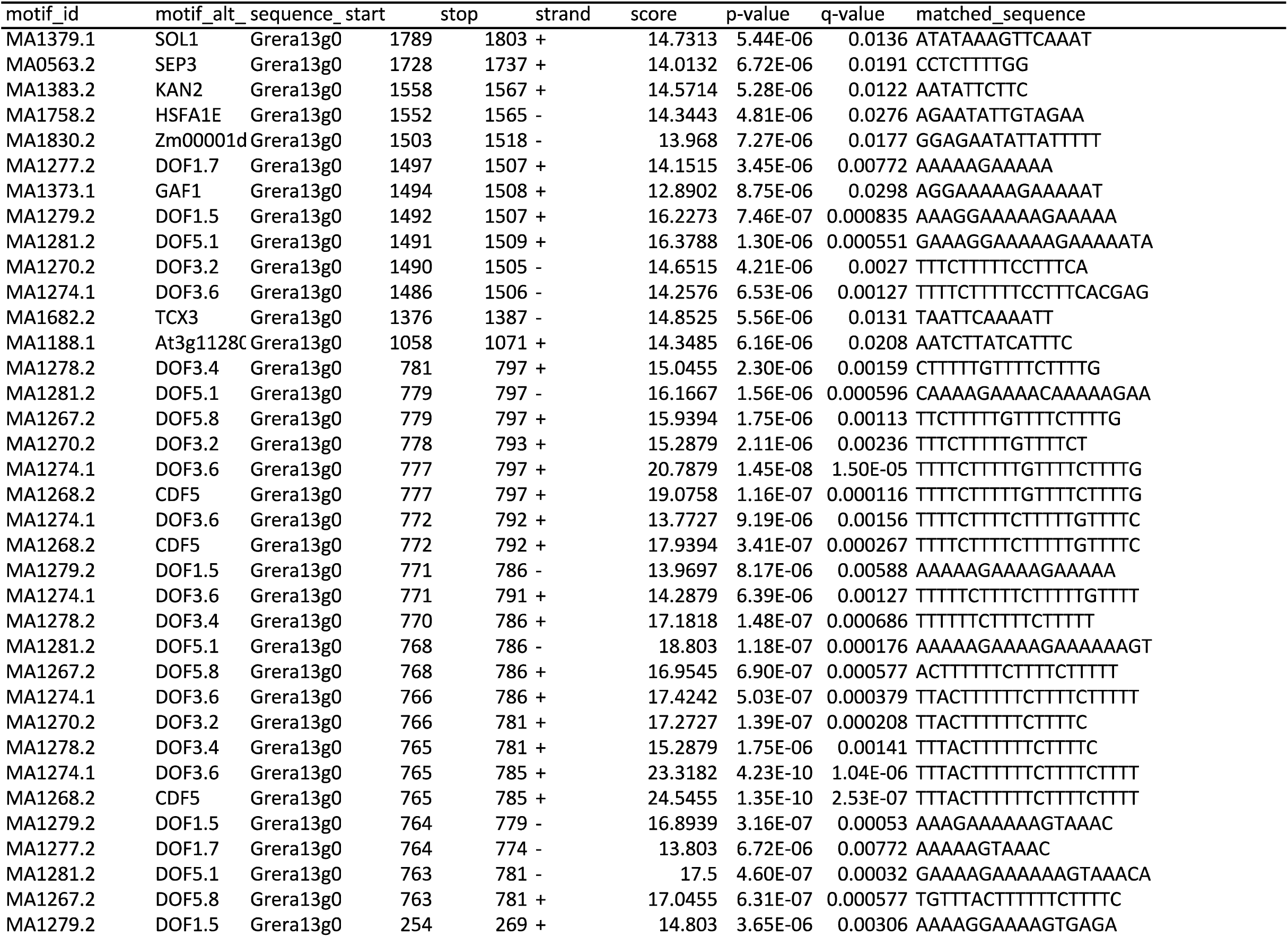
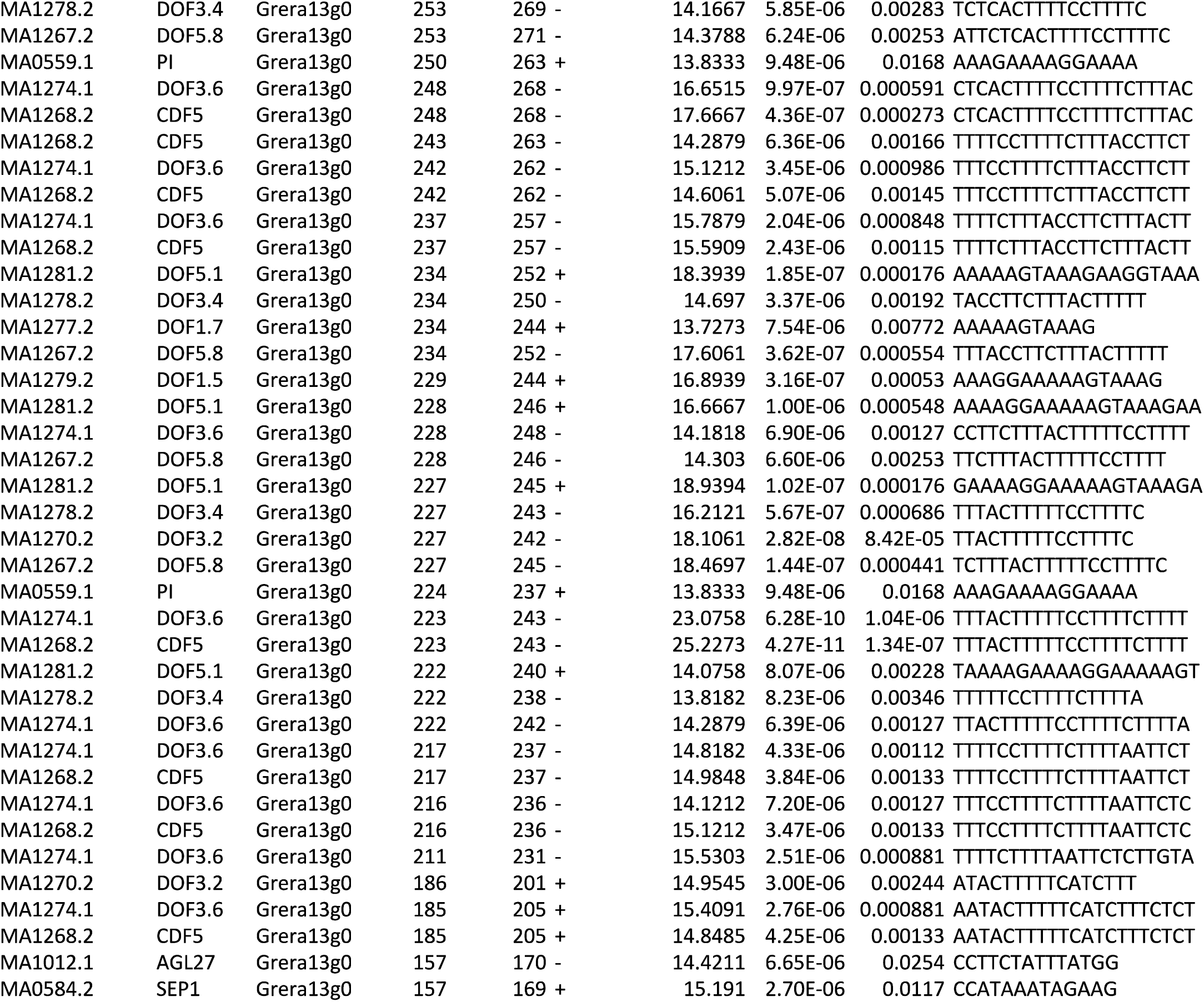
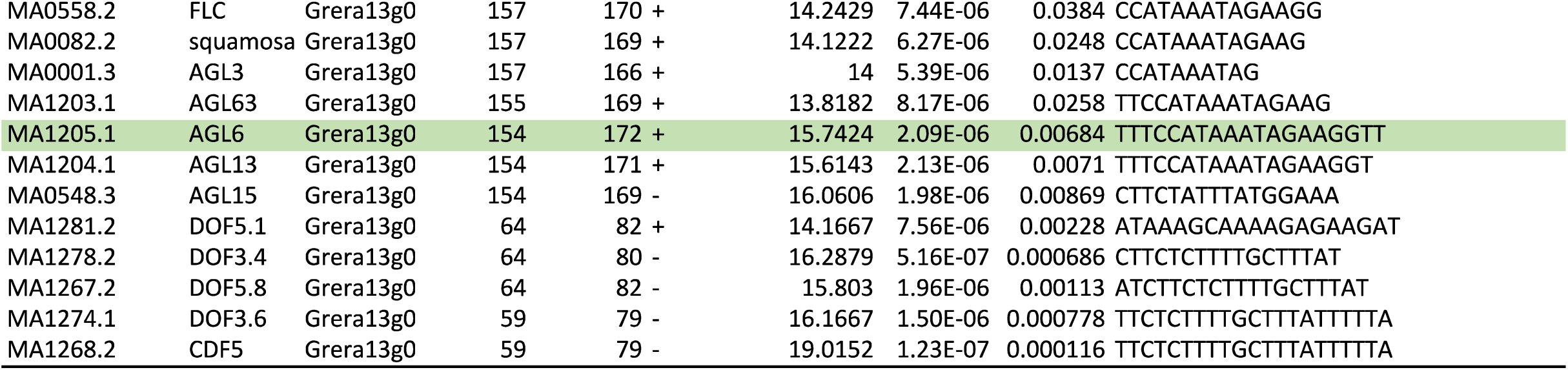
Predicted TFBSs across the 2-kb upstream promoter region of *FT* in *G. radlkoferi* (A of ATG located at position 2000). The AGL6 binding site is highlighted in green.

**Table S12.**
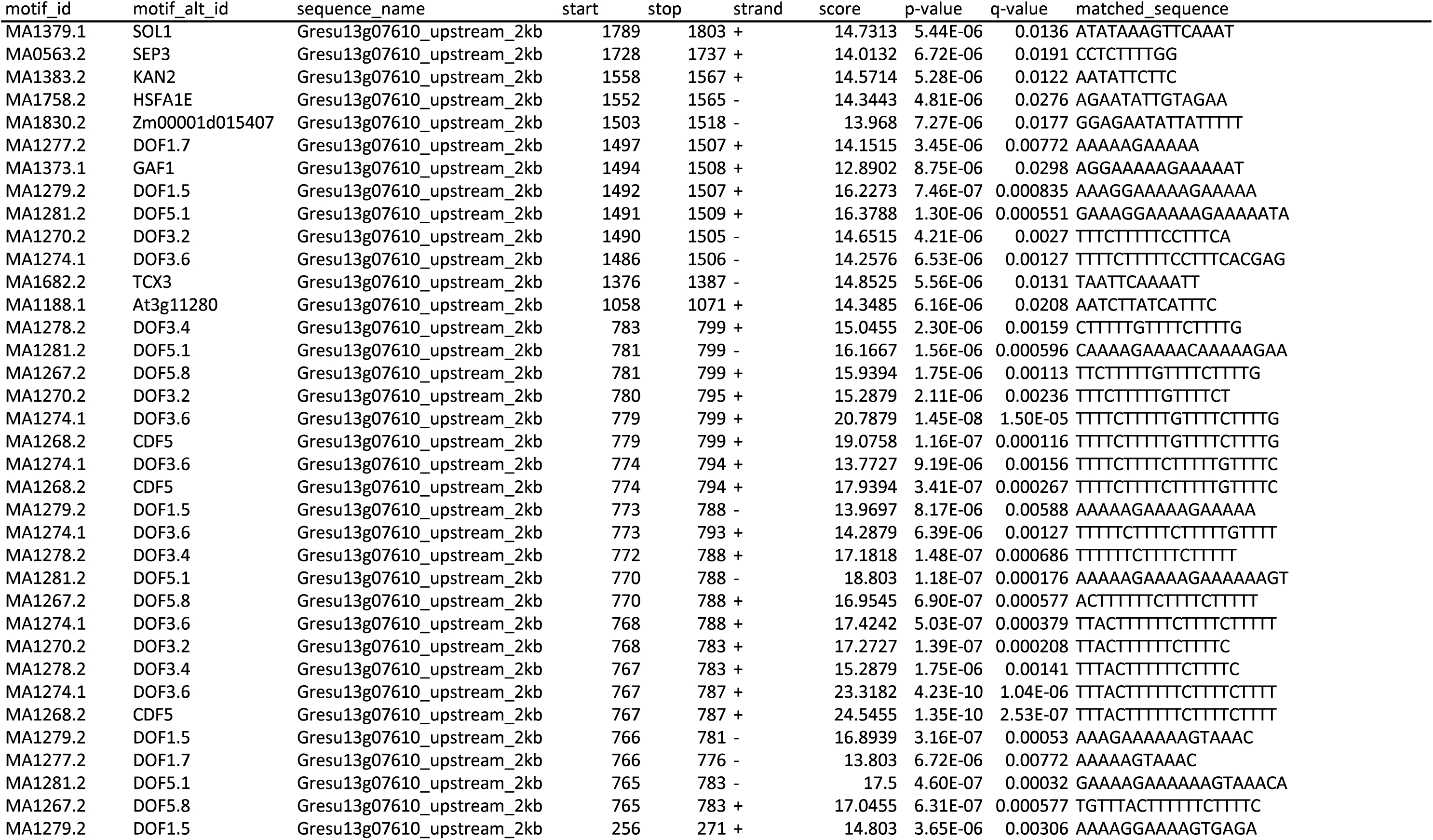
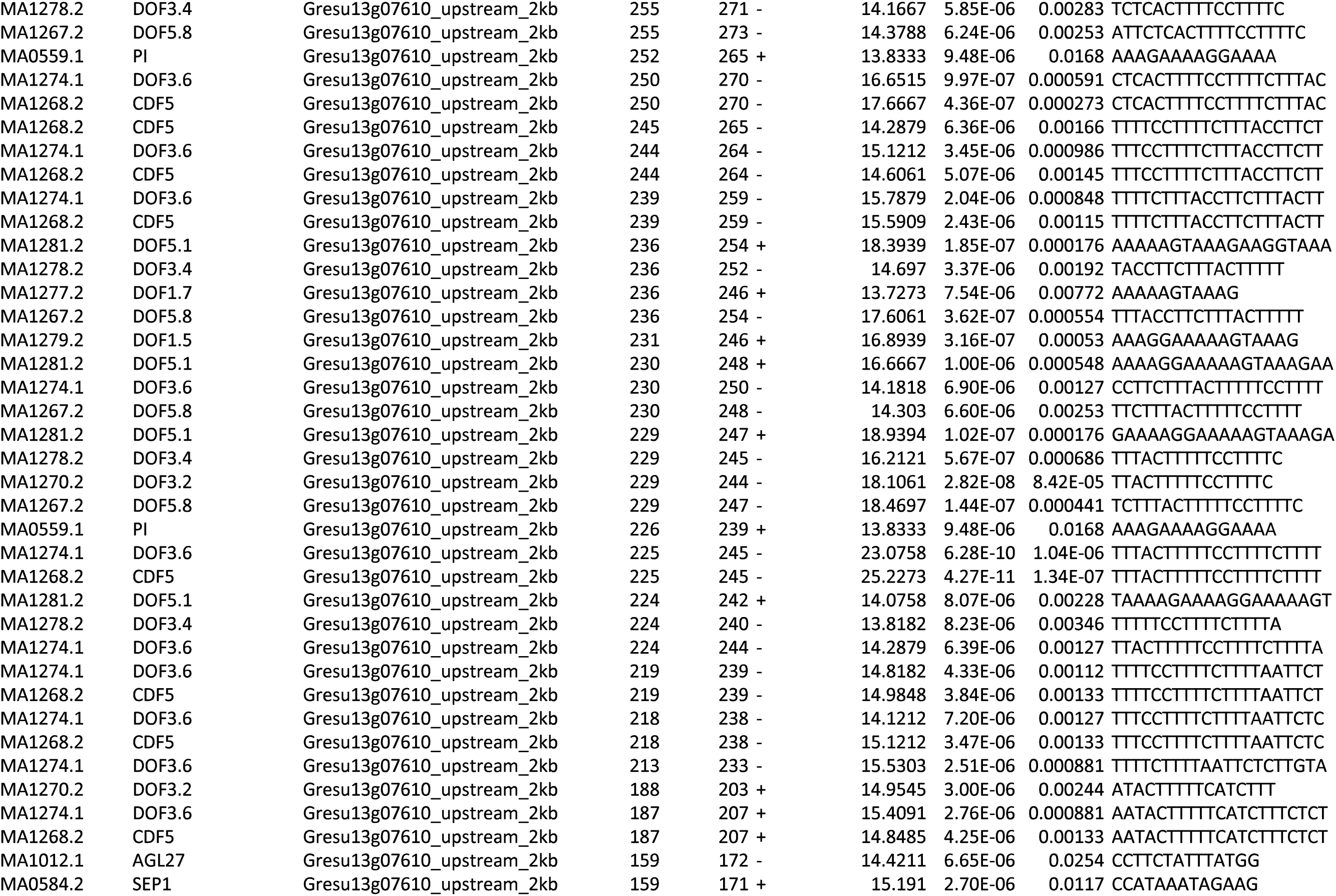
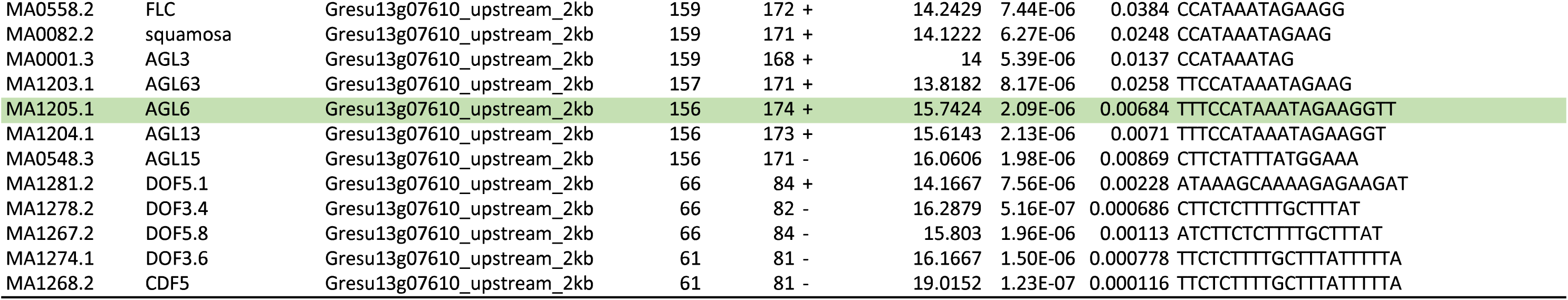
Predicted TFBSs across the 2-kb upstream promoter region of *FT* in *G. sutherlandii* (A of ATG located at position 2000). The AGL6 binding site is highlighted in green.

